# Collateral impacts of pandemic COVID-19 drive the nosocomial spread of antibiotic resistance

**DOI:** 10.1101/2022.08.15.503946

**Authors:** David R. M. Smith, George Shirreff, Laura Temime, Lulla Opatowski

## Abstract

Circulation of multidrug-resistant bacteria (MRB) in healthcare facilities is a major public health problem. These settings have been greatly impacted by the COVID-19 pandemic, notably due to surges in COVID-19 caseloads and the implementation of infection control measures. Yet collateral impacts of pandemic COVID-19 on MRB epidemiology remain poorly understood. Here, we present a dynamic transmission model in which SARS-CoV-2 and MRB co-circulate among patients and staff in a hospital population in an early pandemic context. Responses to SARS-CoV-2 outbreaks are captured mechanistically, reflecting impacts on factors relevant for MRB transmission, including contact behaviour, hand hygiene compliance, antibiotic prescribing and population structure. In a first set of simulations, broad parameter ranges are accounted for, representative of diverse bacterial species and hospital settings. On average, COVID-19 control measures coincide with MRB prevention, including fewer incident cases and fewer cumulative person-days of patient MRB colonization. However, surges in COVID-19 caseloads favour MRB transmission and lead to increased rates of antibiotic resistance, especially in the absence of concomitant control measures. In a second set of simulations, methicillin-resistant *Staphylococcus aureus* and extended-spectrum beta-lactamase-producing *Escherichia coli* are simulated in specific hospital wards and pandemic response scenarios. Antibiotic resistance dynamics are highly context-specific in these cases, and SARS-CoV-2 outbreaks significantly impact bacterial epidemiology only in facilities with high underlying risk of bacterial transmission. Crucially, antibiotic resistance burden is reduced in facilities with timelier, more effective implementation of COVID-19 control measures. This highlights the control of antibiotic resistance as an important collateral benefit of robust pandemic preparedness.

**Significance Statement:** Impacts of COVID-19 on the spread of antibiotic resistance are poorly understood. Here, an epidemiological model accounting for the simultaneous spread of SARS-CoV-2 and antibiotic-resistant bacteria is presented. The model is tailored to healthcare settings during the first wave of the COVID-19 pandemic, and accounts for hand hygiene, inter-individual contact behaviour, and other factors relevant for pathogen spread. Simulations demonstrate that public health policies enacted to slow the spread of COVID-19 also tend to limit bacterial transmission. However, surges in COVID-19 cases simultaneously select for higher rates of antibiotic resistance. Selection for resistance is thus mitigated by prompt implementation of effective COVID-19 prevention policies. This highlights the control of antibiotic resistance as an important collateral benefit of pandemic preparedness.

## Introduction

The COVID-19 pandemic has impacted the epidemiology of diverse infectious diseases, including sexually transmitted infections (e.g. HIV),(1) vector-borne illnesses (e.g. dengue virus),(2) and invasive bacterial diseases (e.g. *Streptococcus pneumoniae*).(3) Antibiotic resistance is a leading global driver of infectious morbidity and mortality,(4) yet impacts of the pandemic on the transmission and control of antibiotic-resistant bacteria remain poorly understood. There are various ways by which COVID-19 may be expected to influence the epidemiological dynamics of antibiotic resistance, particularly in healthcare settings, which face a disproportionately large share of the epidemiological burden of both antibiotic-resistant bacteria and COVID-19. On one hand, surges in COVID-19 cases may lead to conditions favourable for the proliferation of antibiotic-resistant bacteria, including hospital disorganization, increased demand on healthcare workers, abandonment of antimicrobial stewardship programmes, and administration of antibiotic prophylaxis to COVID-19 patients. On the other, public health interventions implemented to control SARS-CoV-2 transmission – including social distancing, hand hygiene education and provisioning of alcohol-based hand rub – may provide the unintended benefit of preventing bacterial transmission.

Early in the pandemic, researchers and public health officials warned that COVID-19 may impact global efforts to curb antibiotic resistance.(5, 6) However, epidemiological surveillance has been greatly challenged by the pandemic,(7) and studies to date report heterogeneous impacts of COVID-19 on antibiotic-resistant bacteria. One review highlights decreased incidence of healthcare-associated infections caused by vancomycin-resistant Enterococci (VRE) and methicillin-resistant *Staphylococcus aureus* (MRSA) relative to pre-pandemic levels.(8) Yet in an analysis of microbiological data from 81 hospitals in the USA, infections due to MRSA, VRE and multidrug-resistant Gram-negative bacteria all spiked during local surges in COVID-19 cases.(9) In a UK hospital network, bloodstream infection (BSI) due to MRSA and coagulase-negative staphylococci also spiked during surges in COVID-19 cases, while those due to Enterobacterales reached historic lows in two hospitals.(10, 11) These conflicting reports suggest that impacts of COVID-19 on antibiotic resistance likely depend on the particular population, setting and bacteria in question, and may be highly context-specific.

To anticipate and mitigate collateral impacts of SARS-CoV-2 outbreaks – and potential outbreaks of other, as-yet unknown pathogens – there is a need to better understand how the COVID-19 pandemic has both selected for and controlled against antibiotic resistance. Here, we propose a mathematical model describing simultaneous transmission of SARS-CoV-2 and commensal bacteria among patients and staff in a healthcare setting. We include mechanistic impacts of SARS-CoV-2 outbreaks on antibiotic consumption, inter-individual contact behaviour, infection prevention and control (IPC) practices, and the size and make-up of the hospital population. Simulations are used to understand and quantify how outbreaks of SARS-CoV-2 may have influenced antibiotic resistance epidemiology in an early pandemic context. Findings demonstrate how common responses to COVID-19 help to control against bacterial transmission, while simultaneously selecting for antibiotic resistance.

## Results

### 1. A nosocomial transmission model for SARS-CoV-2 and antibiotic-resistant bacteria

We present a dynamic, compartmental model describing pathogen transmission among and between patients and healthcare workers (HCWs) in an inpatient setting (**Figure 1**). The model includes: (i) infection with SARS-CoV-2 among patients and HCWs; (ii) patient colonization with either antibiotic-sensitive (“sensitive”) or antibiotic-resistant (“resistant”) strains of commensal bacteria; and (iii) transient carriage of these bacteria by HCWs as they provide care, such that HCWs act as vectors for patient colonization. HCW carriage is potentially cleared through compliance with hand hygiene, while patient colonization is potentially cleared through antibiotic use. However, antibiotics also select for resistant bacteria, both at the between-host level due to preferential clearance of sensitive bacteria, and at the within-host level due to increased rates of endogenous acquisition during antibiotic exposure.(12) See **Methods** for further model description.

**Figure 1.**
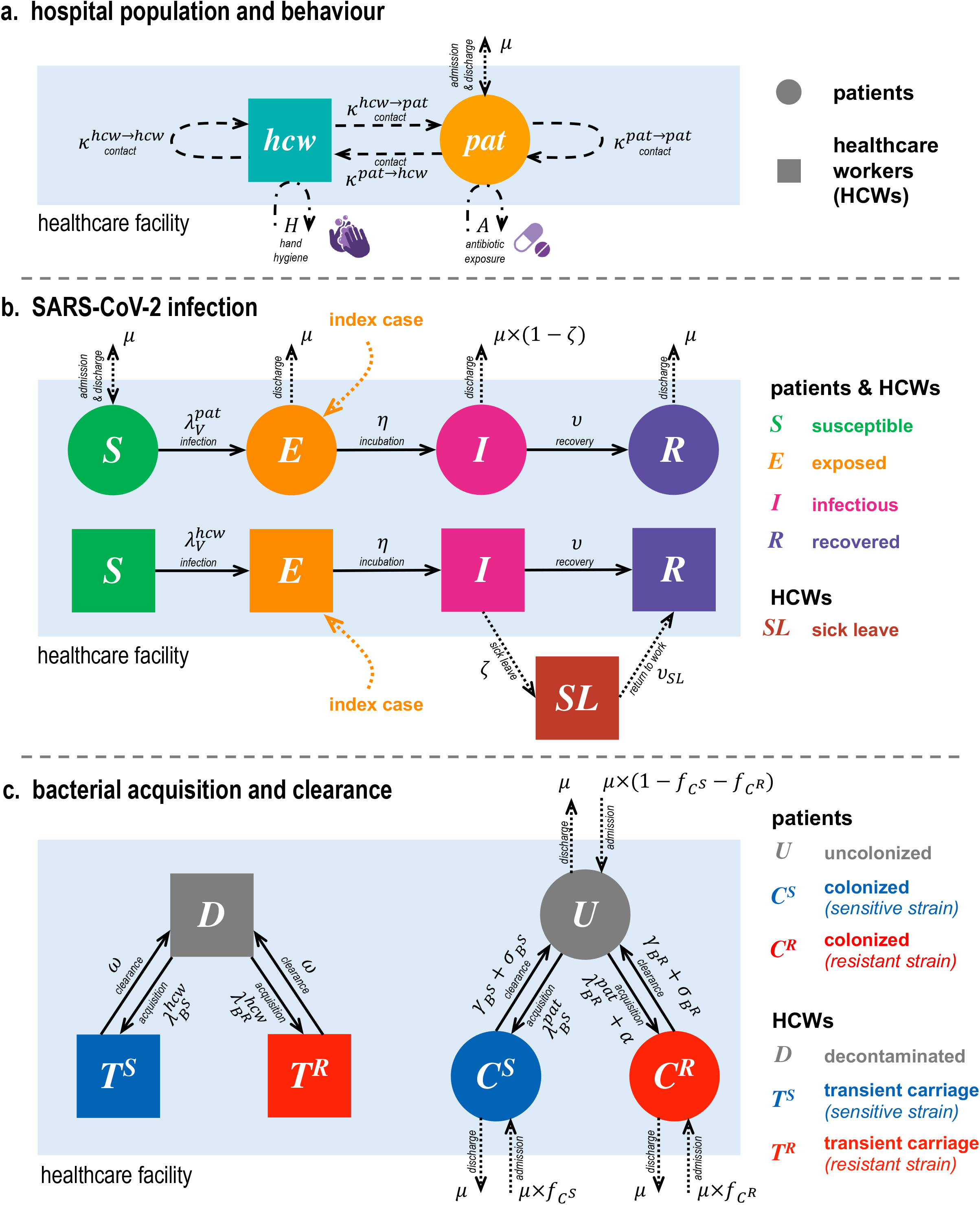
Model schematic describing the three levels of complexity included in the hospital population. (**a**) Behaviours within the healthcare facility, including asymmetric contact patterns among and between patients and HCWs, patient admission and discharge, patient exposure to antibiotics, and HCW compliance with hand hygiene. (**b**) SARS-CoV-2 infection progression among patients and HCWs, modelled as a modified Susceptible-Exposed-Infectious-Recovered (SEIR) process. (**c**) Bacterial acquisition and clearance, including patient colonization and clearance via antibiotics, and HCW carriage and clearance via hand hygiene.

Ten *COVID-19 response parameters* (τ_*i*_) are included in the model, each reflecting a distinct way in which SARS-CoV-2 outbreaks impact the organization of healthcare settings and delivery of care (detailed in **Table 1**). These parameters are normalized (0 ≤ τ ≤ 1), where τ = 0 signifies no response and τ = 1 signifies the maximum response. Each COVID-19 response is classified as either a *policy response* willingly enacted during SARS-CoV-2 outbreaks to limit viral transmission, or as a *caseload response* that unwillingly results from surges in COVID-19 patients or HCW infection (see **Methods** and **Figure S1**). The model is structured such that these parameters have mechanistic impacts on pathogen transmission (**Figure S2**).

**Table 1.** Responses to COVID-19 included in the transmission model. See Methods for description of how COVID-19 response parameters are implemented in model equations. Policy responses and caseload responses are coloured blue and black, respectively. Symptomatic refers to COVID-19 symptoms among individuals infected with SARS-CoV-2. IPC = infection prevention and control.

| COVID-19 response | Evidence | Model implementation | Category / Cause | Interpretation |  |
| --- | --- | --- | --- | --- | --- |
| | | | | $\tau = 0$ | $\tau = 1$ |
| $\tau_{as}$ Abandoned stewardship | Reduction in antibiotic stewardship activities (13) | Increased proportion of patients exposed to antibiotics ($A$) | Antibiotics / Caseload | No change in antibiotic use | Large increase in antibiotic use during COVID-19 surges |
| $\tau_{cp}$ COVID-19 prophylaxis | COVID-19 patients receive antibiotics to prevent bacterial co-infection (14) | Increased proportion of symptomatic COVID-19 patients exposed to antibiotics ($A_i$) | Antibiotics / Policy | No symptomatic patients receive antibiotic prophylaxis | All symptomatic patients receive antibiotic prophylaxis |
| $\tau_{cd}$ Cohorting disorganization | HCWs forced to provide additional care beyond their usual patients (15) | Increased daily rate of patient-HCW contact ($\kappa^{pat \rightarrow hcw}$) | Contact / Caseload | No change in contact behaviour | Large increase in daily patient-HCW contacts during COVID-19 surges |
| $\tau_{pl}$ Patient lockdown | Social interactions among patients limited or forbidden (16) | Decreased daily rate of patient-patient contact ($\kappa^{pat \rightarrow pat}$) | Contact / Policy | No change in contact behaviour | Elimination of all patient-patient contact |
| $\tau_{um}$ Universal masking | HCWs and patients wear face masks to prevent transmission (17) | Decreased SARS-CoV-2 transmissibility per contact ($\pi_v$) | IPC / Policy | No change in SARS-CoV-2 transmissibility | SARS-CoV-2 rendered non-transmissible (perfect mask effectiveness) |
| $\tau_{hh}$ Hand hygiene | Increase in HCW hand-washing performance (18) | Increased hand hygiene compliance ($H$) | IPC / Policy | No change in hand hygiene compliance | Perfect hand hygiene compliance |
| $\tau_{cs}$ COVID-19 stays | COVID-19 patients remain in healthcare facility until recovered (19) | Decreased discharge rate for symptomatic COVID-19 patients ($\mu_i$) | Disease / Caseload | No impact of SARS-CoV-2 infection on patient length of stay | All patients remain in hospital while symptomatic |
| $\tau_{ss}$ Staff sick leave | HCWs with COVID-19 stay home from work (20) | A proportion of symptomatic HCWs removed from population for 7 days (until recovered) | Disease / Caseload | No symptomatic staff go on sick leave | Symptomatic staff go on sick leave after being infectious for 1 day |
| $\tau_{ra}$ Reduced admission | Decreased number of hospital admissions during COVID-19 surges (21) | Decreased patient admission rate ($\mu$) | Admission / Caseload | No change in patient admissions | Large reduction in patient admissions during COVID-19 surges |
| $\tau_{sc}$ Sicker casemix | Elective admissions delayed or cancelled during COVID-19 surges, restricting admissions to more critically ill patients (10) | Increased rate of bacterial carriage among patient admissions ($f_{c^R}$) | Admission / Caseload | No change in the probability of colonization upon admission | Large increase in the probability of colonization with resistant bacteria upon admission during COVID-19 surges |

Using this model, simulations are conducted to quantify impacts of SARS-CoV-2 outbreaks and corresponding COVID-19 responses on the epidemiological dynamics of antibiotic resistance. When all COVID-19 responses are combined with random magnitude, nosocomial SARS-CoV-2 outbreaks have complex impacts on hospital demography and healthcare-associated behaviours (**Figure 2, panels a-e**), with heterogeneous consequences for epidemiological dynamics of SARS-CoV-2 and antibiotic resistance (**Figure 2, panels f-i**). To account for extensive parameter uncertainty reflecting heterogeneity across different bacteria and healthcare facilities, two distinct sets of probabilistic Monte Carlo simulations are conducted and presented below.

**Figure 2.**
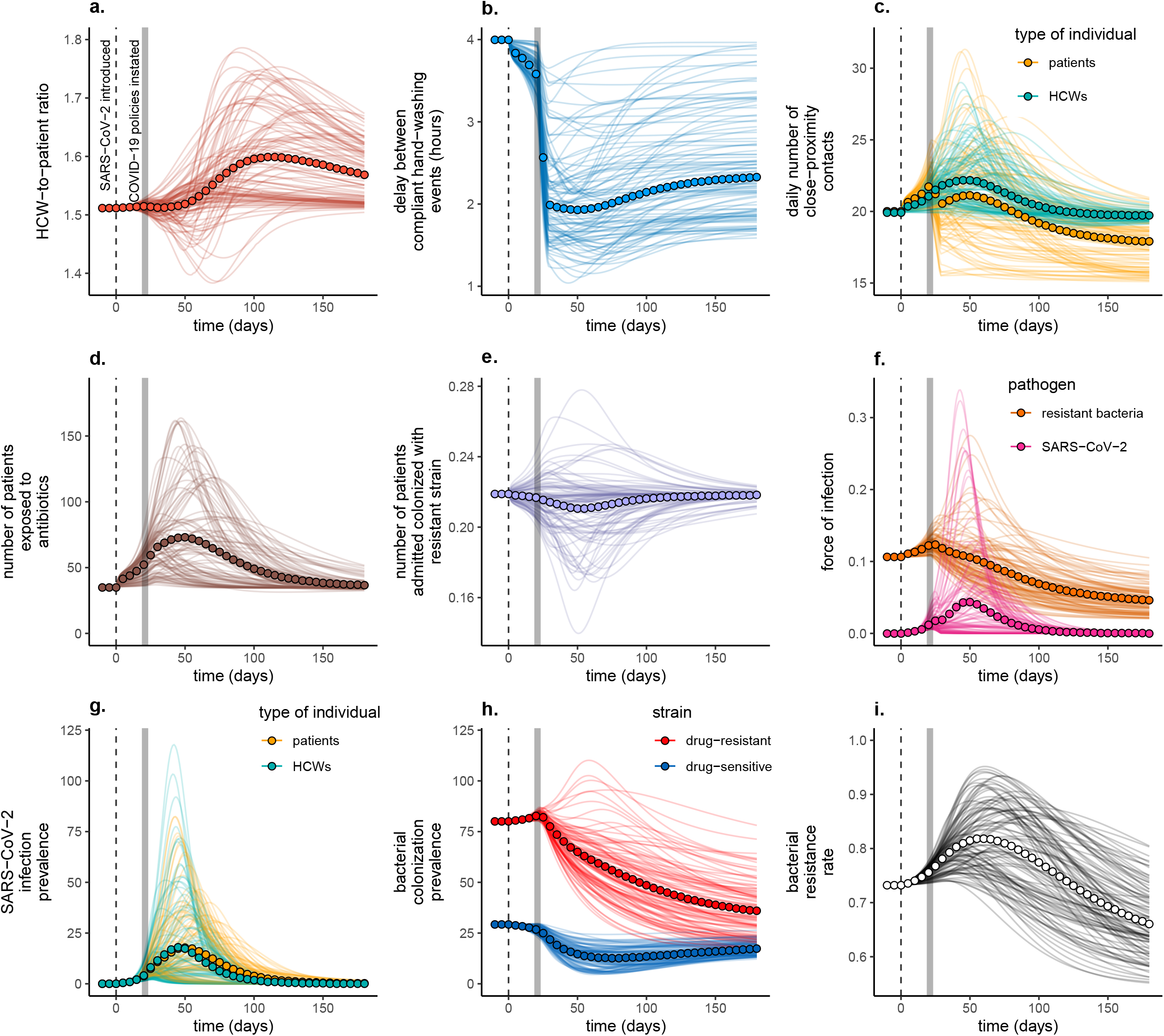
Combined influence of all COVID-19 responses on epidemiological dynamics of SARS-CoV-2 and a generic commensal bacterium in a simulated long-term care hospital with 350 beds. Baseline conditions include a 1.51:1 HCW:patient ratio, 10% antibiotic exposure prevalence (*A*_*base*_ = 0.1), 40% HCW compliance with hand hygiene after HCW-patient contact (*H*_*base*_ = 0.4), and an average 80 day patient length of stay (*µ* = 0.0125). Two index SARS-CoV-2 infections are introduced at *t* = 0 (vertical dashed lines), and policy responses are implemened at *t*_*policy*_ = 21 days (vertical grey bars) with an intervention burn-in period of *t*_*impl*_ =7 days. Lines represent dynamics across *n* = 100 independent simulations, in which all model parameters are fixed except for COVID-19 response parameters (τ), which are drawn randomly (τ∼𝒰[0,1]) for each τ in each simulation (see **Tables S1, S2**). Circles represent means across all simulations. (**a**) The ratio of HCWs to patients in the hospital (*N*^*hcw*^⁄*N*^*pat*^). (**b**) The average delay between compliant HCW handwashing events (ω/day^-1^ x 24 hours/day). (**c**) The average number of contacts that patients have with patients and HCWs (*K*^*pat*→*pat*^ + *K*^*pat*→*hcw*^, gold), and that HCWs have with patients and HCWs (*K*^*hcw*→*hcw*^ + *K*^*hcw*→*pat*^, green). (**d**) The average number of patients exposed to antibiotics 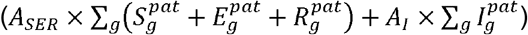. (**e**) The average number of patients admitted already colonized with resistant bacteria 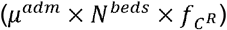. (**f**) Forces of infection for SARS-CoV-2 (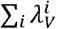, magenta) and antibiotic-resistant bacteria (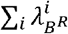, orange). (**g**) The number of active SARS-CoV-2 infections among patients (*V*^*pat*^, gold) and HCWs (*V*^*hcw*^, green). (**h**) The number of patients colonized with antibiotic-sensitive bacteria (*C*^*S*^, blue) and antibiotic-resistant bacteria (*C*^*R*^, red). (**i**) The resistance rate, the proportion of colonized patients bearing antibiotic-resistant bacteria [*C*^*R*^⁄(*C*^*S*^ + *C*^*R*^)].

### 2. Impacts of COVID-19 responses on generic MRB in generic hospitals

The first simulation set accounts for broad parameter ranges, representing “generic multidrug-resistant bacteria” (MRB) across “generic hospitals” in the context of COVID-19 responses of intermediate magnitude (τ = 0.5) (see parameter distributions in **Table S2**). In the absence of COVID-19, these hospitals and MRB are characterized by substantial epidemiological heterogeneity (**Figures S8, S9**).

#### 2.1. Combined COVID-19 responses prevent MRB colonization, but favour resistance

When all COVID-19 responses are combined (τ = 0.5), nosocomial SARS-CoV-2 outbreaks have varied impacts on healthcare-associated behaviours, hospital demography and, consequently, the epidemiological burden of MRB (**Figure 3**). Collateral impacts favouring MRB colonization include increased rates of HCW contact and increased patient exposure to antibiotics. Collateral impacts preventing MRB colonization include reduced rates of patient contact, increased rates of HCW hand decontamination and an increased HCW:patient ratio.

**Figure 3.**
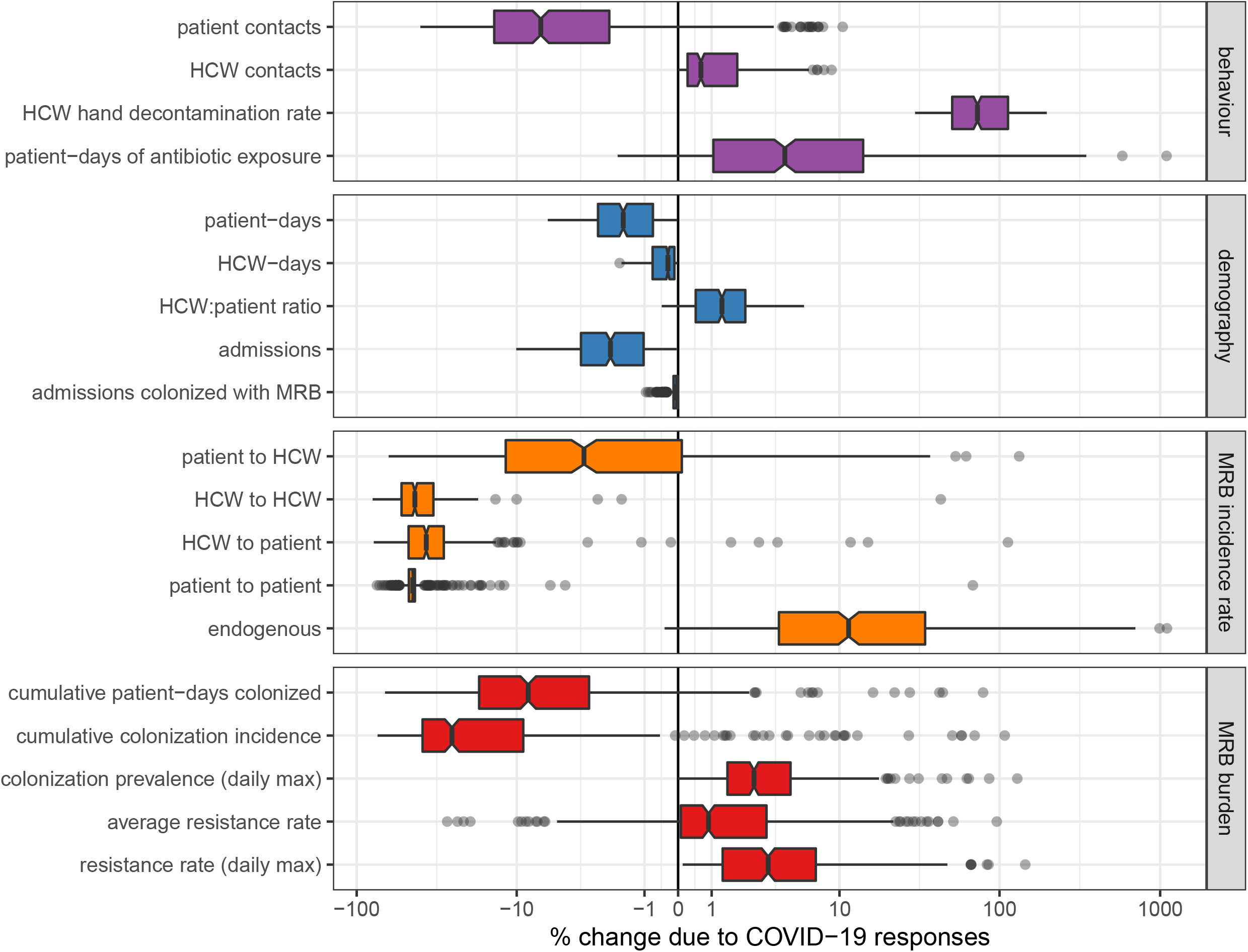
Combined COVID-19 responses impact the behaviour and demography of generic hospital populations, and consequently impact the epidemiological dynamics of generic multidrug-resistant bacteria (MRB). Each data point represents one of *n* = 500 unique MRB-hospital pairs. For each indicator (row), change due to COVID-19 responses is calculated as the difference between parameter-matched simulations including all COVID-19 responses simultaneously (*τ* = 0.5) versus those including no COVID-19 responses (*τ* = 0). Indicators are calculated cumulatively over *t* = 180 days of simulation, after introduction of two index cases of SARS-CoV-2 into the hospital at *t* = 0. Boxplots represent the *IQR*; whiskers extend to the furthest value up to ±1.5 × *IQR*; and notches extend 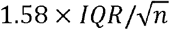, giving an approximate 95% confidence interval for comparing medians. Scales are pseudo-log_10_-transformed using an inverse hyperbolic sine function (R package ggallin).

Combined COVID-19 responses result in a mean 88.1% (58.8%–99.7%) reduction in cumulative nosocomial SARS-CoV-2 infection incidence, including reductions across all acquisition routes (**Figure S10**), but have more heterogeneous impacts on MRB epidemiology (**Figure 3**). COVID-19 responses lead to a mean 11.6% (−2.7%–44.1%) reduction in the cumulative number of patient-days colonized with MRB and a mean 24.2% (−7.8%–59.3%) reduction in the cumulative incidence of nosocomial colonization. However, incidence rates decrease for some acquisition routes (e.g. HCW-to-patient transmission) but increase for others (e.g. endogenous acquisition). COVID-19 responses also lead to transient increases in patient MRB colonization, with a mean 4.7% (0.4%–18.7%) increase in peak colonization prevalence, as well as a 2.9% (−5.4%–23.2%) increase in the average resistance rate (the cumulative share of colonized patient-days caused by resistant bacteria).

#### 2.2. Distinct COVID-19 responses have distinct epidemiological impacts

Individual COVID-19 responses have distinct impacts on MRB epidemiology (**Figure 4a**). Several COVID-19 responses are responsible for reducing the cumulative number of patient-days of MRB colonization, including reduced patient admission (τ_*ra*_), improved HCW hand hygiene (τ_*hh*_) and patient lockdown (τ_*pl*_). However, no COVID-19 responses lead to meaningful reductions in the average resistance rate, although hand hygiene and patient lockdown are associated with high variance in this outcome despite negligible change on average. The COVID-19 response that most favours increasing resistance is abandoned stewardship (τ_*as*_), which in the absence of other COVID-19 responses causes a mean 9.4% (0.1%–49.0%) increase in the average resistance rate. This is followed by sicker casemix (τ_*sc*_) with a 2.5% (<0.1%–7.7%) increase in the average resistance rate, and COVID-19 prophylaxis (τ_*cp*_) with a 1.0% (<0.1%–6.5%) increase.

**Figure 4.**
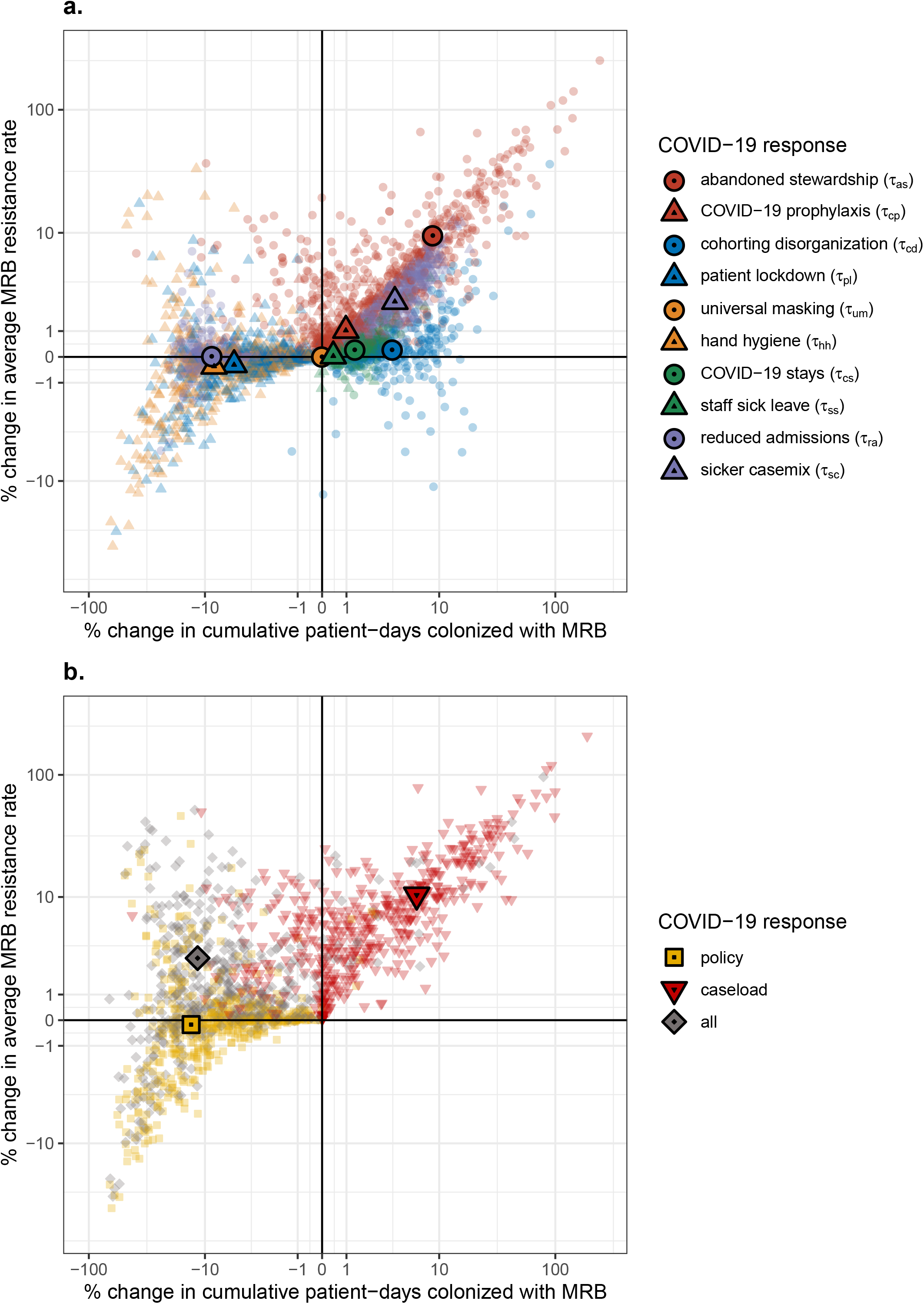
Some COVID-19 responses prevent, while others favour MRB colonization. Scatter plots depict change in the cumulative number of patient-days of MRB colonization (x-axis) and the average MRB resistance rate (y-axis) resulting from: (**a**) individual COVID-19 responses, each given by a unique colour-shape combination, and (**b**) combinations of COVID-19 responses. Policy responses include COVID-19 prophylaxis (*τ*_*cp*_), patient lockdown (*τ*_*pl*_), universal masking (*τ*_*um*_) and hand hygiene (*τ*_*hh*_). Caseload responses include abandoned stewardship (*τ*_*ss*_), cohorting disorganization (*τ*_*cd*_), COVID-19 stays (*τ*_*cs*_), staff sick leave (*τ*_*ss*_), reduced admission (*τ*_*ra*_) and sicker casemix (*τ*_*sc*_). Small translucent points represents unique MRB-hospital pairs, and larger opaque points represent means across *n* = 500 pairs. For each indicator, change due to COVID-19 responses is calculated as the difference between parameter-matched simulations including respective COVID-19 responses (*τ* = 0.5) versus those including no COVID-19 responses (*τ* = 0). Indicators are calculated cumulatively over *t* = 180 days of simulation, after introduction of two index cases of SARS-CoV-2 into the hospital at *t* = 0. Scales are pseudo-log_10_-transformed using an inverse hyperbolic sine function (R package ggallin).

Individual COVID-19 responses also have distinct impacts on the incidence of SARS-CoV-2 infection and MRB colonization, with changes varying across different routes of acquisition. Reductions in SARS-CoV-2 incidence are led by reduced transmission from all individuals due to universal masking (τ_*um*_), reduced transmission from HCWs due to staff sick leave (τ_*ss*_), reduced transmission to patients due to reduced admissions (τ_*ra*_), and reduced patient-to-patient transmission due to patient lockdown (τ_*pl*_) (**Figure S11**). Reductions in MRB incidence are largely due to reduced transmission from HCWs as a result of improved hand hygiene (τ_*hh*_), reduced transmission from patients due to patient lockdown (τ_*pl*_), and reduced transmission to and from all individuals due to reduced patient admission (_*τra*_) (**Figure S12**). These COVID-19 responses outweigh the impacts of competing COVID-19 responses that favour greater SARS-CoV-2 incidence [cohorting disorganization (τ_*cd*_) and COVID-19 stays (τ_*es*_)] and/or MRB incidence [abandoned stewardship (τ_*as*_), cohorting disorganization (τ_*cd*_), sicker casemix (τ_*sc*_), COVID-19 stays (τ_*cs*_), staff sick leave (τ_*ss*_) and COVID-19 prophylaxis (τ_*cp*_)] (**Figures S13, S14**).

#### 2.3. Surges in COVID-19 caseloads lead to strong selection for resistance

Impacts of COVID-19 responses on MRB epidemiology vary across policy responses and caseload responses (**Figure 4b**). The cumulative number of patient-days colonized with MRB decreases by a mean 13.2% (0.6%–46.9%) with policy responses, but increases by a mean 6.3% (−5.2%–42.8%) with caseload responses. Mirroring these trends, MRB colonization incidence decreases by 28.2% (2.5%–60.2%) with policy responses but increases by 13.8% (−3.5%–77.0%) with caseload responses (**Figure S13**). By contrast, policy responses have little impact on the average resistance rate, while caseload responses lead to a mean 10.4% (0.2%–46.9%) increase.

### 3. Impacts of COVID-19 on MRSA and ESBL-*E. coli* across wards and scenarios

The second simulation set accounts for a series of case studies representing specific bacteria (MRSA and ESBL-*E. coli*; see **Table S3**), specific healthcare settings (a geriatric rehabilitation ward, a short-stay geriatric ward, and a general paediatric ward; see **Table S4**), and specific COVID-19 response scenarios (an organized response, an intermediate response, and an overwhelmed response; see **Table S5**).

#### 3.1. Baseline nosocomial dynamics differ across wards

Baseline healthcare-associated behaviours and demography vary across wards (**Figure S15**), resulting in ward-specific epidemiological dynamics of MRSA and ESBL-*E. coli* colonization (prior to introduction of SARS-CoV-2) (**Figure S16**). Due to different factors including ward-specific differences in antibiotic exposure, patient length of stay and inter-individual contact rates, the relative importance of different colonization acquisition routes varies across settings and bacteria. For MRSA, for instance, endogenous acquisition dominates in the short-stay ward, HCW-to-patient transmission dominates in the general ward, and patient-to-patient transmission dominates in the rehabilitation ward (**Figure S17**). Upon introduction of SARS-CoV-2, ward-specific characteristics further translate to variability in SARS-CoV-2 risk and infection dynamics (**Figure S18**). In both the short-stay ward and the general ward, most SARS-CoV-2 transmission results from HCWs, with negligible transmission from patients to HCWs or other patients. Conversely, in the rehabilitation ward, patient-to-patient transmission is the dominant acquisition route.

#### 3.2. Overwhelmed COVID-19 responses exacerbate antibiotic resistance

Impacts of SARS-CoV-2 outbreaks on antibiotic resistance epidemiology vary across wards, bacterial species and COVID-19 response scenarios (**Figure 5**). SARS-CoV-2 outbreaks have little impact on bacteria acquired predominantly via endogenous acquisition, including ESBL-*E. coli* in the general ward and both MRSA and ESBL-*E. coli* in the short-stay ward (**Figure S17**). For remaining contexts with substantial bacterial transmission to and from patients, SARS-CoV-2 outbreaks have significant impacts on bacterial colonization incidence and the bacterial resistance rate (**Figure 5**). Overwhelmed COVID-19 responses are associated with higher colonization incidence and higher resistance rates than organized responses. For instance, given an organized response in the rehabilitation ward, colonization incidence of MRSA decreases by a mean 65.2%% (95%UI: 48.8%–79.9%), with little change in the resistance rate. Conversely, given an overwhelmed response, there is little change in incidence, while the resistance rate increases by a mean 18.7% (95%UI: 8.5%–28.3%). These impacts result from how different COVID-19 response scenarios modify healthcare-associated behaviours and demography in each ward. In the rehabilitation ward, for example, patient antibiotic exposure and the daily number of contacts per patient tend to increase in the overwhelmed scenario, but decrease in the organized scenario (**Figure S19**).

**Figure 5.**
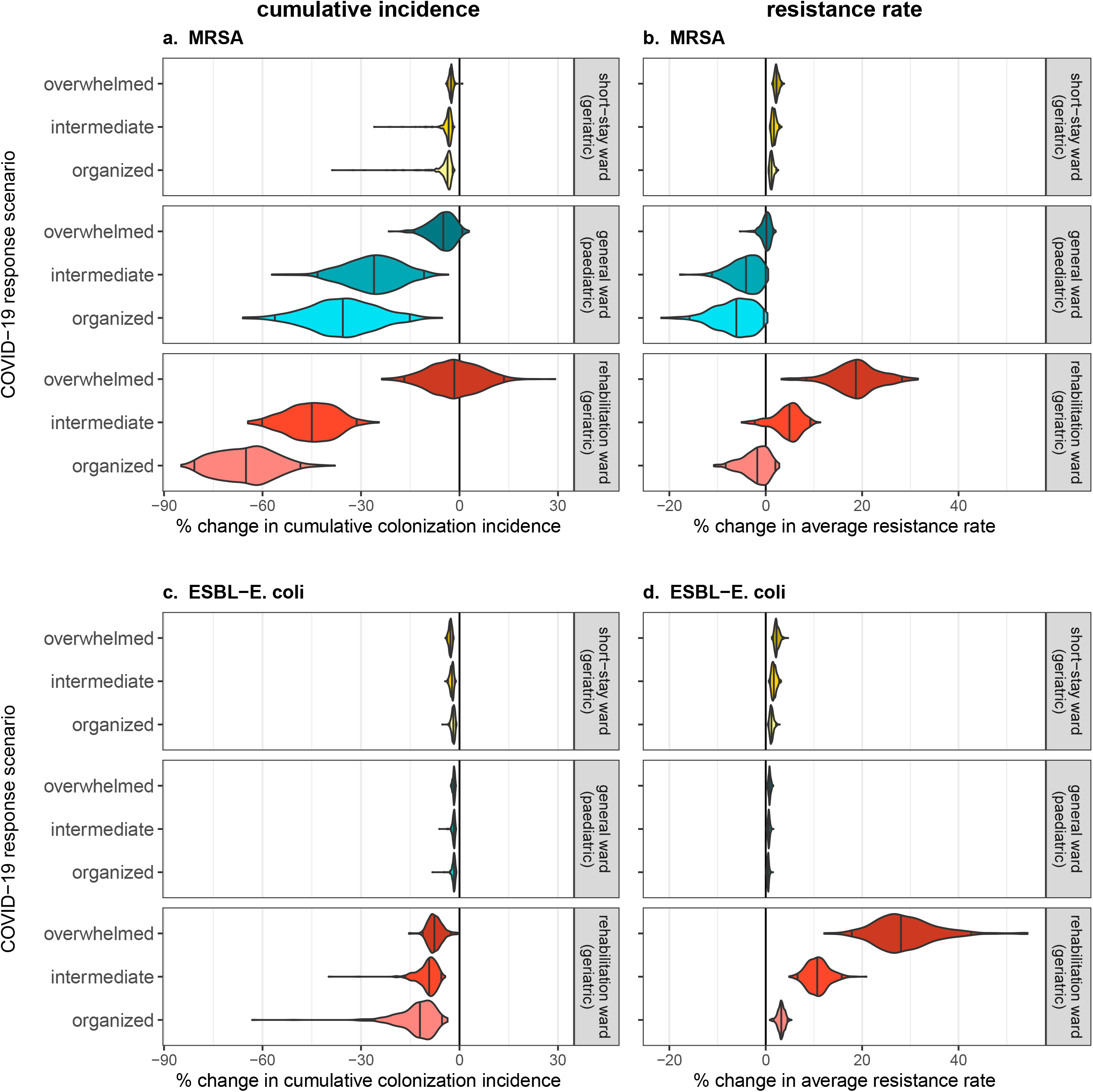
Overwhelmed responses to COVID-19 result in greater colonization burden of antibiotic-resistant bacteria relative to organized responses. Violin plots represent outcome distributions from *n* = 500 Monte Carlo simulations, and depict cumulative change in epidemiological indicators due to nosocomial SARS-CoV-2 outbreaks (x-axis) across different COVID-19 response scenarios (y-axis). Results are presented for (**a**) cumulative MRSA colonization incidence, (**b**) the average MRSA resistance rate, (**c**) cumulative ESBL-*E. coli* colonization incidence and (**d**) the average ESBL-*E. coli* resistance rate. For each hospital ward, bacterial species and COVID-19 response scenario, change due to COVID-19 responses is calculated as the difference between parameter-matched simulations including respective COVID-19 responses (organized, intermediate or overwhelmed; see **Table S5**) versus those including no COVID-19 responses (*τ* = 0), assuming baseline values of SARS-CoV-2 transmissibility (*β*_*V*_ = 1.28) and policy implementation timing (*t*_*policy*_ = 21). Indicators are calculated cumulatively over *t* = 180 days of simulation, after introduction of two index cases of SARS-CoV-2 into the hospital at *t* = 0.

Impacts of COVID-19 on antibiotic resistance also depend on the transmissibility of SARS-CoV-2 (*β*_*V*_) and the timing of COVID-19 policy implementation (*t*_*policy*_) (visualized for MRSA in the rehabilitation ward in **Figure 6**; see all bacteria and wards in **Figures S20** and **S21**). Increasing the SARS-CoV-2 transmission rate results in larger SARS-CoV-2 outbreaks, and in turn greater selection for resistance across all wards, bacteria and control scenarios. However, impacts of policy timing depend on the nature of the policies being implemented. Earlier implementation of organized responses generally results in lower resistance rates, due to their ability to help control the spread of and selection for resistant bacteria. Conversely, earlier implementation of overwhelmed responses generally results in higher resistance rates, as these responses tend to exert additional selection for resistant bacteria.

**Figure 6.**
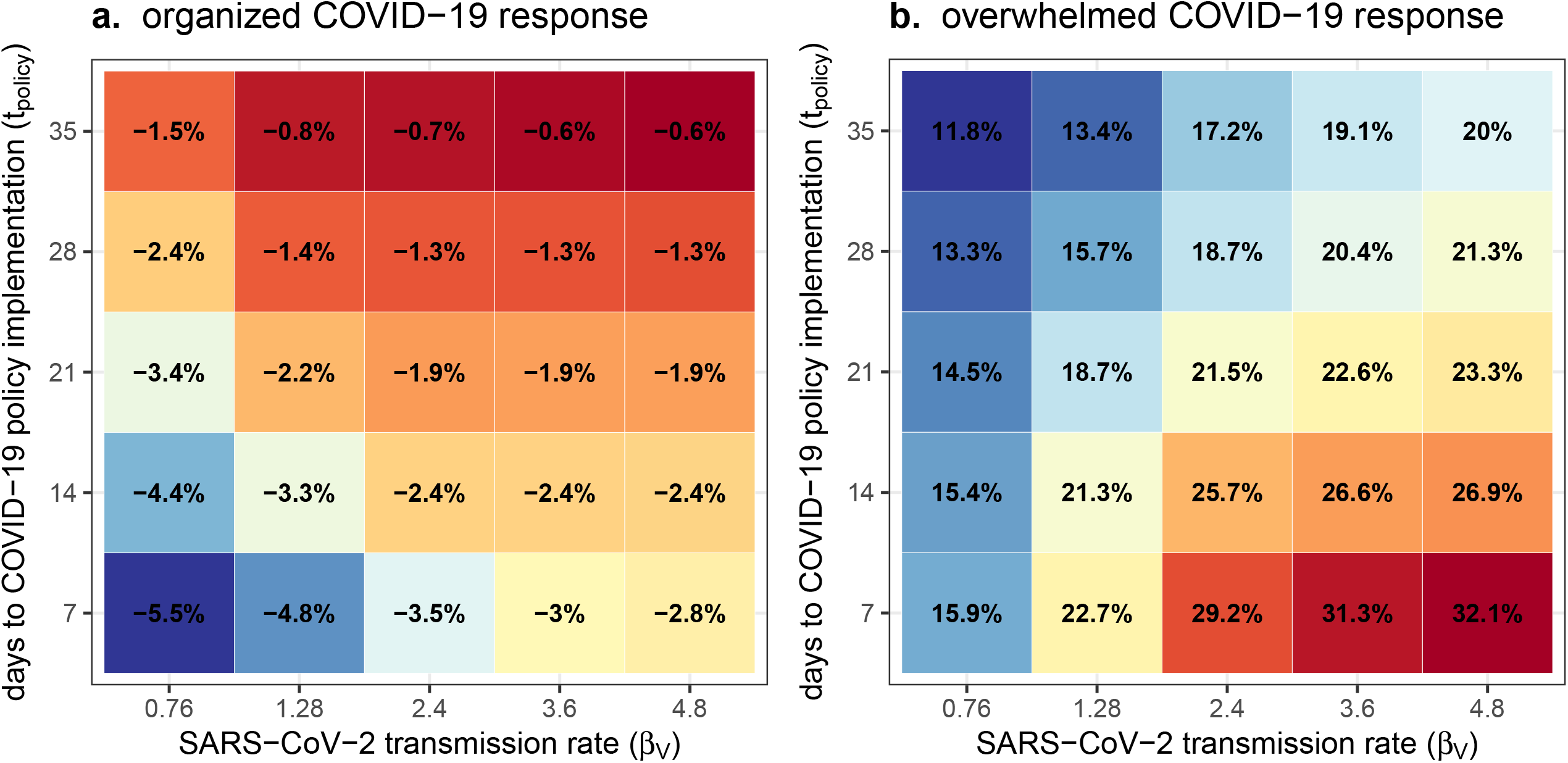
Antibiotic resistance is mitigated by earlier implementation of organized COVID-19 responses, but exacerbated by earlier implementation of overwhelmed COVID-19 responses. Each coloured tile depicts mean change in the average resistance rate of MRSA across *n* = 500 Monte Carlo simulations, which varies with the SARS-CoV-2 transmission rate (x-axis) and the delay to COVID-19 policy implementation (y-axis) in a geriatric rehabilitation ward with (**a**) an organized response to COVID-19 versus (**b**) an overwhelmed response. Change due to COVID-19 responses is calculated as the difference between parameter-matched simulations including respective COVID-19 responses (organized, overwhelmed; see **Table S5**) versus those including no COVID-19 responses (*τ* = 0). Average resistance rate is calculated cumulatively over *t* = 180 days of simulation, after introduction of two index cases of SARS-CoV-2 into the hospital at *t* = 0.

## Discussion

This study demonstrates how collateral impacts of COVID-19 both favour and prevent against the spread of antibiotic resistance in healthcare settings. Surges in COVID-19 cases – and associated consequences like abandonment of antibiotic stewardship programmes and disorganization of patient care – were found to favour the spread of resistant bacteria. Conversely, COVID-19 control policies like patient lockdown and reinforcement of IPC measures were effective for prevention of bacterial colonization. Such policies work not only by directly preventing bacterial transmission, but also by limiting surges in COVID-19 cases and the conditions favourable for bacterial spread that they create. These findings thus highlight that limiting the proliferation of antibiotic resistance is an important collateral benefit of nosocomial COVID-19 prevention. This suggests that various other public health strategies effective for prevention of SARS-CoV-2 transmission in healthcare settings – including vaccination, mass testing and HCW cohorting – may further help to alleviate the spread of antibiotic resistance.(22–24)

Findings also demonstrate that better pandemic preparedness may serve to limit unintended selection for antibiotic-resistant bacteria. Our simulations tailored to an early pandemic context found that patients in better organized healthcare facilities (i.e. those enacting more effective COVID-19 control policies sooner) were less likely to acquire bacterial colonization and experienced lower rates of resistance than patients in facilities overwhelmed by COVID-19. In the context of poorly anticipated outbreaks of any novel respiratory pathogen, healthcare facilities that rapidly instate effective IPC measures while limiting unnecessary surges in antibiotic use may thus avoid concomitant surges in antibiotic resistance. With hindsight, the rapid global spread of SARS-CoV-2 and its associated health system shocks in early 2020 revealed insufficient global capacity to detect and contain novel pathogens with pandemic potential.(25) This has spurred calls for transformational change in international law and governance, and expansive global investment in pandemic preparedness.(26) Our study suggests that mitigating the spread of antibiotic-resistant bacteria should be considered as a collateral benefit of pandemic preparedness initiatives, with implications for their funding and design.

Collateral impacts of COVID-19 have evolved over successive pandemic waves, and will continue to evolve through the transition to endemic COVID-19.(27, 28) Such variability has coincided with the ebbing and flowing of enforcement of nonpharmaceutical COVID-19 control interventions, availability of vaccines and antiviral therapies, and capacity and resilience of healthcare systems. The clinical and epidemiological characteristics of COVID-19 are also in constant flux, due to evolution of intrinsic virulence and immune escape properties of SARS-CoV-2 variants, and great heterogeneity in acquisition and waning of both natural and vaccine-induced immunity. Understanding how these complex, overlapping factors combine to influence the spread of antibiotic-resistant bacteria remains a great challenge. Interrupted epidemiological surveillance and the reallocation of public health resources away from antibiotic resistance programmes and towards COVID-19 control has made this particularly difficult. In many instances, SARS-CoV-2 testing and surveillance infrastructure has been repurposed from existing antimicrobial resistance infrastructure, leading to reduced whole genome sequencing of bacterial isolates, and extensive gaps and delays in the reporting of antimicrobial resistance data since the onset of the pandemic.(7)

In light of such complexity and data limitations, mathematical modelling should continue to be exploited as a tool to understand mechanistic links between COVID-19 and antibiotic resistance beyond the early pandemic context explored here. Future work should evaluate impacts across other types of healthcare and residential facilities (e.g. retirement homes, prisons), nosocomial bacteria (e.g. *Pseudomonas aeruginosa, Acinetobacter baumannii, Enterobacter* spp.), and specific medical procedures associated with both COVID-19 and nosocomial MRB spread (e.g. mechanical ventilation, central venous catheterization). Future work is also needed to understand impacts of COVID-19 on antibiotic resistance across entire health systems and in community settings. An international surge in antibiotic consumption was observed initially in March 2020, associated primarily with antimicrobials used as treatment for COVID-19 and/or prophylaxis for bacterial coinfection (e.g. azithromycin, hydroxychloroquine).(29, 30) Subsequently, there was an estimated 19% reduction in global antimicrobial consumption from April to August 2020 (relative to 2019).(31) In combination with reduced human mobility, contact rates and care-seeking during COVID-19 lockdowns, epidemiological impacts of modified antibiotic consumption in the community remain poorly understood. Data subsequent to first wave lockdowns have shown a decrease in ESBL resistance among *E. coli* isolates in France,(32) and reduced carriage of cephalosporin- and carbapenem-resistant Enterobacterales in Botswana.(33) However, more longitudinal estimates from diverse geographical regions and bacterial species are greatly needed.

Although few studies have reported explicitly on impacts of COVID-19 on distributions of antibiotic-resistant strains or serotypes, COVID-19 lockdowns in early 2020 clearly reduced incidence of disease due to community-associated respiratory bacteria like *S. pneumoniae, Haemophilus influenzae* and *Neisseria meningitidis*.(3, 34–37) Yet over the same time period, intriguingly, emerging data report persistent carriage of *S. pneumoniae* in the community across countries and age groups.(38–40) It has been suggested that reduced incidence of bacterial infection may thus be explained at least in part by concomitant prevention of other respiratory viruses like influenza,(41) which have been shown to favour progression from bacterial colonization to disease.(42) Fully understanding impacts of COVID-19 on any particular form of antibiotic resistance may therefore require taking into account not only SARS-CoV-2 and the bacterium in question, but also other interacting microorganisms. For simplicity, and due to limited evidence of a strong association between SARS-CoV-2 and bacterial coinfection,(14) our model assumes no impact of SARS-CoV-2 infection on bacterial acquisition, growth, transmission or clearance. Yet the extent to which SARS-CoV-2 is prone to within-host virus-virus and/or virus-bacteria interactions remains relatively unclear, may continue to evolve, and could have important consequences for epidemiological dynamics and clinical manifestations of antibiotic resistance.(43)

Our model should be considered in the context of several limitations and simplifying assumptions. First, although impacts of COVID-19 on hospital admissions are accounted for, we do not explicitly model community dynamics, nor do we explore different scenarios of SARS-CoV-2 importation from the community. Yet community SARS-CoV-2 outbreaks also result in surges in COVID-19 hospitalizations and the potential overwhelming of healthcare services, and have likely played an important role in driving selection for antibiotic-resistant bacteria since the onset of the pandemic. In the context of prohibited hospital visitation during the first wave of COVID-19, staff but not patient interactions with the community – and potential acquisition of both SARS-CoV-2 infection and bacterial carriage – may further be important drivers of nosocomial transmission dynamics. Second, HCWs are conceptualized here as transient vectors, but HCW colonization can also impact transmission dynamics. In particular, nares are a key site for MRSA colonization,(44) and chronic HCW colonization has been found to drive prolonged nosocomial MRSA outbreaks.(45) Further, our model is conceptualized as applying to commensal bacteria spread through contact and fomites, so we conservatively assumed that face masks have no impact on bacterial transmission. However, respiratory droplets may play a non-negligible role in MRSA transmission,(46) so impacts of COVID-19 responses on MRSA colonization incidence may be underestimated. Finally, our deterministic modelling approach does not allow for stochastic effects like epidemiological extinctions, which are particularly relevant in small populations like hospital wards.

In conclusion, this work has demonstrated how nosocomial SARS-CoV-2 outbreaks influence the epidemiological dynamics of multidrug-resistant bacteria. Our simulation-based approach facilitated the exploration of diverse scenarios and broad parameter spaces, helping to unravel the complexity and context-specificity of such impacts. Results suggest that surges in antibiotic resistance may be expected as a collateral impact of sudden nosocomial outbreaks of novel respiratory pathogens, but that effective implementation of IPC policies that limit nosocomial transmission can mitigate selection for resistance. Given the persistence of SARS-CoV-2 transmission in human populations and high risk of future zoonotic spillover events,(47) investment in pandemic preparedness should be considered a crucial element in the fight against antibiotic resistance.

## Materials and Methods

We used ordinary differential equations (ODEs) to formalize a deterministic, compartmental model describing simultaneous transmission dynamics of SARS-CoV-2 (*V*) and a commensal bacterium (*B*) among inpatients (*pat*) admitted to a healthcare facility, and among HCWs (*hcw*) providing care to patients (**Figure 1**). SARS-CoV-2 infection is characterized by a modified Susceptible-Exposed-Infectious-Recovered (*SEIR*) process, with potential sick leave among symptomatic HCWs. The bacterium is characterized by ecological competition between antibiotic-sensitive strains (*B*^*s*^) and antibiotic-resistant strains (*B*^*R*^). Among patients, we consider exclusive asymptomatic bacterial colonization (*C*^*S*^ or *C*^*R*^), which is potentially cleared naturally (after 1/ *γ* days) or as a result of antibiotic exposure (after 1/*σ* days). *B*^*R*^ is assumed to resist a greater share of antibiotics than *B*^*S*^, but not necessarily all antibiotics 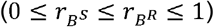, and bears a fitness cost resulting in faster natural clearance 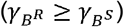. Among HCWs, we consider exclusive transient bacterial carriage (*T*^*S*^ or *T*^*R*^), which is potentially cleared via HCW decontamination (ω), and which depends upon HCW compliance with hand hygiene (*H*) subsequent to HCW-patient contact (*K*^*hck*→*pat*^). HCWs are thus conceptualized as potential vectors for bacterial transmission (HCW colonization is not considered). We assume no within-host ecological interactions between and : bacterial colonization does not directly impact SARS-CoV-2 infection, nor does infection directly impact colonization. See the **Supplementary appendix section 1.1** for full model description and equations. The complete model is programmed in R, and all code is freely available at https://github.com/drmsmith/covR.

### COVID-19 responses: policy vs. caseload

Ten *COVID-19 response parameters* (τ_*i*_) were included in ODEs by changing how individuals flow through the model or modifying relevant parameter values (**Table 1**). *Policy responses* are implemented at a time *t*_*poilicy*_, and reflect evolution of public health policy or practice within the healthcare facility over the course of the epidemic. For each of these responses (τ_*cp*_, τ_*pl*_, τ_*um*_, τ_*hh*_), we assume a phase-in period of duration *t*_*impl*_, during which the policy is gradually adopted, such that it is implemented with full impact at time *t*_*poilicy*_ + *t*_*impl*_. Hence for each policy response the value over time *T*_*t*_(*t*, τ_*t*_)is taken as

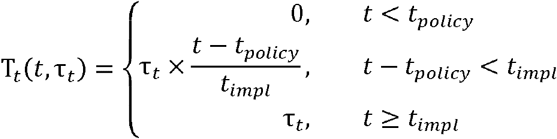

where

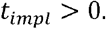

Second, *caseload responses* depend on patient and/or staff SARS-CoV-2 infection prevalence, reflecting impacts of increasing COVID-19 caseloads on provisioning of care. COVID-19 stays (τ_*cs*_) and staff sick leave (τ_*ss*_) impact the numbers of infectious patients and staff in the healthcare facility (see supplementary appendix), while all other dynamic caseload responses (τ_*as*_, τ_*cd*_, τ_*ra*_, τ_*sc*_) scale dynamically with patient infection prevalence. The quantile of the cumulative Beta distribution B_*x*(*t*)_(*α, β*)corresponding with patient SARS-CoV-2 infection prevalence *x*(*t*) is used, fixing *α* = 2 and scaling *β* by the dynamic caseload response τ_*x*_. This gives a modified time- and prevalence-dependent value T_*x*_(*t*),

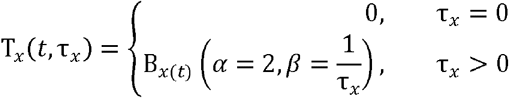

where

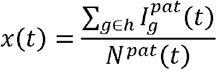

given bacterial colonization status *g* in a set of statuses *h* (see supplementary appendix). The relationship between τ_*x*_, *x*(*t*) and T_*x*_(t) is visualized in **Supplementary figure 1**.

### Simulations

Dynamics were simulated by solving ODEs through numerical integration using the R package *deSolve*.(48) For each simulation, endemic equilibria for bacterial carriage and colonization were found. Then, two cases of SARS-CoV-2 (one patient, one HCW) were introduced into the facility, assuming complete susceptibility to infection among all other individuals, and using estimates of SARS-CoV-2 transmissibility from early 2020 from a long-term care hospital in Paris, France (see parameter values in **Table S1**).(49) Simulations thus aimed at representing poorly anticipated nosocomial SARS-CoV-2 outbreaks in naïve hospital populations, as in the first wave of the COVID-19 pandemic, and were run for a period of 180 days from SARS-CoV-2 introduction.

Monte Carlo simulations were conducted by randomly sampling parameter values from their respective probability distributions, yielding a distinct parameter vector Θ for each simulation (see **Supplementary appendix section 1.2**). Two distinct sets of simulation were considered: (i) generic MRB in generic hospitals, and (ii) case studies of specific bacteria, hospital wards and COVID-19 response scenarios (see **Supplementary appendix section 1.3**). Probability distributions for key parameters underlying these distinct simulation sets are visualized in **Figure S3**. Bootstrap resampling was used to determine the appropriate number of Monte Carlo simulations to conduct (*n* = 500; **Figures S4, S5**).

Epidemiological indicators (Γ) were calculated from simulation outputs, and include the prevalence and incidence of SARS-CoV-2 infection, of patient colonization with *B*^*S*^ and *B*^*R*^, and of HCW carriage of *B*^*S*^ and *B*^*R*^, as well as the cumulative number of patient-days of bacterial colonization and the average resistance rate (the cumulative share of patient colonization due to *B*^*R*^ relative to *B*^*s*^) (see **Supplementary appendix section 1.4**). Multivariate sensitivity analyses were conducted to determine which model parameters drive respective Γ (**Figures S6, S7**).

### Evaluating impacts of COVID-19 on antibiotic resistance

Epidemiological impacts of COVID-19 were assessed by calculating how COVID-19 responses impacted epidemiological indicators in parameter-matched simulations. For Θ^*i*^ corresponding to the *i*^*th*^ Monte Carlo simulation, the model was run both with selected COVID-19 response parameters (τ > 0) and without (τ= 0), and corresponding epidemiological indicators were calculated in their presence 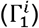 and absence 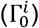. The epidemiological change resulting from COVID-19 responses was thus calculated as the relative difference in each indicator,

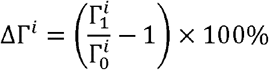

such that positive (negative) values indicate the percentage increase (decrease) in Γ^*i*^ as a result of the COVID-19 response parameters included in Θ^*i*^. Final differences for each indicator are reported as means and 95% uncertainty intervals (95% UI, the 2.5^th^ and 97.5^th^ quantiles) of the resulting distribution ΔΓ^*n*^.

## Acknowledgments

We thank the members of the EMAE-MESuRS Working Group on Nosocomial SARS-CoV-2 Modelling for helpful discussion. This study received funding from the MODCOV project from the Fondation de France (Grant 106059) as part of the alliance framework “Tous unis contre le virus”, the Université Paris-Saclay (AAP Covid-19 2020) and the French government through its National Research Agency project SPHINX-17-CE36-0008-01 and the “Investissement d’Avenir” program, Laboratoire d’Excellence “Integrative Biology of Emerging Infectious Diseases” (Grant ANR-10-LABX-62-IBEID). D.R.M.S. is supported by a Canadian Institutes of Health Research Doctoral Foreign Study Award (Funding Reference Number 164263). The work was also supported directly by internal resources from the French National Institute for Health and Medical Research (Inserm), Institut Pasteur, le Conservatoire National des Arts et Métiers, and l’Université Versailles Saint-Quentin-en-Yvelines/Université Paris-Saclay.

