## Supplementary appendix for "Collateral impacts of pandemic COVID-19 drive the nosocomial spread of antibiotic resistance"

This file includes:

1.1 Model structure
    - Contact behaviour
    - SARS-CoV-2 infection and transmission
    - Bacterial colonization (patients)
    - Bacterial carriage (HCWs)
    - Bacterial clearance
    - Bacterial transmission
    - Hospital demography
    - COVID-19 responses
    - Model equations
    - Supplementary Figures S1, S2
1.2 Model simulation
    - Initialization
    - Monte Carlo simulation
1.3 Model parameterization
    - Simulation set 1: Generic MRB in generic hospitals
    - Simulation set 2: Case studies
    - Transmission rates
    - Contact rates
    - Colonization upon admission
    - Policy responses
    - Parameter tables (Supplementary Tables S1 – S5; Figure S3)
1.4 Epidemiological indicators
    - Prevalence
    - Incidence
    - Patient-days colonized
    - Resistance rate
1.5 Simulation convergence and sensitivity analysis
    - Bootstrapping outcomes (Supplementary Figures S4, S5)
    - Partial rank correlation coefficients (Supplementary Figures S6, S7)
1.6 Supplementary results: Generic MRB in generic hospitals
    - Supplementary Figures S8 – S14
1.7 Supplementary results: Case studies
    - Supplementary Figures S15 – S21

***1.1 Model structure***

The complete model is given below as a system of ordinary differential equations (ODEs), and is preceded by a description of the underlying assumptions informing model structure, including functional impacts of COVID-19 response parameters ($\tau_{i}$) on model dynamics.

*Contact behaviour*

Contact rates of patients ($pat$) and HCWs ($hcw$) are given by

$$\left( \begin{matrix} \kappa^{pat\to pat}(t,\tau_{pl}) & \kappa^{pat\to hcw}(t,\tau_{cd}) \\ \kappa^{hcw\to pat}(t,\tau_{cd}) & \kappa^{hcw\to hcw}(t) \end{matrix} \right)$$

where $\kappa^{i\to j}$ is the average daily number of close-proximity contacts of $i$ with $j$. $\kappa^{hcw\to pat}$ is assumed to scale with $\kappa^{pat\to hcw}$ via the patient:HCW ratio,

$$\kappa^{hcw\to pat}(t,\tau_{cd})=\kappa^{pat\to hcw}(t,\tau_{cd})\times\frac{N^{pat}(t)}{N^{hcw}(t)}$$

such that cohorting disorganization ($\tau_{cd}$) increases both patient contacts with HCWs and HCW contacts with patients (see *COVID-19 responses* section below). In the absence of COVID-19, the patient population size ($N^{pat}$) is assumed to equal the number of beds ($N^{beds}$), reflecting a healthcare facility operating at fixed capacity, and the HCW population size ($N^{hcw}$) is assumed to be constant, reflecting a stable workforce.

*SARS-CoV-2 infection*

SARS-CoV-2 infection follows a Susceptible-Exposed-Infectious-Recovered ($SEIR$) process, determined by an incubation period (lasting $1/\eta$ days) and natural recovery from infectiousness (after $1/\upsilon$ days). SARS-CoV-2 is assumed to transmit via exhaled respiratory aerosols/droplets, with a negligible role for fomite-mediated transmission [(1, 2)](https://sciwheel.com/work/citation?ids=10296329,9752188&pre=&pre=&suf=&suf=&sa=0,0&dbf=0&dbf=0), and to be equally transmissible across all close-proximity contacts $\kappa^{j\to i}$. Susceptible patients ($S^{pat}$) and HCWs ($S^{hcw}$) become infected with SARS-CoV-2 upon contact with infectious individuals ($I^{pat}$, $I^{hcw}$). Assuming frequency-dependent transmission, the force of infection from $i$to $j$ at any time $t$ is the product of $j$’s contacts with $i$ ($\kappa^{j\to i}$), the probability of SARS-CoV-2 transmission per contact ($\pi_{V}$) and the prevalence of SARS-CoV-2 infection among $i$ at time $t$,

$$\lambda_{V}^{i\to j}(t,\tau_{cd},\tau_{pl},\tau_{um})={\pi_{V}\left( t,\tau_{um} \right)\times\kappa}^{j\to i}(t,\tau_{cd},\tau_{pl})\times\frac{\sum_{g\in h} I_{g}^{i}(t)}{N^{i}(t)}$$

where state variables $X_{g}^{k}$ represent individuals' SARS-CoV-2 infection status $X$, demographic category $k$ ($pat$ or $hcw$), and bacterial colonization/carriage status $g$ in a set of statuses $h$, which depend on $k$

$$h=\left\{ \begin{aligned} \{U, C^{S}, C^{R}\}, &k=pat \\ \{D, T^{S}, T^{R}\}, &k=hcw \end{aligned} \right.$$

*Bacterial colonization (patients)*

In the same population, patient colonization with commensal bacterium $B$ is included. Exclusive colonization is assumed to describe ecological competition between co-circulating strains of $B$, those that are antibiotic-sensitive ($B^{S}$) and those that are antibiotic-resistant ($B^{R}$) [(3)](https://sciwheel.com/work/citation?ids=1795601&pre=&suf=&sa=0&dbf=0). Patient colonization is denoted as $C^{S}$ for $B^{S}$ and $C^{R}$ for $B^{R}$, and non-colonization as$U$. Colonization is naturally cleared by the host after $1/\gamma$ days. To reflect metabolic costs ($c$) of bearing and expressing antibiotic resistance genes [(4)](https://sciwheel.com/work/citation?ids=1591706&pre=&suf=&sa=0&dbf=0),$B^{R}$ is naturally cleared by the host more quickly than $B^{S}$

$$\gamma_{B^{R}}=\gamma_{B^{S}}+c$$

*Bacterial carriage (HCWs)*

HCWs are assumed to act as transient vectors ($T^{b}$) for $B^{b}$, potentially carrying bacteria on their hands or equipment after contacting colonized patients or other HCW vectors. $T^{b}$ subsequently transmit $B^{b}$ to uncolonized patients (to $U$, resulting in$C^{b}$) or decontaminated HCWs, i.e. those not already carrying $B^{b}$ (to $D$, resulting in $T^{b}$).

*Bacterial clearance*

Patient colonization is cleared via antibiotic exposure, while transient HCW carriage is cleared via hand hygiene. Specifically, colonization with strain $B^{b}$ is cleared by antibiotics at rate

$$\sigma_{b}(t,\tau_{as},\tau_{cp})= A(t,\tau_{as},\tau_{cp})\times(1-r_{b})\times\theta$$

where $A$ is antibiotic exposure prevalence (the proportion of patients undergoing antibiotic therapy at any time), $r_{b}$represents antibiotic resistance (the proportion of ineffective antibiotics, i.e. the proportion that have no effect on strain $b$), and$\theta$ is the effective colony kill rate (the rate at which effective therapy clears colonization). Antibiotic exposure is assumed to be independent of colonization status, reflecting high rates of bystander selection for commensal bacteria.[(5)](https://sciwheel.com/work/citation?ids=6323434&pre=&suf=&sa=0&dbf=0)

Modified from van Kleef et al.,[(6)](https://sciwheel.com/work/citation?ids=9162958&pre=&suf=&sa=0&dbf=0) transient HCW carriage is cleared by effective hand hygiene at rate

$$\omega(t,\tau_{cd},\tau_{hh})=H(t,\tau_{hh}) \times\kappa^{hcw\to pat}(t,\tau_{cd})\times\frac{N^{hcw}(t)}{N^{pat}(t)}$$

where the rate of HCW decontamination ($\omega$) depends on hand hygiene compliance $H$ (defined here as the probability of successful decontamination after handwashing), and assumes that opportunities for hand hygiene moments increase with staff-patient contacts and scale with the HCW:patient ratio ($N_{hcw}/N_{pat}$). This and other functional assumptions are visualized in **Figure S1**.

*Bacterial transmission*

Like SARS-CoV-2 transmission, frequency-dependent transmission of $B^{b}$ from $i$ to $j$ is assumed to depend on $j$’s contacts with $i$, bacterial transmissibility per contact, and prevalence of colonization (if $i=pat$) or transient carriage (if $i=hcw$) across the set of SARS-CoV-2 infection statuses that excludes HCWs absent due to sick leave ($Y=\{S,E,I,R\}$),

$$\lambda_{b}^{i\to j}\left( t,\tau_{cd},\tau_{pl} \right)=\pi_{B}\times\kappa^{j\to i}(t,\tau_{cd},\tau_{pl})\times\frac{\sum_{X\in Y} X_{b}^{i}(t)}{N^{i}(t)}$$

assuming no difference in transmissibility between strains ($\pi_{B}=\pi_{B^{S}}=\pi_{B^{R}}$). Unlike SARS-CoV-2, $B$ is conceptualized as transmitting via fomites and physical touch, not respiratory droplets. Reflecting that care-based patient-HCW contacts involve closer contact than typical patient-patient or HCW-HCW contacts, a symmetric contact intimacy scaling coefficient $\rho$ is introduced (see force of infection equations).

In addition to inter-individual transmission, endogenous acquisition of $B^{R}$ is assumed to occur at rate $\alpha$ among hosts exposed to the antibiotics against which $B^{R}$ is resistant,

$$\alpha(t,\tau_{as},\tau_{cp})=A(t,\tau_{as},\tau_{cp})\times r_{B^{R}}\times\alpha'$$

reflecting antibiotic selection for outgrowth of subdominant antibiotic-resistant bacteria.[(7, 8)](https://sciwheel.com/work/citation?ids=10533358,926401&pre=&pre=&suf=&suf=&sa=0,0&dbf=0&dbf=0)

*Hospital Demography*

Patients are admitted and discharged at a symmetric rate $\mu$ (given as $\mu^{adm}$ for admission, $\mu^{dis}$ for discharge). $B$ is assumed to be endemic among patients in the community, so a proportion $f_{g}\in\{f_{U}, f_{C^{S}},f_{C^{R}}\}$ are admitted with colonization status $g$. Assuming full patient occupancy under baseline conditions ($N^{pat}= N^{beds}$), admission into and discharge from $X_{g}^{pat}$ is given by

$$\Delta_{X_{g}}(t,\tau_{cs},\tau_{ra},\tau_{sc})= \mu^{adm}(t,\tau_{ra})\times N^{beds}\times f_{g}(t,\tau_{sc})-\mu^{dis}(t,\tau_{cs})\times X_{g}^{pat}(t)$$

Conversely, $V$ is assumed to be novel in the population, so all patients are susceptible to SARS-CoV-2 infection upon admission. Finally, HCWs are assumed to be full-time members of staff, and remain in the facility over simulation time (no turnover). For simplicity, no other interactions with the community are included (e.g. visitors), and no introduction of bacteria into the facility occurs other than through patient admission.

*COVID-19 responses*

The ten COVID-19 responses from **Table 1** are built into the model as follows.

*COVID-19 responses: Antibiotics*

With abandoned stewardship ($\tau_{as}$), the proportion $A$ of patients exposed to antibiotics increases by a factor $A_{as}$ relative to baseline,

$$A(t,\tau_{as}){=A_{base}+A}_{as}(t,\tau_{as})$$

assuming that increases only occur among the proportion $\left( 1-A_{base} \right)$ of patients that are not already exposed to antibiotics

$$A_{as}(t,\tau_{as})=\left( 1-A_{base} \right)\times T_{as}\left( t,\tau_{as} \right)$$

with

$$0\leq A_{base}+A_{as}(t,\tau_{as})\leq1$$

With COVID-19 prophylaxis ($\tau_{cp}$), a symptomatic proportion $\zeta$ of patients with active SARS-CoV-2 infection receive antibiotics. This is assumed to apply only to the share of COVID-19 patients not already receiving antibiotics, where

$$A_{SER}(t,\tau_{as})=A_{base}+A_{as}(t,\tau_{as})$$

is the proportion of Susceptible, Exposed and Recovered patients receiving antibiotics, and

$$A_{I}(t,\tau_{as},\tau_{cp})=A_{SER}(t,\tau_{as})+\left( 1-A_{SER}\left( t,\tau_{as} \right) \right)\times\zeta\times T_{cp}(t,\tau_{cp})$$

is the proportion of symptomatic, infectious SARS-CoV-2-infected patients receiving antibiotics, with

$$0\leq A_{SER}(t,\tau_{as})\leq A_{I}(t,\tau_{as},\tau_{cp})\leq1$$

*COVID-19 responses: Contact*

With cohorting disorganization ($\tau_{cd}$), the daily number of patient-HCW contacts increases, assuming a theoretical maximum tripling of contacts given 100% SARS-CoV-2 infection prevalence,

$$\kappa^{pat\to hcw}(t,\tau_{cd})=\kappa^{pat\to hcw}\times\left( 1+2\times T_{cd}\left( t,\tau_{cd} \right) \right)$$

With patient-lockdown ($\tau_{pl}$), the daily number of patient-patient contacts decreases,

$$\kappa^{pat\to pat}\left( t,\tau_{pl} \right)=\kappa^{pat\to pat}\times\left( 1-T_{pl}\left( t,\tau_{pl} \right) \right)$$

*COVID-19 responses: IPC*

With universal masking ($\tau_{um}$), transmissibility of SARS-CoV-2 decreases by the same proportion from both patients and HCWs,

$$\pi_{V}(t,\tau_{um})=\pi_{V}\times\left( 1-T_{um}\left( t,\tau_{um} \right) \right)$$

With hand hygiene ($\tau_{hh}$), HCW hand hygiene compliance increases relative to the baseline level of compliance,

$$H\left( t,\tau_{hh} \right)=H_{base}+\left( 1-H_{base} \right)\times T_{hh}\left( t,\tau_{hh} \right)$$

*COVID-19 responses: Disease*

With COVID-19 stays ($\tau_{cs}$), the discharge rate decreases for the fraction $\zeta$ of patients who are symptomatic, reflecting their need for ongoing care,

$$\mu_{I}^{dis}(\tau_{cs})=\mu\times\left( 1-\tau_{cs}\times\zeta\right)$$

This results in a longer average length of stay, and a corresponding increase in $N^{pat}$ due to a decrease in the average rate of patient discharge relative to an unchanged rate of patient admission.

With staff sick leave ($\tau_{ss}$), a symptomatic HCWs take sick leave at a daily rate $\tau_{ss}$. This fraction $\zeta$ of HCWs is thus removed from the healthcare facility at rate

$$\delta\left( \tau_{ss} \right)=\zeta\times\tau_{ss}$$

This temporarily decreases $N^{hcw}$ and is given by an ODE representing $SL$, the number of HCWs undergoing sick leave,

$$\frac{dSL}{dt}=\sum_{g\in h} I_{g}^{hcw}(t)\times\delta(\tau_{ss})-SL(t)\times\upsilon_{SL}$$

After sick leave lasting $\upsilon_{SL}^{-1}$ days, HCWs return to work as $R_{D}^{hcw}$ (recovered from SARS-CoV-2 infection, not transiently carrying $B$).

*COVID-19 responses: Admissions*

With reduced admissions ($\tau_{ra}$), the daily rate of patient admission decreases,

$$\mu^{adm}(t,\tau_{ra})=\mu^{adm}\times\left( 1-T_{ra}\left( t,\tau_{ra} \right) \right)$$

reflecting cancellation of elective admissions or redirection to other healthcare facilities during ongoing SARS-CoV-2 epidemics.

With sicker casemix ($\tau_{sc}$), the proportion of newly admitted patients who are uncolonized decreases relative to the baseline proportion already colonized upon admission,

$$f_{U}(t,\tau_{sc})=f_{U}-f_{C^{R}}\times T_{sc}\left( t,\tau_{sc} \right)$$

with corresponding increases in patients admitted with drug-resistant bacteria,

$$f_{C^{R}}\left( t,\tau_{sc} \right)=f_{C^{R}}\times\left( {1+T}_{sc}\left( t,\tau_{sc} \right) \right)$$

assuming no change in the proportion admitted with drug-sensitive bacteria. This reflects a changing denominator in the population of newly admitted patients during SARS-CoV-2 outbreaks, i.e. that individuals who are more sick and who have greater contact with the healthcare system – and who are hence more likely to be colonized with antibiotic-resistant bacteria – represent a greater share of the incoming patient population.

*Model equations*

The complete ODE system integrates the above assumptions and the COVID-19 responses described in the main text. ODEs are written for patients as:

$$\frac{dS_{U}^{pat}}{dt}=\Delta_{S_{U}}\left( t,\tau_{ra},\tau_{sc} \right)-S_{U}^{pat}\left( t \right)\times\left( \alpha\left( t,\tau_{as} \right)+\lambda_{B^{S}}^{pat}\left( t,\tau_{cd},\tau_{pl} \right)+\lambda_{B^{R}}^{pat}\left( t,\tau_{cd},\tau_{pl} \right)+\lambda_{V}^{pat}\left( t,\tau_{cd},\tau_{pl},\tau_{um} \right) \right)+S_{C^{S}}^{pat}\left( t \right)\times\left( \gamma_{B^{S}}+\sigma_{B^{S}}\left( t,\tau_{as} \right) \right)+S_{C^{R}}^{pat}\left( t \right)\times\left( \gamma_{B^{R}}+\sigma_{B^{R}}\left( t,\tau_{as} \right) \right)$$

$$\frac{dS_{C^{S}}^{pat}}{dt}= \Delta_{S_{C^{S}}}\left( t,\tau_{ra} \right)+S_{U}^{pat}\left( t \right)\times\lambda_{B^{S}}^{pat}\left( t,\tau_{cd},\tau_{pl} \right)-S_{C^{S}}^{pat}\left( t \right)\times\left( \lambda_{V}^{pat}\left( t,\tau_{cd},\tau_{pl},\tau_{um} \right)+\gamma_{B^{S}}+\sigma_{B^{S}}\left( t,\tau_{as} \right) \right)$$

$$\frac{dS_{C^{R}}^{pat}}{dt}= \Delta_{S_{C^{R}}}\left( t,\tau_{ra},\tau_{sc} \right)+S_{U}^{pat}\left( t \right)\times\left( \alpha\left( t,\tau_{as} \right)+\lambda_{B^{R}}^{pat}\left( t,\tau_{cd},\tau_{pl} \right) \right)-S_{C^{R}}^{pat}\left( t \right)\times\left( \lambda_{V}^{pat}\left( t,\tau_{cd},\tau_{pl},\tau_{um} \right)+\gamma_{B^{R}}+\sigma_{B^{R}}\left( t,\tau_{as} \right) \right)$$

$$\frac{dE_{U}^{pat}}{dt}= \Delta_{E_{U}}\left( t \right)+S_{U}^{pat}\left( t \right)\times\lambda_{V}^{pat}\left( t,\tau_{cd},\tau_{pl},\tau_{um} \right)-E_{U}^{pat}\left( t \right)\times\left( \alpha\left( t,\tau_{as} \right)+\lambda_{B^{S}}^{pat}\left( t,\tau_{cd},\tau_{pl} \right)+\lambda_{B^{R}}^{pat}\left( t,\tau_{cd},\tau_{pl} \right)+\eta\right)+E_{C^{S}}^{pat}\left( t \right)\times\left( \gamma_{B^{S}}+\sigma_{B^{S}}\left( t,\tau_{as} \right) \right)+E_{C^{R}}^{pat}\left( t \right)\times\left( \gamma_{B^{R}}+\sigma_{B^{R}}\left( t,\tau_{as} \right) \right)$$

$$\frac{dE_{C^{S}}^{pat}}{dt}= \Delta_{E_{C^{S}}}\left( t \right)+S_{C^{S}}^{pat}\left( t \right)\times\lambda_{V}^{pat}\left( t,\tau_{cd},\tau_{pl},\tau_{um} \right)+E_{U}^{pat}\left( t \right)\times\lambda_{B^{S}}^{pat}\left( t,\tau_{cd},\tau_{pl} \right)-E_{C^{S}}^{pat}\left( t \right)\times\left( \gamma_{B^{S}}+\sigma_{B^{S}}\left( t,\tau_{as} \right)+\eta\right)$$

$$\frac{dE_{C^{R}}^{pat}}{dt}= \Delta_{E_{C^{R}}}\left( t \right)+S_{C^{R}}^{pat}\left( t \right)\times\lambda_{V}^{pat}\left( t,\tau_{cd},\tau_{pl},\tau_{um} \right)+E_{U}^{pat}\left( t \right)\times\left( \alpha\left( t,\tau_{as} \right)+\lambda_{B^{R}}^{pat}\left( t,\tau_{cd},\tau_{pl} \right) \right)-E_{C^{R}}^{pat}\left( t \right)\times\left( \gamma_{B^{R}}+\sigma_{B^{R}}\left( t,\tau_{as} \right)+\eta\right)$$

$$\frac{dI_{U}^{pat}}{dt}=\Delta_{I_{U}}\left( t,\tau_{cs} \right)+E_{U}^{pat}\left( t \right)\times\eta-I_{U}^{pat}\left( t \right)\times\left( \alpha\left( t,\tau_{as},\tau_{cp} \right)+\lambda_{B^{S}}^{pat}\left( t,\tau_{cd},\tau_{pl} \right)+\lambda_{B^{R}}^{pat}\left( t,\tau_{cd},\tau_{pl} \right)+\upsilon\right)+I_{C^{S}}^{pat}\left( t \right)\times\left( \gamma_{B^{S}}+\sigma_{B^{S}}\left( t,\tau_{as},\tau_{cp} \right) \right)+I_{C^{R}}^{pat}\left( t \right)\times\left( \gamma_{B^{R}}+\sigma_{B^{R}}\left( t,\tau_{as},\tau_{cp} \right) \right)$$

$$\frac{dI_{C^{S}}^{pat}}{dt}= \Delta_{I_{C^{S}}}\left( t,\tau_{cs} \right)+E_{C^{S}}^{pat}\left( t \right)\times\eta+I_{U}^{pat}\left( t \right)\times\lambda_{B^{S}}^{pat}\left( t,\tau_{cd},\tau_{pl} \right)-I_{C^{S}}^{pat}\left( t \right)\times\left( \gamma_{B^{S}}+\sigma_{B^{S}}\left( t,\tau_{as},\tau_{cp} \right)+\upsilon\right)$$

$$\frac{dI_{C^{R}}^{pat}}{dt}= \Delta_{I_{C^{R}}}\left( t,\tau_{cs} \right)+E_{C^{R}}^{pat}\left( t \right)\times\eta+I_{U}^{pat}\left( t \right)\times\left( {\alpha\left( t,\tau_{as},\tau_{cp} \right)+\lambda}_{B^{R}}^{pat}\left( t,\tau_{cd},\tau_{pl} \right) \right)-I_{C^{R}}^{pat}\left( t \right)\times\left( \gamma_{B^{R}}+\sigma_{B^{R}}\left( t,\tau_{as},\tau_{cp} \right)+\upsilon\right)$$

$$\frac{dR_{U}^{pat}}{dt}= \Delta_{R_{U}}\left( t \right)+I_{U}^{pat}\left( t \right)\times\upsilon-R_{U}^{pat}\left( t \right)\times\left( \alpha\left( t,\tau_{as} \right)+\lambda_{B^{S}}^{pat}\left( t,\tau_{cd},\tau_{pl} \right)+\lambda_{B^{R}}^{pat}\left( t,\tau_{cd},\tau_{pl} \right) \right)+R_{C^{S}}^{pat}\left( t \right)\times\left( \gamma_{B^{S}}+\sigma_{B^{S}}\left( t,\tau_{as} \right) \right)+R_{C^{R}}^{pat}\left( t \right)\times\left( \gamma_{B^{R}}+\sigma_{B^{R}}\left( t,\tau_{as} \right) \right)$$

$$\frac{dR_{C^{S}}^{pat}}{dt}= \Delta_{R_{C^{S}}}\left( t \right)+I_{C^{S}}^{pat}\left( t \right)\times\upsilon+R_{U}^{pat}\left( t \right)\times\lambda_{B^{S}}^{pat}\left( t,\tau_{cd},\tau_{pl} \right)-R_{C^{S}}^{pat}\left( t \right)\times\left( \gamma_{B^{S}}+\sigma_{B^{S}}\left( t,\tau_{as} \right) \right)$$

$$\frac{dR_{C^{R}}^{pat}}{dt}= \Delta_{R_{C^{R}}}\left( t \right)+I_{C^{R}}^{pat}\left( t \right)\times\upsilon+R_{U}^{pat}\left( t \right)\times\left( {\alpha\left( t,\tau_{as} \right)+\lambda}_{B^{R}}^{pat}\left( t,\tau_{cd},\tau_{pl} \right) \right)-R_{C^{R}}^{pat}\left( t \right)\times\left( \gamma_{B^{R}}+\sigma_{B^{R}}\left( t,\tau_{as} \right) \right)$$

and for healthcare workers as

$$\frac{dS_{D}^{hcw}}{dt}=-S_{D}^{hcw}\left( t \right)\times\left( \lambda_{B^{S}}^{hcw}\left( t,\tau_{cd} \right)+\lambda_{B^{R}}^{hcw}\left( t,\tau_{cd} \right)+\lambda_{V}^{hcw}\left( t,\tau_{cd},\tau_{um} \right) \right)+\left( S_{T^{S}}^{hcw}\left( t \right)+S_{T^{R}}^{hcw}\left( t \right) \right)\times\omega\left( t,\tau_{cd},\tau_{hh} \right)$$

$$\frac{dS_{T^{S}}^{hcw}}{dt}=S_{D}^{hcw}\left( t \right)\times\lambda_{B^{S}}^{hcw}\left( t,\tau_{cd} \right)-S_{T^{S}}^{hcw}\left( t \right)\times\left( \lambda_{V}^{hcw}\left( t,\tau_{cd},\tau_{um} \right)+\omega\left( t,\tau_{cd},\tau_{hh} \right) \right)$$

$$\frac{dS_{T^{R}}^{hcw}}{dt}=S_{D}^{hcw}\left( t \right)\times\lambda_{B^{R}}^{hcw}\left( t,\tau_{cd} \right)-S_{T^{R}}^{hcw}\left( t \right)\times\left( \lambda_{V}^{hcw}\left( t,\tau_{cd},\tau_{um} \right)+\omega\left( t,\tau_{cd},\tau_{hh} \right) \right)$$

$$\frac{dE_{D}^{hcw}}{dt}=S_{D}^{hcw}\left( t \right)\times\lambda_{V}^{hcw}\left( t,\tau_{cd},\tau_{um} \right)-E_{D}^{hcw}\left( t \right)\times\left( \lambda_{B^{S}}^{hcw}\left( t,\tau_{cd} \right)+\lambda_{B^{R}}^{hcw}\left( t,\tau_{cd} \right)+\eta\right)+\left( E_{T^{S}}^{hcw}\left( t \right)+E_{T^{R}}^{hcw}\left( t \right) \right)\times\omega\left( t,\tau_{cd},\tau_{hh} \right)$$

$$\frac{dE_{T^{S}}^{hcw}}{dt}=S_{T^{S}}^{hcw}\left( t \right)\times\lambda_{V}^{hcw}\left( t,\tau_{cd},\tau_{um} \right)+E_{D}^{hcw}\left( t \right)\times\lambda_{B^{S}}^{hcw}\left( t,\tau_{cd} \right)-E_{T^{S}}^{hcw}\left( t \right)\times\left( \eta+\omega\left( t,\tau_{cd},\tau_{hh} \right) \right)$$

$$\frac{dE_{T^{R}}^{hcw}}{dt}=S_{T^{R}}^{hcw}\left( t \right)\times\lambda_{V}^{hcw}\left( t,\tau_{cd},\tau_{um} \right)+E_{D}^{hcw}\left( t \right)\times\lambda_{B^{R}}^{hcw}\left( t,\tau_{cd} \right)-E_{T^{R}}^{hcw}\left( t \right)\times\left( \eta+\omega\left( t,\tau_{cd},\tau_{hh} \right) \right)$$

$$\frac{dI_{D}^{hcw}}{dt}=E_{D}^{hcw}\left( t \right)\times\eta-I_{D}^{hcw}\left( t \right)\times\left( \lambda_{B^{S}}^{hcw}\left( t,\tau_{cd} \right)+\lambda_{B^{R}}^{hcw}\left( t,\tau_{cd} \right)+\upsilon+\delta\left( \tau_{ss} \right) \right)+\left( I_{T^{S}}^{hcw}\left( t \right)+I_{T^{R}}^{hcw}\left( t \right) \right)\times\omega\left( t,\tau_{cd},\tau_{hh} \right)$$

$$\frac{dI_{T^{S}}^{hcw}}{dt}=E_{T^{S}}^{hcw}\left( t \right)\times\eta+I_{D}^{hcw}\left( t \right)\times\lambda_{B^{S}}^{hcw}\left( t,\tau_{cd} \right)-I_{T^{S}}^{hcw}\left( t \right)\times\left( \upsilon+\omega\left( t,\tau_{cd},\tau_{hh} \right)+\delta\left( \tau_{ss} \right) \right)$$

$$\frac{dI_{T^{R}}^{hcw}}{dt}=E_{T^{R}}^{hcw}\left( t \right)\times\eta+I_{D}^{hcw}\left( t \right)\times\lambda_{B^{R}}^{hcw}\left( t,\tau_{cd} \right)-I_{T^{R}}^{hcw}\left( t \right)\times\left( \upsilon+\omega\left( t,\tau_{cd},\tau_{hh} \right)+\delta\left( \tau_{ss} \right) \right)$$

$$\frac{dR_{D}^{hcw}}{dt}=I_{D}^{hcw}\left( t \right)\times\upsilon-R_{D}^{hcw}\left( t \right)\times\left( \lambda_{B^{S}}^{hcw}\left( t,\tau_{cd} \right)+\lambda_{B^{R}}^{hcw}\left( t,\tau_{cd} \right) \right)+\left( R_{T^{S}}^{hcw}\left( t \right)+R_{T^{R}}^{hcw}\left( t \right) \right)\times\omega\left( t,\tau_{cd},\tau_{hh} \right)+SL(t)\times\upsilon_{SL}$$

$$\frac{dR_{T^{S}}^{hcw}}{dt}=I_{T^{S}}^{hcw}\left( t \right)\times\upsilon+R_{D}^{hcw}\left( t \right)\times\lambda_{B^{S}}^{hcw}\left( t,\tau_{cd} \right)-R_{T^{S}}^{hcw}\left( t \right)\times\omega\left( t,\tau_{cd},\tau_{hh} \right)$$

$$\frac{dR_{T^{R}}^{hcw}}{dt}=I_{T^{R}}^{hcw}\left( t \right)\times\upsilon+R_{D}^{hcw}\left( t \right)\times\lambda_{B^{R}}^{hcw}\left( t,\tau_{cd} \right)-R_{T^{R}}^{hcw}\left( t \right)\times\omega\left( t,\tau_{cd},\tau_{hh} \right)$$

$$\frac{dSL}{dt}=\left( I_{D}^{hcw}\left( t \right)+I_{D}^{hcw}\left( t \right)+I_{D}^{hcw}\left( t \right) \right)\times\delta(\tau_{ss})-SL\left( t \right)\times\upsilon_{SL}$$

Dynamic forces of infection $\lambda_{m}^{k}(t)$ of each microorganism $m$ to each host type $k$ expand to,

$$\lambda_{B^{S}}^{pat}\left( t,\tau_{cd},\tau_{pl} \right)=\pi_{B}\times\left( \kappa^{pat\to pat}\left( t,\tau_{pl} \right)\times\frac{1}{\rho}\times\sum_{X\in Y} \frac{X_{C^{S}}^{pat}\left( t \right)}{N^{pat}\left( t \right)}+\kappa^{pat\to hcw}\left( t,\tau_{cd} \right)\times\rho\times\sum_{X\in Y} \frac{X_{T^{S}}^{hcw}\left( t \right)}{N^{hcw}\left( t \right)} \right)$$

$$\lambda_{B^{R}}^{pat}\left( t,\tau_{cd},\tau_{pl} \right)=\pi_{B}\times\left( \kappa^{pat\to pat}\left( t,\tau_{pl} \right)\times\frac{1}{\rho}\times\sum_{X\in Y} \frac{X_{C^{R}}^{pat}\left( t \right)}{N^{pat}\left( t \right)}+\kappa^{pat\to hcw}\left( t,\tau_{cd} \right)\times\rho\times\sum_{X\in Y} \frac{X_{T^{R}}^{hcw}\left( t \right)}{N^{hcw}\left( t \right)} \right)$$

$$\lambda_{V}^{pat}\left( t,\tau_{cd},\tau_{pl},\tau_{um} \right)=\pi_{V}\left( t,\tau_{um} \right)\times\left( \kappa^{pat\to pat}\left( t,\tau_{pl} \right)\times\sum_{g\in h} \frac{I_{g}^{pat}\left( t \right)}{N^{pat}\left( t \right)}+\kappa^{pat\to hcw}\left( t,\tau_{cd} \right)\times\sum_{g\in h} \frac{I_{g}^{hcw}\left( t \right)}{N^{hcw}\left( t \right)} \right)$$

$$\lambda_{B^{S}}^{hcw}\left( t,\tau_{cd} \right)=\pi_{B}\times\left( \kappa^{hcw\to hcw}\left( t \right)\times\frac{1}{\rho}\times\sum_{X\in Y} \frac{X_{T^{S}}^{hcw}\left( t \right)}{N^{hcw}\left( t \right)}+\kappa^{hcw\to pat}\left( t,\tau_{cd} \right)\times\rho\times\sum_{X\in Y} \frac{X_{C^{S}}^{pat}(t)}{N^{pat}\left( t \right)} \right)$$

$$\lambda_{B^{R}}^{hcw}\left( t,\tau_{cd} \right)=\pi_{B}\times\left( \kappa^{hcw\to hcw}\left( t \right)\times\frac{1}{\rho}\times\sum_{X\in Y} \frac{X_{T^{R}}^{hcw}\left( t \right)}{N^{hcw}\left( t \right)}+\kappa^{hcw\to pat}\left( t,\tau_{cd} \right)\times\rho\times\sum_{X\in Y} \frac{X_{C^{R}}^{pat}(t)}{N^{pat}\left( t \right)} \right)$$

$$\lambda_{V}^{hcw}\left( t,\tau_{cd},\tau_{um} \right)=\pi_{V}\left( t,\tau_{um} \right)\times\left( \kappa^{hcw\to hcw}\left( t \right)\times\sum_{g\in h} \frac{I_{g}^{hcw}\left( t \right)}{N^{hcw}\left( t \right)}+\kappa^{hcw\to pat}\left( t,\tau_{cd} \right)\times\sum_{g\in h} \frac{I_{g}^{pat}\left( t \right)}{N^{pat}\left( t \right)} \right)$$

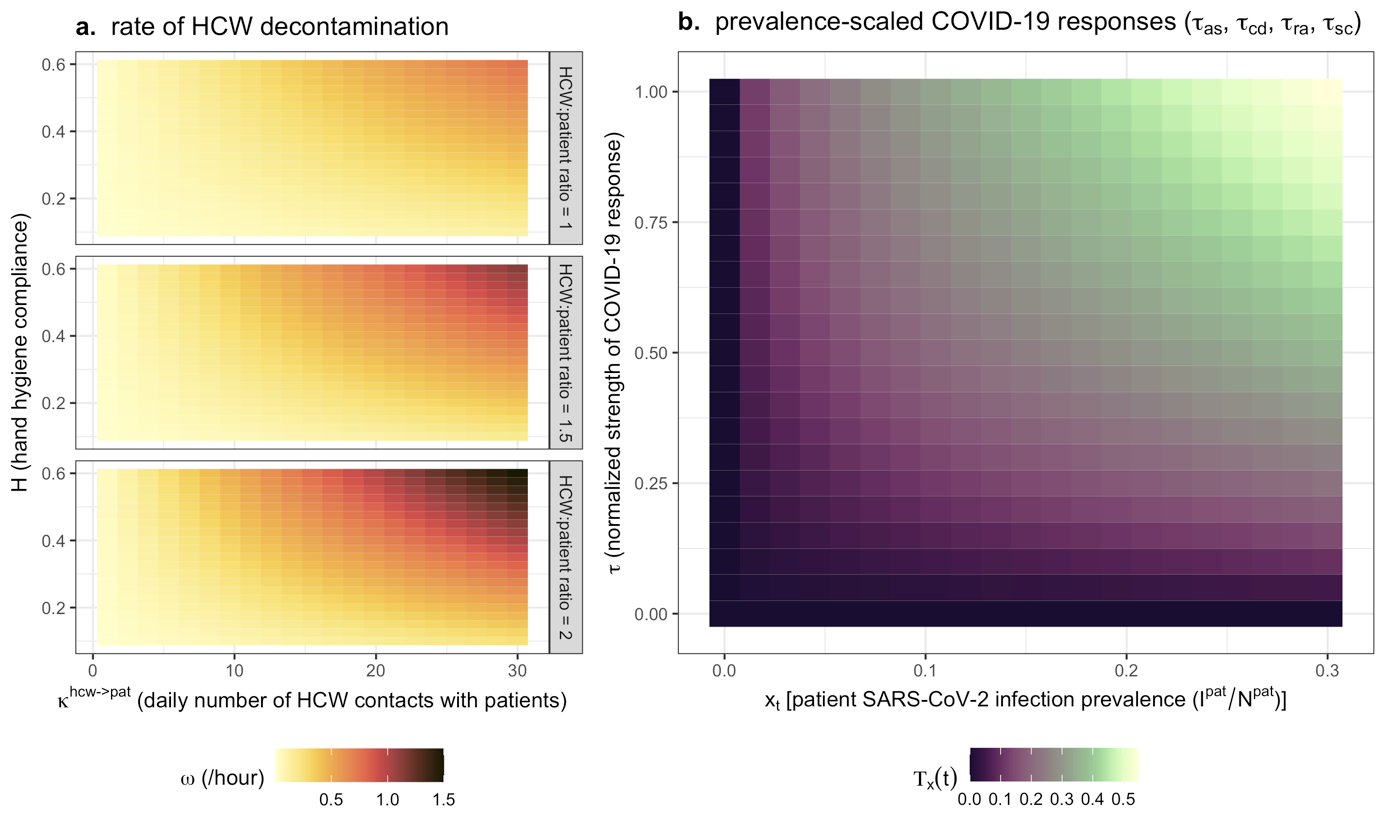

**Figure S1.** Contour plots depicting functional assumptions underlying model dynamics. (**a**) HCWs are recommended to wash their hands after contact with patients, so the rate of successful HCW decontamination $\omega$ (colour) increases with the HCW-to-patient contact rate $\kappa^{hcw\to pat} ($x-axis), but depending on the degree of compliance with handwashing recommendations $H$ (y-axis). Ability to comply with handwashing recommendations is assumed to be more difficult in settings with limited staffing, so $\omega$ also scales positively with the HCW:patient ratio (rows). (**b**) For COVID-19 responses assumed to increase with patient SARS-CoV-2 infection prevalence, the magnitude of the response $T_{x}\left( t \right)$ (colour) depends on the prevalence of infection among patients $x\left( t \right)$ and the assumed value of $\tau$.

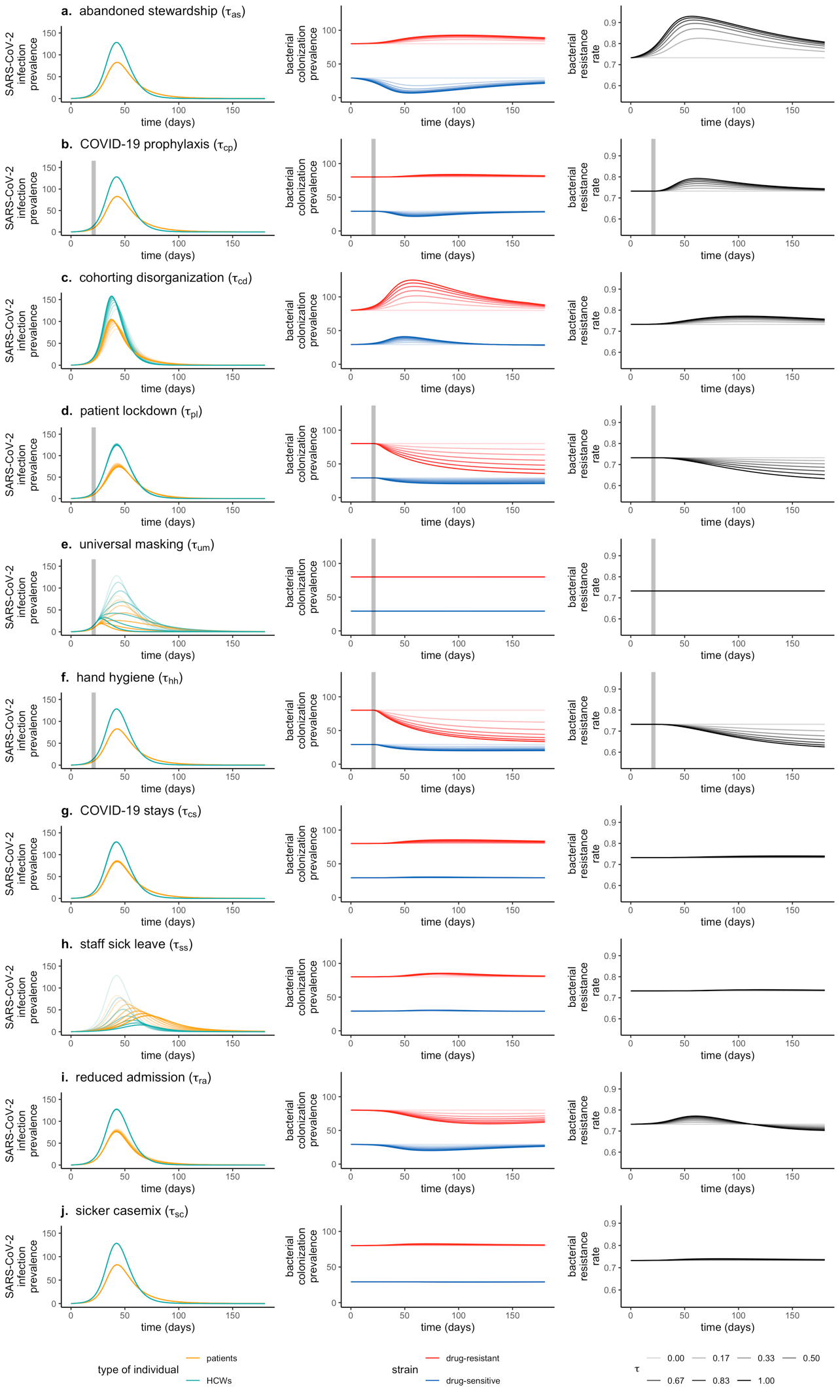

**Figure S2.** COVID-19 responses ($\tau$, rows) drive epidemiological dynamics of SARS-CoV-2 infection among patients and staff (left column), patient colonization with competing antibiotic-sensitive and antibiotic-resistant strains of bacteria (middle column), and the rate of resistance among colonized patients (right column). In all plots, shading represents the strength of the COVID-19 response, ranging from $\tau=0$ (no response, lightest) to $\tau=1$ (maximum response, darkest). All other parameter values are point estimates taken from **Tables S1** and **S2** and match values in **Figure 2** in the main text. ODEs are solved through numerical integration using the R package *deSolve*. Increases in antibiotic use resulting from abandoned stewardship ($\tau_{as}$) or COVID-19 prophylaxis ($\tau_{cp}$) are accompanied by declines in sensitive bacteria and increases in resistant bacteria. Cohorting disorganization ($\tau_{cd}$) increases HCW-patient contacts, while patient lockdown ($\tau_{pl}$) decreases patient-patient contacts, respectively increasing and decreasing opportunities for pathogen transmission. Universal masking ($\tau_{um}$) decreases SARS-CoV-2 transmissibility, while improved hand hygiene compliance ($\tau_{hh}$) facilitates HCW decontamination, reducing bacterial transmission. COVID-19 stays ($\tau_{cs}$) result in a longer average length of stay for patients, while staff sick leaves ($\tau_{ss}$) decrease the number of staff present, with corresponding consequences for transmission of both pathogens. Lastly, reduced admission ($\tau_{ra}$) decreases the daily rate of new patient hospitalization, hence decreasing the patient population size, while a sicker casemix ($\tau_{sc}$) augments the share of patients colonized with resistant bacteria upon admission.

***1.2 Model simulation***

*Initialization*

For each parameter vector $\Theta$, epidemiological dynamics were first simulated in the absence of SARS-CoV-2,

$$\left\{ \begin{aligned} \sum_{g\in h} S_{g}^{pat}(0)=N^{pat}=N^{beds} \\ \sum_{g\in h} \left( E_{g}^{pat}(0)+I_{g}^{pat}\left( 0 \right)+R_{g}^{pat}\left( 0 \right) \right)=0 \\ \sum_{g\in h} S_{g}^{hcw}\left( 0 \right)=N^{hcw} \\ \sum_{g\in h} \left( E_{g}^{hcw}(0)+I_{g}^{hcw}\left( 0 \right)+R_{g}^{hcw}\left( 0 \right)+SL(0) \right)=0 \end{aligned} \right.$$

Given these initial conditions, ODEs were integrated until endemic equilibrium ($\hat{X_{g}^{k}}$) was found, representing stable baseline prevalence of bacterial colonization and HCW carriage for the bacterium and healthcare setting described by $\Theta$. Then, after resetting simulation time ($t_{0}=0$) and substituting endemic equilibria as initial conditions,

$$X_{g}^{k}\left( t_{0} \right)=\hat{X_{g}^{k}}$$

one susceptible patient was replaced by one SARS-CoV-2-exposed patient, and one susceptible HCW $S_{U}^{hcw}$ by one SARS-CoV-2-exposed HCW $E_{U}^{hcw}$,

$$\left\{ \begin{aligned} S_{U}^{pat}\left( t_{0} \right)=\hat{S_{U}^{pat}}-1 \\ E_{U}^{pat}\left( t_{0} \right)=1 \\ \begin{aligned} S_{U}^{hcw}\left( t_{0} \right)=\hat{S_{U}^{hcw}}-1 \\ E_{U}^{hcw}\left( t_{0} \right)=1 \end{aligned} \end{aligned} \right.$$

representing two index cases of SARS-CoV-2 infection at simulation outset in an otherwise immunologically naïve population. ODEs were then integrated twice over $t=180$ days: (i) first in the absence of COVID-19 responses ($\tau=0$), yielding equilibrium dynamics of $B^{S}$ and $B^{R}$ over a roughly 6-month period in the context of an unmitigated nosocomial outbreak of $V$ that has no mechanistic impact on $B$; and (ii) second after substituting selected COVID-19 responses ($\tau>0$) into $\Theta$, yielding epidemiological dynamics of $B^{S}$ and $B^{R}$ in the context of COVID-19 responses that mechanistically link the dynamics of $V$ and $B$.

*Monte Carlo simulation*

Parameter uncertainty was accounted for by conducting Monte Carlo simulations. For each $i^{th}$ simulation, a distinct parameter vector $\Theta^{i}$ was generated by randomly drawing values from the probability distributions defined for each parameter. Each complete parameter vector $\Theta^{i}$ is composed of sub-vectors accounting for SARS-CoV-2 parameters ($\Theta_{V}^{i}$), bacterial parameters ($\Theta_{B}^{i}$), healthcare facility parameters ($\Theta_{F}^{i}$), and COVID-19 response parameters ($\Theta_{\tau}^{i}$). Two sets of Monte Carlo simulations were conducted, the first considering generic multidrug-resistant bacteria (MRB) in generic hospital settings, and the second considering case studies of specific bacteria, hospital wards and COVID-19 response scenarios. Model parameterization for each simulation set is described below.

***1.3 Model parameterization***

Parameter tables for model simulations are given below in **Tables S1 – S5**. These are preceded by a description of the two main lots of simulation conducted, as well as details about calculation of particular parameter values.

*Simulation set 1: Generic MRB in generic hospitals*

In the first simulation set, SARS-CoV-2 parameters were fixed ($\Theta_{V}$), while probability distributions with high variance were defined for all parameters pertaining to bacterial ecology and healthcare facility characteristics, such that each simulated bacterium ($\Theta_{B}^{i}$) represents a “generic MRB” and each simulated facility ($\Theta_{F}^{i}$) represents a “generic hospital”. Parameter distributions were primarily adapted from previous modelling studies of antibiotic-resistant bacteria in healthcare settings, with probability laws defined to cover the broad parameter spaces considered therein (**Table S2**).

For this simulation set, COVID-19 response parameters ($\Theta_{\tau}$) were not varied randomly, but were instead fixed across $j=18$ distinct combinations of $\tau$. The magnitude of each COVID-19 response parameter was fixed at $\tau=0.5$ when included and $\tau=0$ when excluded. We included: (i) each $\tau$ independently; (ii) each category simultaneously, i.e. antibiotic responses ($\tau_{cp}$, $\tau_{as}$), contact responses ($\tau_{pl}$, $\tau_{cd}$), IPC responses ($\tau_{um}$, $\tau_{hh}$), disease responses ($\tau_{cc}$, $\tau_{ss}$), and admission responses ($\tau_{ra}$, $\tau_{sc}$); policy responses simultaneously ($\tau_{cp},\tau_{pl},\tau_{um},\tau_{hh}$); caseload responses simultaneously ($\tau_{as},\tau_{cd},\tau_{cc},\tau_{ss},\tau_{ra},\tau_{sc}$); and finally all ten COVID-19 responses simultaneously. Each parameter vector ($\Theta^{i,j}$) thus represents a distinct, theoretical species-setting pair in the context of the $j^{th}$ COVID-19 response(s), and final outcome distributions ${\Delta\Gamma}^{n,j}$ represent impacts of that response on epidemiological dynamics of diverse bacterial species across diverse healthcare settings. Over $n=500$ MRB-hospital pairs and 18 COVID-19 responses, a total 9,000 distinct parameter vectors were generated and corresponding simulations conducted. The appropriate value of $n$ was determined through bootstrap resampling (see below, **Supplementary appendix section 1.5**).

*Simulation set 2: Case studies*

In the second simulation set, parameter estimates from the literature were used to define probability distributions for specific bacterial species, hospital wards and COVID-19 response scenarios, while also varying selected SARS-CoV-2 parameters. First, selected bacteria were (i) *Staphylococcus aureus*, where methicillin-resistant *S. aureus* (MRSA) was assumed to compete ecologically with methicillin-sensitive *S. aureus* (MSSA); and (ii) *Escherichia coli*, where extended-spectrum beta-lactamase-producing *E. coli* (ESBL-*E. coli*) was assumed to compete with “antibiotic sensitive” *E. coli* (**Table S3**). Second, selected hospital ward settings were a geriatric rehabilitation ward, a short-stay geriatric ward, and a general paediatric ward. Detailed estimates of inter-individual contact behaviour recorded through wearable radio frequency identification devices were available for each of these wards, and were supplemented with estimates of pre-pandemic antibiotic exposure prevalence ($A_{base}$) and HCW hand hygiene compliance ($H_{base}$) from other studies (**Table S4**). Third, observational studies from the first wave of the COVID-19 pandemic, modelling studies and interventional trials were used to define characteristics of the three considered COVID-19 response scenarios (**Table S5**). We assumed no difference in caseload responses across scenarios, on the basis that these are more difficult to anticipate and less likely to vary across healthcare facilities than policy responses, particularly in the context of the first wave of COVID-19. Finally, SARS-CoV-2 parameters varied across a pre-determined range were the SARS-CoV-2 transmission rate ($\beta_{V}\in\{0.76,1.28,2.4,3.6,4.8\}$) and timing of policy implementation ($t_{policy}\in\{7,14,21,28,35\}$).

Parameter sub-vectors ($\Theta_{B}^{i}$, $\Theta_{F}^{i}$, $\Theta_{\tau}^{i}$, $\Theta_{V}^{i}$) were generated independently and then combined for each simulation to ensure perfect parameter matching across model runs. For instance, identical $\Theta_{B}^{n}$ underlie simulations of MSRA in all wards and scenarios, and identical $\Theta_{\tau}^{n}$ underlie simulations of overwhelmed responses for all bacteria and wards. Over $n=500$ sub-vectors of each of two bacteria, three wards and three COVID-19 response scenarios – each evaluated in the context of 25 sub-vectors of SARS-CoV-2 parameters – a total 225,000 distinct parameter vectors were generated and corresponding simulations conducted.

*Transmission rates*

It was necessary to translate daily transmission rates available in the literature ($\beta$) into transmission rates per contact ($\pi$). For SARS-CoV-2 transmission, selected $\beta_{V}$ estimates from the literature reflect cumulative transmission among both patients and HCWs, so $\pi_{V}$ was calculated accounting for the cumulative daily number of contacts across both,

$$\pi_{V}=\frac{\beta_{V}}{\kappa^{pat\to pat}+\kappa^{pat\to hcw}+\kappa^{hcw\to hcw}+\kappa^{hcw\to pat}}$$

For bacterial transmission, $\beta_{B}$ estimates from the literature exclusively reflect patient colonization acquisition, so $\pi_{B}$ was calculated accounting only for contacts with patients,

$$\pi_{B}=\frac{\beta_{B}}{\kappa^{pat\to pat}+\kappa^{hcw\to pat}}$$

*Contact rates*

For case studies, it was necessary to translate contact data from specific healthcare institutions available in the literature into model parameter values.

For the geriatric rehabilitation ward, we used data from Duval *et al*.[(9)](https://sciwheel.com/work/citation?ids=8684589&pre=&suf=&sa=0&dbf=0) In this study, HCWs represent a group combining caregivers, nurses, nurse managers and nurse interns. Contact behaviours were reported across five distinct wards, but here we extracted demographic and contact data specifically for the geriatric ward. For admission and discharge into this ward, we used the facility-wide estimate, $\mu=1/{49 \mathrm{days}^{-1}}$.

For the short-stay geriatric ward, we used data from Vanhems *et al*.[(10)](https://sciwheel.com/work/citation?ids=732780&pre=&suf=&sa=0&dbf=0) We included both doctors ($d$) and nurses ($n$) as HCWs, and calculated contact rates as,

$$\kappa^{pat\to pat}=\kappa^{pat\to pat}$$

$$\kappa^{pat\to hcw}=\kappa^{pat\to nur}+\kappa^{pat\to doc}$$

$$\kappa^{hcw\to pat}=\kappa^{pat\to hcw}\times\frac{N^{pat}}{N^{nur}+N^{doc}}$$

$$\kappa^{hcw\to hcw}=\left( \kappa^{nur\to doc}+\kappa^{nur\to nur} \right)\times\frac{N^{nur}}{N^{nur}+N^{doc}}+\left( \kappa^{doc\to nur}+\kappa^{doc\to doc} \right)\times\frac{N^{doc}}{N^{nur}+N^{doc}}$$

Admission rate was calculated according to available parameters as,

$$\mu=\frac{N^{admissions}}{N^{days}}\times\frac{1}{N^{beds}}=\frac{31}{4}\times\frac{1}{19}=0.408$$

For the general paediatric ward, we used data from Isella *et al*. We included assistants ($ass$), doctors ($doc$) and nurses ($nur$) as HCWs, and calculated contact rates as,

$$\kappa^{pat\to hcw}=\kappa^{pat\to ass}+\kappa^{pat\to doc}+\kappa^{pat\to nur}$$

$$\kappa^{hcw\to pat}=\kappa^{pat\to hcw}\times\frac{N^{pat}}{N^{ass}+N^{nur}+N^{doc}}$$

$$\kappa^{hcw\to hcw}=\left( \kappa^{ass\to ass}+\kappa^{ass\to nur}+\kappa^{ass\to doc} \right)\times\frac{N^{ass}}{{N^{ass}+N}^{nur}+N^{doc}}+\left( {\kappa^{nur\to ass}+\kappa}^{nur\to nur}+\kappa^{nur\to doc} \right)\times\frac{N^{nur}}{{N^{ass}+N}^{nur}+N^{doc}}+\left( {\kappa^{doc\to ass}+\kappa}^{doc\to nur}+\kappa^{doc\to doc} \right)\times\frac{N^{doc}}{{N^{ass}+N}^{nur}+N^{doc}}$$

*Colonization upon admission*

For case studies, to estimate admission fractions for both strains, we distinguished between the proportion of patients admitted carrying any strain of the focal species ($f_{C}$), and, among those carriers, the proportion bearing a strain with the focal resistance element ($f_{R}$). As such, admission fractions were calculated as,

$$f_{C^{S}}=f_{C}\times(1-f_{R})$$

$$f_{C^{R}}=f_{C}\times f_{R}$$

$$f_{U}=1-f_{C^{S}}-f_{C^{R}}$$

*Policy responses*

Observed increases in hand hygiene compliance have been reported in the literature in various settings both during the COVID-19 pandemic and prior to the pandemic in interventional trials. Estimated pre ($H_{0}$) and post ($H_{1}$) values of compliance were used to estimate $\tau_{hh}$,

$$\tau_{hh}=\frac{H_{1}-H_{0}}{(1-H_{0})}$$

Efficacy of face masks for prevention of the expulsion of respiratory aerosols have been estimated in the literature. As in Bartsch *et al*.,[(11)](https://sciwheel.com/work/citation?ids=12629215&pre=&suf=&sa=0&dbf=0) we assumed,

$$\tau_{um}=f\times F$$

where $f$ is the proportion of aerosol droplets blocked by the correctly worn mask, and $F$ is the degree of compliance with correct mask-wearing procedures.

*Parameter tables*

**Table S1**. Parameter values for SARS-CoV-2 infection and transmission, and timing of SARS-CoV-2 control policy responses.

| **Parameter** | | | **Parameter value (range)** | **Notes and references** |
| --- | --- | --- | --- | --- |
| Symbol | Name | Unit |  |  |
| $\beta_{V}$ | transmission rate | day^-1^ | $1.28 (0.76-4.8)$ | Estimate from a long-term care hospital in France in April 2020; range considered in sensitivity analysis [(12)](https://sciwheel.com/work/citation?ids=13046772&pre=&suf=&sa=0&dbf=0) |
| $\eta$ | incubation rate | day^-1^ | $\frac{1}{5}$ | [(13, 14)](https://sciwheel.com/work/citation?ids=8159772,8378827&pre=&pre=&suf=&suf=&sa=0,0&dbf=0&dbf=0) |
| $\upsilon$ | recovery rate | day^-1^ | $\frac{1}{7}$ | Infectiousness estimated to decline within 7 days [(15)](https://sciwheel.com/work/citation?ids=8690687&pre=&suf=&sa=0&dbf=0) |
| $\upsilon_{SL}$ | return from sick leave rate | day^-1^ | $\frac{1}{7}$ | Sick leave duration assumed equal to duration of COVID-19 infection |
| $\zeta$ | proportion symptomatic | % | $52.5$ | In a systematic review and meta-analysis, the proportion of nursing home residents and staff who were symptomatic [(16)](https://sciwheel.com/work/citation?ids=12855219&pre=&suf=&sa=0&dbf=0) |
| $t_{policy}$ | time to policy change | days | $21 (7- 35)$ | Assumed; range considered in sensitivity analysis |
| $t_{impl}$ | time to full policy implementation | days | $7$ | Approximate delay between implementation of public health measures and observed impact on SARS-CoV-2 transmission prevention in hospital data [(12)](https://sciwheel.com/work/citation?ids=13046772&pre=&suf=&sa=0&dbf=0) |

**Table S2**. Parameter distributions for generic MRB and generic hospitals. Point values correspond to values assumed in **Figure 2** in the main text and **Figure S2** in the supplement, while parameter distributions correspond to Monte Carlo simulations. Where possible, parameter distributions were defined to cover the range of values considered in previous modelling studies cited in the notes and references column. For distributions: $\mathcal{U}\left( a,b \right)$ indicates a Uniform distribution with minimum $a$ and maximum$b$; $B\left( \alpha,\beta\right)$ indicates a Beta distribution with shape parameters $\alpha$ and $\beta$; and $\Gamma\left( X,Y \right)$ indicates a Gamma distribution with shape parameter $\alpha$ and rate parameter $\beta$. Point values indicated as being calculated from hospital data are based upon data from the long-term care hospital described in Shirreff *et al*. in which the transmission rate of SARS-CoV-2 was estimated.[(12)](https://sciwheel.com/work/citation?ids=13046772&pre=&suf=&sa=0&dbf=0) LOS = length of stay, HCW = healthcare worker, S strain = antibiotic-sensitive strain ($B^{S}$), R strain = antibiotic-resistant strain ($B^{R}$).

| **Parameter** | | | **Point value** | **Parameter distribution** | **Notes and references** |
| --- | --- | --- | --- | --- | --- |
| Symbol | Name | Unit |  |  |  |
| $\beta_{B}$ | transmission rate | day^-1^ | $0.2$ | $\mathrm{Lognormal}$  $\left( \log\left( 0.2 \right),\log\left( 3 \right) \right)$ | [(6, 17–22)](https://sciwheel.com/work/citation?ids=9162958,6089731,11770059,5333453,1795566,12405002,2894755&pre=&pre=&pre=&pre=&pre=&pre=&pre=&suf=&suf=&suf=&suf=&suf=&suf=&suf=&sa=0,0,0,0,0,0,0&dbf=0&dbf=0&dbf=0&dbf=0&dbf=0&dbf=0&dbf=0) |
| $\alpha$ | endogenous acquisition rate | day^-1^ | $0.01$ | $\Gamma\left( 0.4,10 \right)$ | [(17, 18, 21, 23)](https://sciwheel.com/work/citation?ids=6089731,6258975,11770059,12405002&pre=&pre=&pre=&pre=&suf=&suf=&suf=&suf=&sa=0,0,0,0&dbf=0&dbf=0&dbf=0&dbf=0) |
| $\gamma$ | natural clearance rate | day^-1^ | $0.03$ | $\mathrm{Lognormal}$  $\left( \log\left( 0.03 \right),\log\left( 1.5 \right) \right)$ | [(6, 18, 20, 21, 23)](https://sciwheel.com/work/citation?ids=9162958,6258975,11770059,1795566,12405002&pre=&pre=&pre=&pre=&pre=&suf=&suf=&suf=&suf=&suf=&sa=0,0,0,0,0&dbf=0&dbf=0&dbf=0&dbf=0&dbf=0) |
| $c$ | fitness cost of resistance | / | $0.1$ | $B\left( 0.5,2 \right)$ | [(17, 19, 22, 23)](https://sciwheel.com/work/citation?ids=6089731,6258975,5333453,2894755&pre=&pre=&pre=&pre=&suf=&suf=&suf=&suf=&sa=0,0,0,0&dbf=0&dbf=0&dbf=0&dbf=0) |
| $f_{C^{S}}$ | admission fraction (colonized with S strain) | % | $15$ | $\mathcal{U}\left( 5,30 \right)$ | Assumed |
| $f_{C^{R}}$ | admission fraction (colonized with R strain) | % | $5$ | $\mathcal{U}\left( 0.1,15 \right)$ | Assumed |
| $r_{S}$ | antibiotic resistance level (S strain) | % | $10$ | $\mathcal{U}\left( 0,25 \right)$ | Assumed |
| $r_{R}$ | antibiotic resistance level (R strain) | % | $90$ | $\mathcal{U}\left( 75,100 \right)$ | Assumed |
| $\theta$ | effective colony kill rate | day^-1^ | $\frac{1}{4}$ | $B\left( 0.8,2 \right)$ | [(18–23)](https://sciwheel.com/work/citation?ids=6258975,11770059,5333453,1795566,12405002,2894755&pre=&pre=&pre=&pre=&pre=&pre=&suf=&suf=&suf=&suf=&suf=&suf=&sa=0,0,0,0,0,0&dbf=0&dbf=0&dbf=0&dbf=0&dbf=0&dbf=0) |
| $\mu$ | admission & discharge rate | day^-1^ | $\frac{1}{80}$ | $\mathrm{Lognormal}$  $\left( \log\left( 10 \right),\log\left( 3 \right) \right)$ | Point value is the inverse LOS from hospital data [(6, 17, 20–25)](https://sciwheel.com/work/citation?ids=9162958,12143649,6089731,6258975,6260479,1795566,12405002,2894755&pre=&pre=&pre=&pre=&pre=&pre=&pre=&pre=&suf=&suf=&suf=&suf=&suf=&suf=&suf=&suf=&sa=0,0,0,0,0,0,0,0&dbf=0&dbf=0&dbf=0&dbf=0&dbf=0&dbf=0&dbf=0&dbf=0) |
| $N^{beds}$ | # of beds in the facility | / | $350$ | $\mathcal{U}\left( 100,1000 \right)$ | Point value is the number of beds in hospital data |
| $N^{hcw}$ | # of HCWs working/day | / | $529$ | $N^{beds}\times\mathcal{U}\left( 0.25,3 \right)$ | Point value is the number of HCWs in hospital data |
| $\kappa^{pat\to pat}$ | # of contacts (patient-to-patient) | day^-1^ | $5$ | $\mathrm{Lognormal}$  $\left( \log\left( 5 \right),\log\left( 2 \right) \right)$ | [(9, 10, 26)](https://sciwheel.com/work/citation?ids=8684589,732780,732383&pre=&pre=&pre=&suf=&suf=&suf=&sa=0,0,0&dbf=0&dbf=0&dbf=0) |
| $\kappa^{hcw\to hcw}$ | # of contacts (HCW-to-HCW) | day^-1^ | $10$ | $\mathrm{Lognormal}$  $\left( \log\left( 15 \right),\log\left( 2 \right) \right)$ | [(9, 10, 26)](https://sciwheel.com/work/citation?ids=8684589,732780,732383&pre=&pre=&pre=&suf=&suf=&suf=&sa=0,0,0&dbf=0&dbf=0&dbf=0) |
| $\kappa^{pat\to hcw}$ | # of contacts (patient-to-HCW) | day^-1^ | $15$ | $\mathrm{Lognormal}$  $\left( \log\left( 15 \right),\log\left( 2 \right) \right)$ | [(9, 10, 26)](https://sciwheel.com/work/citation?ids=8684589,732780,732383&pre=&pre=&pre=&suf=&suf=&suf=&sa=0,0,0&dbf=0&dbf=0&dbf=0) |
| $\rho$ | contact intimacy scaling coefficient | / | $3$ | $\mathcal{U}\left( 1,5 \right)$ | Adapted from [(27)](https://sciwheel.com/work/citation?ids=9092216&pre=&suf=&sa=0&dbf=0) |
| $H_{base}$ | hand hygiene compliance | % | $40$ | $\mathcal{U}\left( 20,60 \right)$ | Centred on mean from [(6)](https://sciwheel.com/work/citation?ids=9162958&pre=&suf=&sa=0&dbf=0) |
| $A_{base}$ | antibiotic exposure prevalence | % | $10$ | $B\left( 1.5,5 \right)\times100$ | Assumed |

**Table S3.** Parameter distributions for bacterial species considered in case studies (*Staphylococcus aureus* and *Escherichia coli*). For distributions: $\mathcal{N}\left( \mu,\sigma\right)$ indicates a Normal distribution with mean $\mu$ and standard deviation $\sigma$; $\mathcal{U}\left( a,b \right)$ indicates a Uniform distribution with minimum $a$ and maximum$b$; and $\mathcal{C}\left( x_{0},\gamma\right)$ indicates a Cauchy distribution with location parameter $x_{0}$ and scale parameter$\gamma$. When parameter uncertainty was unavailable, normal distributions with standard deviation equal to 10% of the mean were assumed. MSSA = methicillin-sensitive *S. aureus*; MRSA = methicillin-resistant *S. aureus*; ESBL = extended-spectrum beta-lactamase.

| **Parameter** | | | **Species** | **Parameter distribution** | **Notes and references** |
| --- | --- | --- | --- | --- | --- |
| Symbol | Name | Unit |  |  |  |
| $\beta$ | transmission rate | day^-1^ | *S. aureus* | $\mathcal{N}\left( 0.057,0.0057 \right)$ | Estimated in Norwegian hospitals [(28)](https://sciwheel.com/work/citation?ids=7377956&pre=&suf=&sa=0&dbf=0) |
|  |  |  | *E. coli* | $\mathcal{N}\left( 0.0078,0.0035 \right)$ | Estimated in 13 European ICUs [(29)](https://sciwheel.com/work/citation?ids=7174876&pre=&suf=&sa=0&dbf=0) |
| $\alpha$ | endogenous acquisition rate | day^-1^ | *S. aureus* | $\mathcal{N}\left( 0.0016,0.0057 \right)\mathcal{\times C}\left( 2.97,0.28 \right)$ | Proxy measure: the estimated rate of progression from colonization to infection in hospital patients, multiplied by excess risk of endogenous outgrowth subsequent to antibiotic exposure [(18, 28, 30)](https://sciwheel.com/work/citation?ids=5496695,7377956,11770059&pre=&pre=&pre=&suf=&suf=&suf=&sa=0,0,0&dbf=0&dbf=0&dbf=0) |
|  |  |  | *E. coli* | $\mathcal{N}\left( 0.0024,0.000663 \right)\mathcal{\times C}\left( 11.80,0.80 \right)$ | Proxy measure: rate of endogenous outgrowth estimated in 13 European ICUs, multiplied by excess risk of endogenous outgrowth subsequent to antibiotic exposure [(18, 29)](https://sciwheel.com/work/citation?ids=7174876,11770059&pre=&pre=&suf=&suf=&sa=0,0&dbf=0&dbf=0) |
| $\gamma$ | natural clearance rate | day^-1^ | *S. aureus* | $\frac{1}{\mathcal{N}\left( 287,17.9 \right)}$ | Inverse of estimated duration of colonization [(31)](https://sciwheel.com/work/citation?ids=4942267&pre=&suf=&sa=0&dbf=0) |
|  |  |  | *E. coli* | $\mathcal{N}\left( 0.00269,0.000216 \right)$ | Exponential decay model fit to longitudinal colonization data [(18, 32)](https://sciwheel.com/work/citation?ids=2914123,11770059&pre=&pre=&suf=&suf=&sa=0,0&dbf=0&dbf=0) |
| $c$ | fitness cost of resistance | / | *S. aureus* | $\mathcal{N}\left( 0.2,0.02 \right)$ | Growth cultures from French hospital isolates showed 20% fitness benefit to MSSA over MRSA strains [(33, 34)](https://sciwheel.com/work/citation?ids=2894934,2679877&pre=&pre=&suf=&suf=&sa=0,0&dbf=0&dbf=0) |
|  |  |  | *E. coli* | $0$ | No fitness cost observed in vitro [(35)](https://sciwheel.com/work/citation?ids=6683421&pre=&suf=&sa=0&dbf=0) |
| $f_{C}$ | admission fraction (colonized) | % | *S. aureus* | $\mathcal{N}\left( 7.57,0.3364 \right)$ | Estimated as the proportion of patients arriving to a French hospital with MRSA colonization, divided by the estimated proportion of *S. aureus* strains that are methicillin-resistant in France [(36, 37)](https://sciwheel.com/work/citation?ids=7643014,8810222&pre=&pre=&suf=&suf=&sa=0,0&dbf=0&dbf=0) |
|  |  |  | *E. coli* | $\mathcal{N}\left( 27.5,1.4 \right)$ | Estimated as the proportion of patients arriving to 13 European ICUs with ESBL-*E. coli* carriage, divided by the estimated proportion of *E. coli* that are ESBL-producing in a Hungarian hospital [(29, 38)](https://sciwheel.com/work/citation?ids=6250840,7174876&pre=&pre=&suf=&suf=&sa=0,0&dbf=0&dbf=0) |
| $f_{R}$ | admission fraction (bearing R strain) | % | *S. aureus* | $\mathcal{N}\left( 16,1.6 \right)$ | Proportion of *S. aureus* that are methicillin-resistant in a French hospital setting [(36)](https://sciwheel.com/work/citation?ids=7643014&pre=&suf=&sa=0&dbf=0) |
|  |  |  | *E. coli* | $\mathcal{N}\left( 11.9,4.13 \right)$ | Proportion of fecal *E. coli* that were ESBL-producing from a non-outbreak setting [(38)](https://sciwheel.com/work/citation?ids=6250840&pre=&suf=&sa=0&dbf=0) |
| $r_{S}$ | antibiotic resistance level (S strain) | % | *S. aureus* | $\mathcal{U}(17.2,48.9)$ | Cumulative MSSA resistance level across simulated antibiotic consumption data and standard hospital antibiograms [(18)](https://sciwheel.com/work/citation?ids=11770059&pre=&suf=&sa=0&dbf=0) |
|  |  |  | *E. coli* | $\mathcal{U}(9.6,36.5)$ | Cumulative *E. coli* resistance level across simulated antibiotic consumption data and standard hospital antibiograms [(18)](https://sciwheel.com/work/citation?ids=11770059&pre=&suf=&sa=0&dbf=0) |
| $r_{R}$ | antibiotic resistance level (R strain) | % | *S. aureus* | $\mathcal{U}(90.8,98.2)$ | Cumulative MRSA resistance level across simulated antibiotic consumption data and standard hospital antibiograms [(18)](https://sciwheel.com/work/citation?ids=11770059&pre=&suf=&sa=0&dbf=0) |
|  |  |  | *E. coli* | $\mathcal{U}(77.4,92.2)$ | Cumulative ESBL-*E. coli* resistance level across simulated antibiotic consumption data and standard hospital antibiograms [(18)](https://sciwheel.com/work/citation?ids=11770059&pre=&suf=&sa=0&dbf=0) |
| $\theta$ | effective colony kill rate | day^-1^ | *S. aureus* | $\frac{1}{\mathcal{U}\left( 1,10 \right)}$ | Modelling study [(23)](https://sciwheel.com/work/citation?ids=6258975&pre=&suf=&sa=0&dbf=0) |
|  |  |  | *E. coli* | $\frac{1}{\mathcal{U}\left( 1,10 \right)}$ | Modelling study [(23)](https://sciwheel.com/work/citation?ids=6258975&pre=&suf=&sa=0&dbf=0) |

**Table S4**. Parameter distributions for healthcare settings considered in case studies. The rehabilitation ward is largely characterized using data from a geriatric rehabilitation ward within a rehabilitation hospital in France (see Duval *et al*.) [(9)](https://sciwheel.com/work/citation?ids=8684589&pre=&suf=&sa=0&dbf=0). The geriatric ward is largely characterized using data from a short-stay geriatric unit within a tertiary hospital in France (see Vanhems *et al*.) [(10)](https://sciwheel.com/work/citation?ids=732780&pre=&suf=&sa=0&dbf=0). The paediatric ward is largely characterized using data from a general ward within a tertiary paediatric hospital in Italy (see Isella *et al*.) [(26)](https://sciwheel.com/work/citation?ids=732383&pre=&suf=&sa=0&dbf=0). For distributions: $\mathcal{N}\left( \mu,\sigma\right)$ indicates a Normal distribution with mean $\mu$ and standard deviation $\sigma$; $\mathcal{U}\left( a,b \right)$ indicates a Uniform distribution with minimum $a$ and maximum$b$. When parameter uncertainty was unavailable, normal distributions with standard deviation equal to 10% of the mean were assumed.

| **Parameter** | | | **Rehabilitation ward** | | **Geriatric ward** | | **Paediatric ward** | |
| --- | --- | --- | --- | --- | --- | --- | --- | --- |
| Symbol | Name | Unit | Distribution | Notes | Distribution | Notes | Distribution | Notes |
| $\mu$ | admission & discharge rate | day^-1^ | $\frac{1}{\mathcal{N}(49,4.9)}$ | [(9)](https://sciwheel.com/work/citation?ids=8684589&pre=&suf=&sa=0&dbf=0) | $\frac{1}{\mathcal{N}(2.5,0.25)}$ | [(10)](https://sciwheel.com/work/citation?ids=732780&pre=&suf=&sa=0&dbf=0) | $\frac{1}{\mathcal{N}(7,0.7)}$ | [(26)](https://sciwheel.com/work/citation?ids=732383&pre=&suf=&sa=0&dbf=0) |
| $N^{beds}$ | # of beds in the facility | / | $\mathcal{N}(30,3)$ | Average # of patients / week [(9)](https://sciwheel.com/work/citation?ids=8684589&pre=&suf=&sa=0&dbf=0) | $\mathcal{N}(19,1.9)$ | [(10)](https://sciwheel.com/work/citation?ids=732780&pre=&suf=&sa=0&dbf=0) | $\mathcal{N}(37,3.7)$ | Average # of patients / week [(26)](https://sciwheel.com/work/citation?ids=732383&pre=&suf=&sa=0&dbf=0) |
| $N^{hcw}$ | # of HCWs working/day | / | $\mathcal{N}(20,2)$ | Average # of HCWs / week [(9)](https://sciwheel.com/work/citation?ids=8684589&pre=&suf=&sa=0&dbf=0) | $\mathcal{N}(38,3.8)$ | 27 nurses + 11 doctors over a 4-day study period [(10)](https://sciwheel.com/work/citation?ids=732780&pre=&suf=&sa=0&dbf=0) | $\mathcal{N}(36,3.6)$ | 6 assistants + 20 nurses + 10 physicians present on a typical workday [(26)](https://sciwheel.com/work/citation?ids=732383&pre=&suf=&sa=0&dbf=0) |
| $\kappa^{pat\to pat}$ | # of contacts (patient-to-patient) | day^-1^ | $\mathcal{N}(7.1,0.71)$ | [(9)](https://sciwheel.com/work/citation?ids=8684589&pre=&suf=&sa=0&dbf=0) | $\mathcal{N}(1,0.1)$ | [(10)](https://sciwheel.com/work/citation?ids=732780&pre=&suf=&sa=0&dbf=0) | $\mathcal{N}(0.1,0.01)$ | [(26)](https://sciwheel.com/work/citation?ids=732383&pre=&suf=&sa=0&dbf=0) |
| $\kappa^{hcw\to hcw}$ | # of contacts (HCW-to-HCW) | day^-1^ | $\mathcal{N}(4.9,0.49)$ | [(9)](https://sciwheel.com/work/citation?ids=8684589&pre=&suf=&sa=0&dbf=0) | $\mathcal{N}(62,6.2)$ | [(10)](https://sciwheel.com/work/citation?ids=732780&pre=&suf=&sa=0&dbf=0) | $\mathcal{N}(34,3.4)$ | [(26)](https://sciwheel.com/work/citation?ids=732383&pre=&suf=&sa=0&dbf=0) |
| $\kappa^{pat\to hcw}$ | # of contacts (patient-to-HCW) | day^-1^ | $\mathcal{N}(3.9,0.39)$ | [(9)](https://sciwheel.com/work/citation?ids=8684589&pre=&suf=&sa=0&dbf=0) | $\mathcal{N}(30,3)$ | [(10)](https://sciwheel.com/work/citation?ids=732780&pre=&suf=&sa=0&dbf=0) | $\mathcal{N}(1.3,0.13)$ | [(26)](https://sciwheel.com/work/citation?ids=732383&pre=&suf=&sa=0&dbf=0) |
| $\rho$ | contact intimacy scaling coefficient | / | $\mathcal{U}(1,3)$ | Assumed | $\mathcal{U}(2,4)$ | Assumed | $\mathcal{U}(3,5)$ | Assumed |
| $H_{base}$ | hand hygiene compliance | % | $\mathcal{N}(12,1.2)$ | Pre-intervention, pre-pandemic compliance in Dutch nursing homes [(39)](https://sciwheel.com/work/citation?ids=9887763&pre=&suf=&sa=0&dbf=0) | $\mathcal{N}(46,4.6)$ | Pre-pandemic compliance in USA acute care hospitals [(40)](https://sciwheel.com/work/citation?ids=9529501&pre=&suf=&sa=0&dbf=0) | $\mathcal{N}(46,4.6)$ | Pre-pandemic compliance in USA acute care hospitals [(40)](https://sciwheel.com/work/citation?ids=9529501&pre=&suf=&sa=0&dbf=0) |
| $A_{base}$ | antibiotic exposure prevalence | % | $\mathcal{N}(4.9,0.49)$ | Proportion of LTCF residents in Europe receiving at least one antimicrobial [(41)](https://sciwheel.com/work/citation?ids=12683639&pre=&suf=&sa=0&dbf=0) | $\mathcal{N}(30.5,3.05)$ | Prevalence of antimicrobial exposure among European hospital inpatients [(42)](https://sciwheel.com/work/citation?ids=11600757&pre=&suf=&sa=0&dbf=0) | $\mathcal{N}(39.5,3.95)$ | Proportion of paediatric inpatients in Europe receiving at least one antibiotic [(43)](https://sciwheel.com/work/citation?ids=7134488&pre=&suf=&sa=0&dbf=0) |

**Table S5**. Parameter distributions for COVID-19 response scenarios considered in case studies. For distributions: $\mathcal{N}\left( \mu,\sigma\right)$ indicates a Normal distribution with mean $\mu$ and standard deviation $\sigma$; $\mathcal{U}\left( a,b \right)$ indicates a Uniform distribution with minimum $a$ and maximum$b$. Only policy responses ($\tau_{cp}$, $\tau_{um}$, $\tau_{hh}, \tau_{pl}$) varied across COVID-19 response scenarios; caseload responses were assumed identical across scenarios.

| **Parameter** | | **COVID-19 response scenario** | | | | | |
| --- | --- | --- | --- | --- | --- | --- | --- |
|  |  | **Organized** | | **Intermediate** | | **Overwhelmed** | |
| Symbol | Name | Distribution | Notes | Distribution | Notes | Distribution | Notes |
| $\tau_{as}$ | Abandoned stewardship | $0$ | Assumed | $0$ | Assumed | $0$ | Assumed |
| $\tau_{cp}$ | COVID-19 prophylaxis | $\mathcal{U}(0.047,0.152)$ | Share of COVID-19 cases with bacterial co-infection [(44)](https://sciwheel.com/work/citation?ids=10265615&pre=&suf=&sa=0&dbf=0) | $\mathcal{U}(0.68,0.81)$ | Share of adult COVID-19 patients receiving antibiotic prophylaxis [(44)](https://sciwheel.com/work/citation?ids=10265615&pre=&suf=&sa=0&dbf=0) | $\mathcal{U}(0.9,1)$ | Assumed prescribing to nearly all symptomatic COVID-19 patients |
| $\tau_{cd}$ | Cohorting dis-organization | $\mathcal{U}(0.4,0.6)$ | Assumed | $\mathcal{U}(0.4,0.6)$ | Assumed | $\mathcal{U}(0.4,0.6)$ | Assumed |
| $\tau_{pl}$ | Patient lockdown | $\mathcal{U}(0.8,1)$ | Assumed highly effective lockdown | $\mathcal{U}(0.4,0.6)$ | Assumed moderately effective lockdown | $\mathcal{U}(0,0.2)$ | Assumed minimally effective lockdown |
| $\tau_{um}$ | Universal masking | $\mathcal{N}(0.99,0.003)\times\mathcal{U}(0.5,1)$ | N95 respirator efficacy (99%, SD 0.3%) [(45)](https://sciwheel.com/work/citation?ids=11275956&pre=&suf=&sa=0&dbf=0) multiplied by assumed compliance | $\mathcal{N}(0.59,0.069)\times\mathcal{U}(0.5,1)$ | Medical mask efficacy (59%, SD 6.9%) [(45)](https://sciwheel.com/work/citation?ids=11275956&pre=&suf=&sa=0&dbf=0) multiplied by assumed compliance | $0$ | No masking |
| $\tau_{hh}$ | Hand hygiene | $\mathcal{U}(0.25,0.3)$ | Increase in compliance from 12% to 36% across 36 Dutch nursing homes in interventional, pre-pandemic context [(39)](https://sciwheel.com/work/citation?ids=9887763&pre=&suf=&sa=0&dbf=0) | $\mathcal{U}(0.15,0.2)$ | Increase in compliance from 46% to 56% during the first wave of COVID-19 across 84 units in 9 USA hospitals [(40)](https://sciwheel.com/work/citation?ids=9529501&pre=&suf=&sa=0&dbf=0) | $\mathcal{U}(0,0.05)$ | Assumed marginal improvement |
| $\tau_{cs}$ | COVID-19 stays | $\mathcal{U}(0.8,1)$ | Assumed | $\mathcal{U}(0.8,1)$ | Assumed | $\mathcal{U}(0.8,1)$ | Assumed |
| $\tau_{ss}$ | Staff sick leave | $\mathcal{U}(0,0.5)$ | Assumed | $\mathcal{U}(0,0.5)$ | Assumed | $\mathcal{U}(0,0.5)$ | Assumed |
| $\tau_{ra}$ | Reduced admission | $\mathcal{U}(0.4,0.6)$ | Assumed | $\mathcal{U}(0.4,0.6)$ | Assumed | $\mathcal{U}(0.4,0.6)$ | Assumed |
| $\tau_{sc}$ | Sicker casemix | $\mathcal{U}(0.4,0.6)$ | Assumed | $\mathcal{U}(0.4,0.6)$ | Assumed | $\mathcal{U}(0.4,0.6)$ | Assumed |

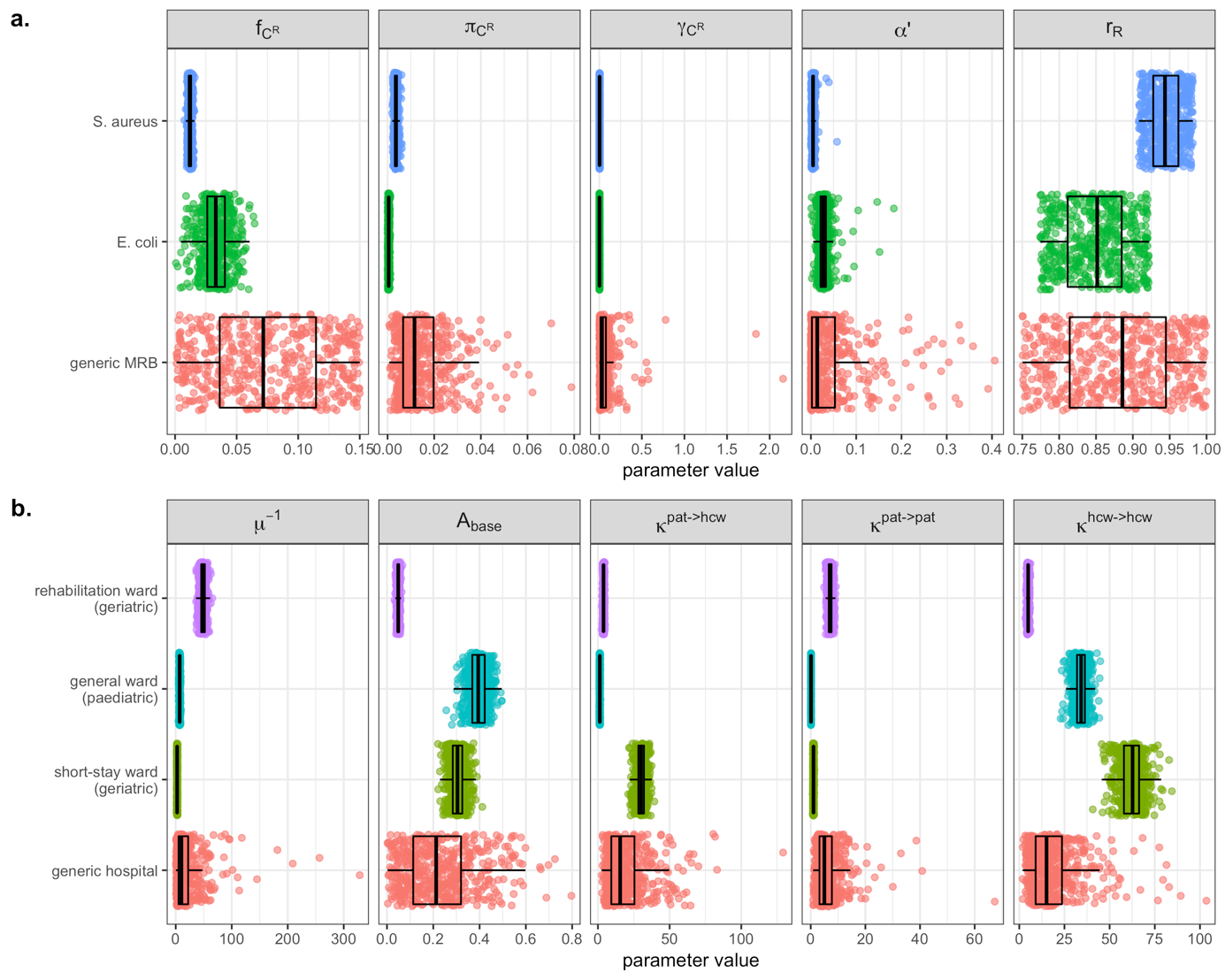

**Figure S3.** Comparison of parameter values generated during Monte Carlo simulations for simulation set 1 (generic MRB in generic hospitals) and simulation set 2 (case studies). A total $n=500$ values of each parameter were generated for each bacterium and each facility. (**a**) Selected bacterial parameters for the focal antibiotic-resistant strain, including the admission fraction ($f_{C^{R}}$), the transmission rate per contact ($\pi_{B^{R}}$), the natural clearance rate ($\gamma_{B^{R}}$), the endogenous acquisition rate ($\alpha'$) and the antibiotic resistance level ($r_{R}$). (**b**) Selected healthcare facility parameters, including the average patient length of stay ($\mu^{-1}$), baseline antibiotic exposure prevalence ($A_{base}$), and daily rates of patient-HCW contact ($\kappa^{pat\to hcw}$), patient-patient contact ($\kappa^{pat\to pat}$) and HCW-HCW contact ($\kappa^{hcw\to hcw}$). Corresponding probability laws are defined in **Tables S2 – S4**.

***1.4 Epidemiological indicators***

*Prevalence*

Daily prevalence of SARS-CoV-2 infection is given by,

$$V^{k}\left( t \right)=\sum_{g\in h} I_{g}^{k}\left( t \right)$$

daily prevalence of colonization with strain $B^{b}$ is given by

$$C^{b}\left( t \right)=\sum_{X\in Y} X_{b}^{pat}\left( t \right)$$

and daily prevalence of HCW carriage of strain $B^{b}$is given by

$$T^{b}(t)=\sum_{X\in Y} X_{b}^{hcw}\left( t \right)$$

*Incidence*

Daily incidence of infection/colonization/carriage $\Lambda_{m}^{k}$ for each microorganism $m$ and host type $k$ from $t$ to $t+1$ is calculated from the integral of the corresponding force of infection,

$$\Lambda_{m}^{k}=\int_{t}^{t+1} q_{m}^{k}(t)\times\lambda_{m}^{k}(t,\tau_{cd},\tau_{pl},\tau_{um}) dt$$

where $q_{m}^{k}\left( t \right)$ represents: the number of patients susceptible to infection {$m=V,k=pat\}$; the number of HCWs susceptible to SARS-CoV-2 infection $\{m=V,k=hcw\}$; the number of patients uncolonized by either strain of the focal bacterium $\{m\in\left\{ B^{S}, B^{R} \right\},k=pat\}$; or the number of staff not carrying bacteria $\{m\in\left\{ B^{S}, B^{R} \right\},k=hcw\}$. Note that daily incidence of patient colonization with $B^{R}$ also accounts for within-host endogenous outgrowth subsequent to antibiotic exposure,

$$\Lambda_{B^{R}}^{pat}=\int_{t}^{t+1} q_{B^{R}}^{pat}\left( t \right)\times\left( \lambda_{B^{R}}^{pat}\left( t,\tau_{cd},\tau_{pl} \right)+\alpha(t,\tau_{as},\tau_{cp}) \right) dt$$

Cumulative incidence is calculated as the integral over simulation time (from $t_{0}=0$ to $t_{end}=180$), while the incidence rate is calculated for each $k$ as the cumulative incidence divided by the cumulative number of patient-days or HCW-days in the model, respectively, accounting for impacts of a changing population size as a result of COVID-19 responses.

*Patient-days colonized*

To capture average dynamics of bacterial colonization over time, the cumulative number of patient-days colonized with strain $B^{b}$ is calculated as

$$P^{b}=\int_{t_{0}}^{t_{end}} C^{b}(t) dt$$

*Resistance rate*

The resistance rate $R$ is used to express the relative share of patients colonized with drug-resistant as opposed to drug-sensitive bacteria. This is given for any time $t$ as

$$R\left( t \right)=\frac{C^{R}(t)}{C^{S}\left( t \right)+C^{R}(t)}$$

and over the full simulation period, the average resistance rate is given as

$$\bar{R}=\frac{P^{R}}{P^{S}+P^{R}}$$

Other secondary outcomes include: the peak incidence of SARS-CoV-2 infection, the peak incidence of bacterial colonization, the timing of peak infection/colonization incidence, the number of patients exposed to antibiotics, the daily number of patient contacts, the daily number of HCW contacts, the average delay between successful HCW decontamination, the HCW:patient ratio and the daily number of new patients already colonized with $B^{R}$ upon admission to the healthcare facility.

***1.5 Simulation convergence and sensitivity analysis***

*Bootstrapping outcomes*

Bootstrap resampling of epidemiological outcomes was used to determine that a sufficient number of Monte Carlo simulations were conducted for epidemiological outcomes to converge. This was evaluated for simulation set 1 by sampling $m\in\{5,10,\ldots,500\}$ values from ${\Delta\Gamma}^{n}$ and bootstrap resampling with 50 replicates, producing a distribution of mean ${\Delta\Gamma}^{m}$. Chosen indicators for bootstrap resampling were change in cumulative SARS-CoV-2 infection incidence ($\Delta\Lambda_{V}$, **Figure S4**) and change in cumulative MRB colonization incidence ($\Delta\Lambda_{B^{R}}$, **Figure S5**)

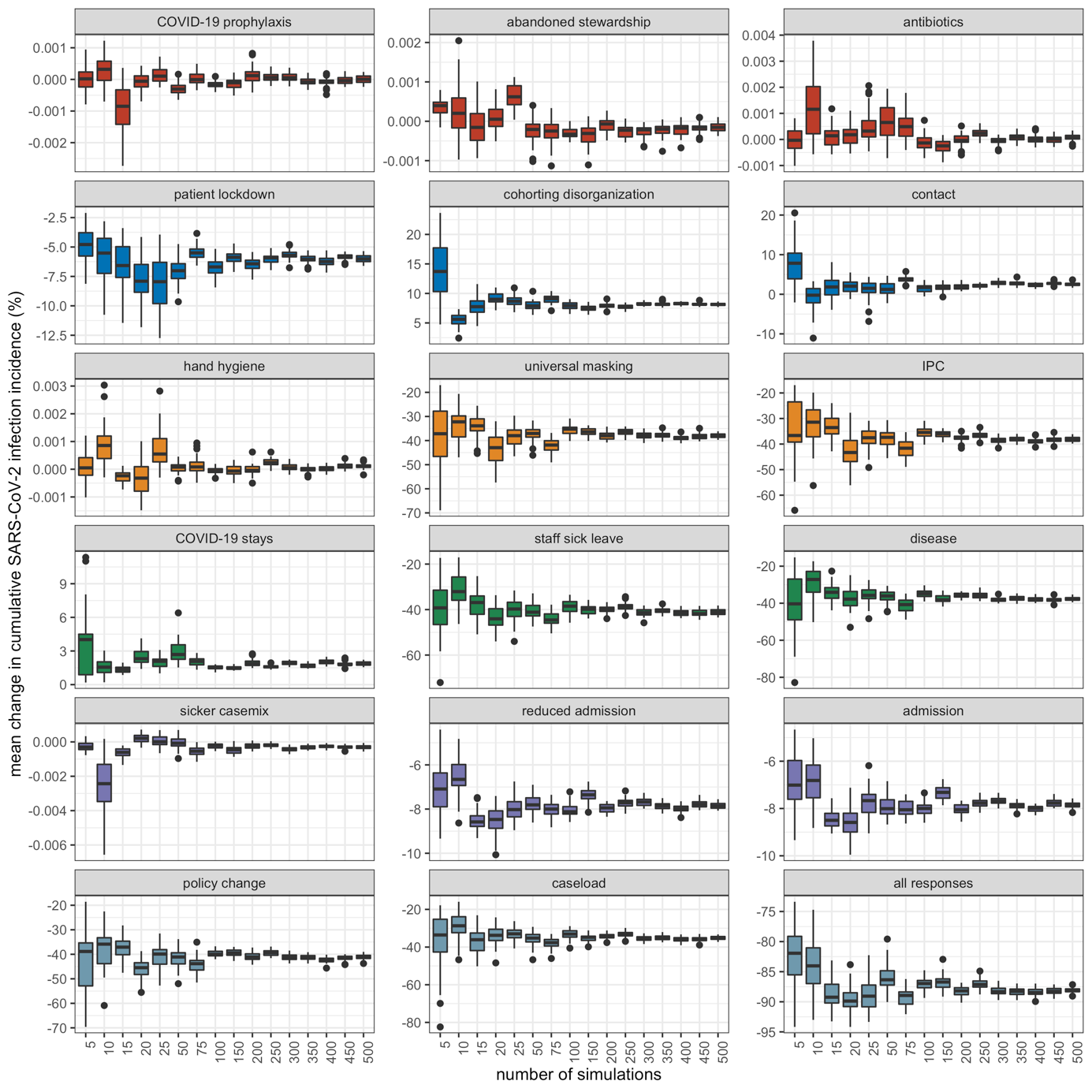

**Figure S4.** Distribution of mean change in cumulative SARS-CoV-2 incidence (y-axis) after bootstrap resampling with 50 replicates among $m$ random samples of simulation outputs (x-axis) for different COVID-19 responses and combinations of COVID-19 responses (panels).

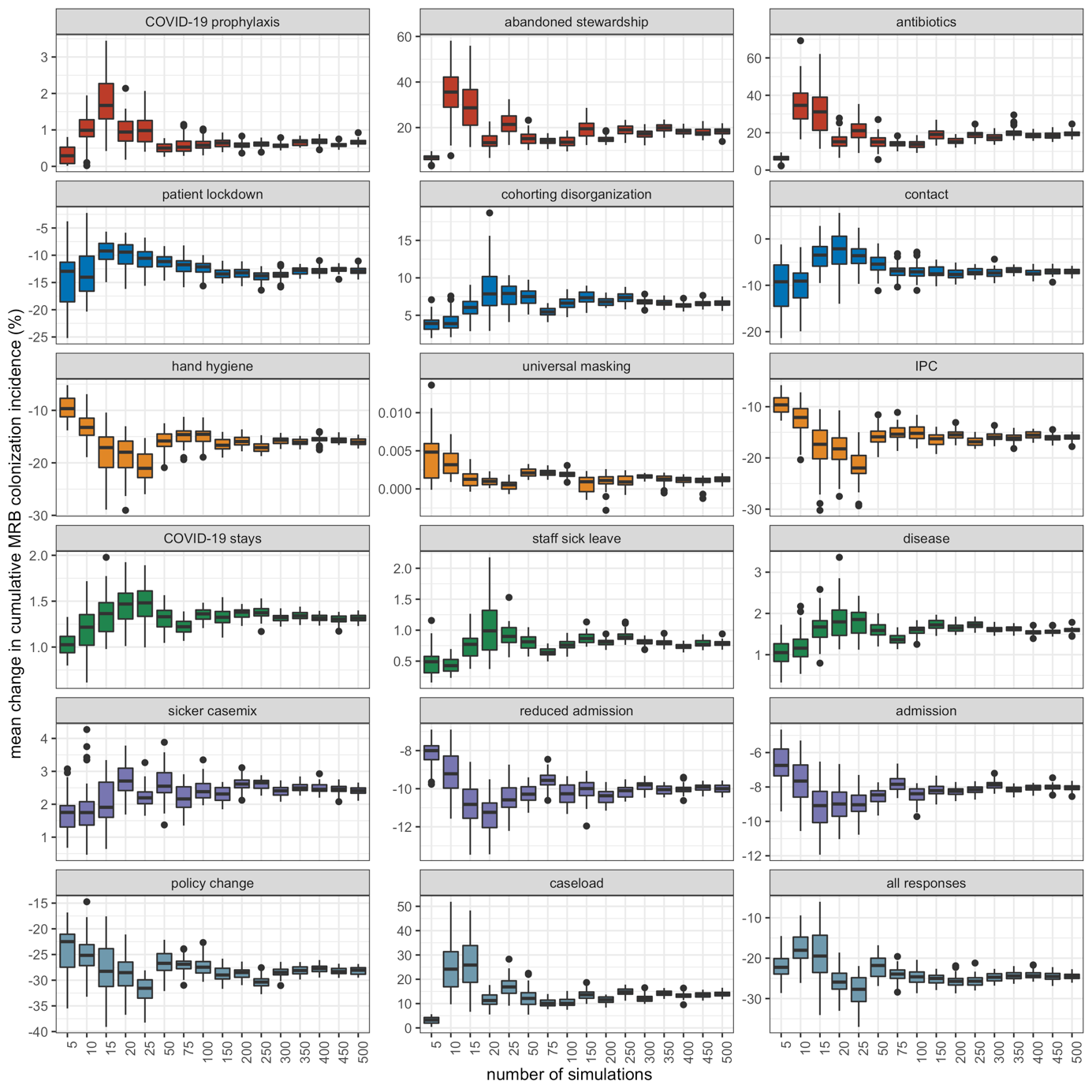

**Figure S5.** Distribution of mean change in cumulative patient MRB colonization incidence (y-axis) after bootstrap resampling with 50 replicates among $m$ random samples of simulation outputs (x-axis) for different COVID-19 responses and combinations of COVID-19 responses (panels).

*Partial rank correlation coefficients*

Multivariate sensitivity analyses were conducted to quantify impacts of parameter uncertainty on model outcomes in the context of the first simulation set (generic MRB in generic hospitals). Partial rank correlation coefficients (PRCCs) were calculated using the function epi.prcc from the R package epiR.[(46)](https://sciwheel.com/work/citation?ids=13307238&pre=&suf=&sa=0&dbf=0) PRCCs were calculated between parameter values varied in these simulations ($\Theta_{B}^{n}, \Theta_{F}^{n}$) and epidemiological indicators $\Gamma^{n}$ for simulations in which no COVID-19 responses were included (all $\tau=0$; **Supplementary Figure S6**) and those in which all responses were included (all $\tau=0.5$; **Supplementary Figure S7**).

Cumulative MRB colonization incidence was most positively associated (highest PRCCs) with rates of endogenous acquisition ($\alpha'$) and bacterial transmission ($\pi_{B}$), baseline antibiotic exposure prevalence ($A_{base}$), the number of beds in the facility ($N^{beds}$), and the share of patients already colonized upon admission ($f_{C^{R}}$). Conversely, the average resistance rate was unassociated with $\pi_{B}$, but also highly positively associated with $\alpha$’, $A_{base}$, and $f_{C^{R}}$, in addition to being highly positively associated with the effective colony kill rate ($\theta$). These parameters emerged as having the greatest PRCCs for each respective outcome in simulations both with and without COVID-19 responses.

Cumulative MRB colonization incidence was most negatively associated with hand hygiene compliance ($H_{base}$), the proportion of patients already colonized upon admission with competing antibiotic-sensitive bacteria ($f_{C^{S}}$), and the natural bacterial clearance rate ($\gamma$). Conversely, the average resistance rate was comparatively unassociated with $H_{base}$, but also highly negatively associated with $f_{C^{S}}$ and $\gamma$, in addition to being highly negatively associated with the patient admission and discharge rate ($\mu$). These parameters emerged as having the most negative PRCCs for each respective outcome in simulations both with and without COVID-19 responses.

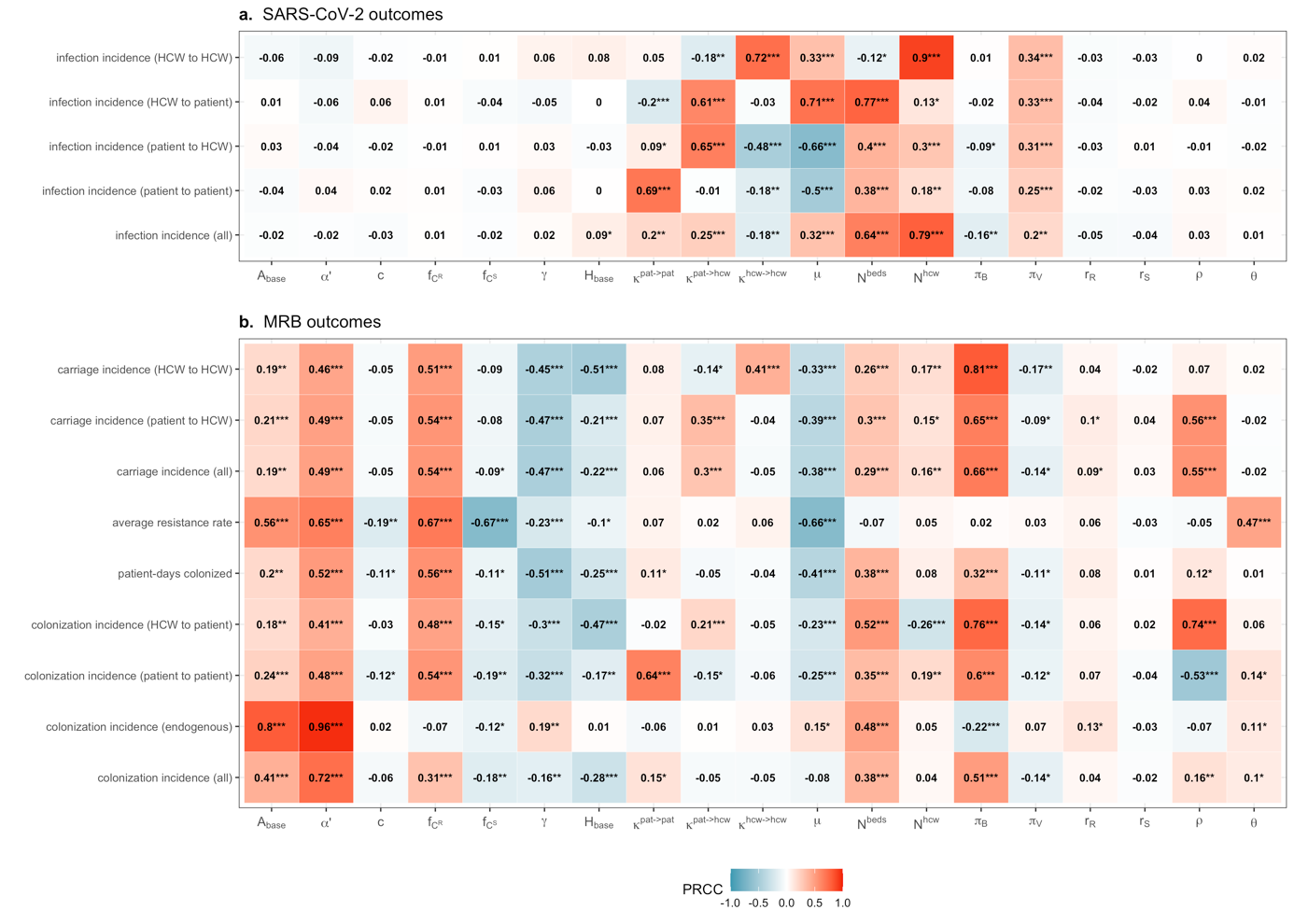

**Figure S6.** Partial rank correlation coefficients (PRCCs) between model parameters (x-axis) and epidemiological indicators (y-axis) from Monte Carlo simulations for generic MRB and generic hospitals (simulation set 1), here in the absence of COVID-19 responses ($\tau=0$). Asterisks denote p-value thresholds ($*=p<0.05; **=p<5\times{10}^{-4}; ***=p<5\times{10}^{-6}$).

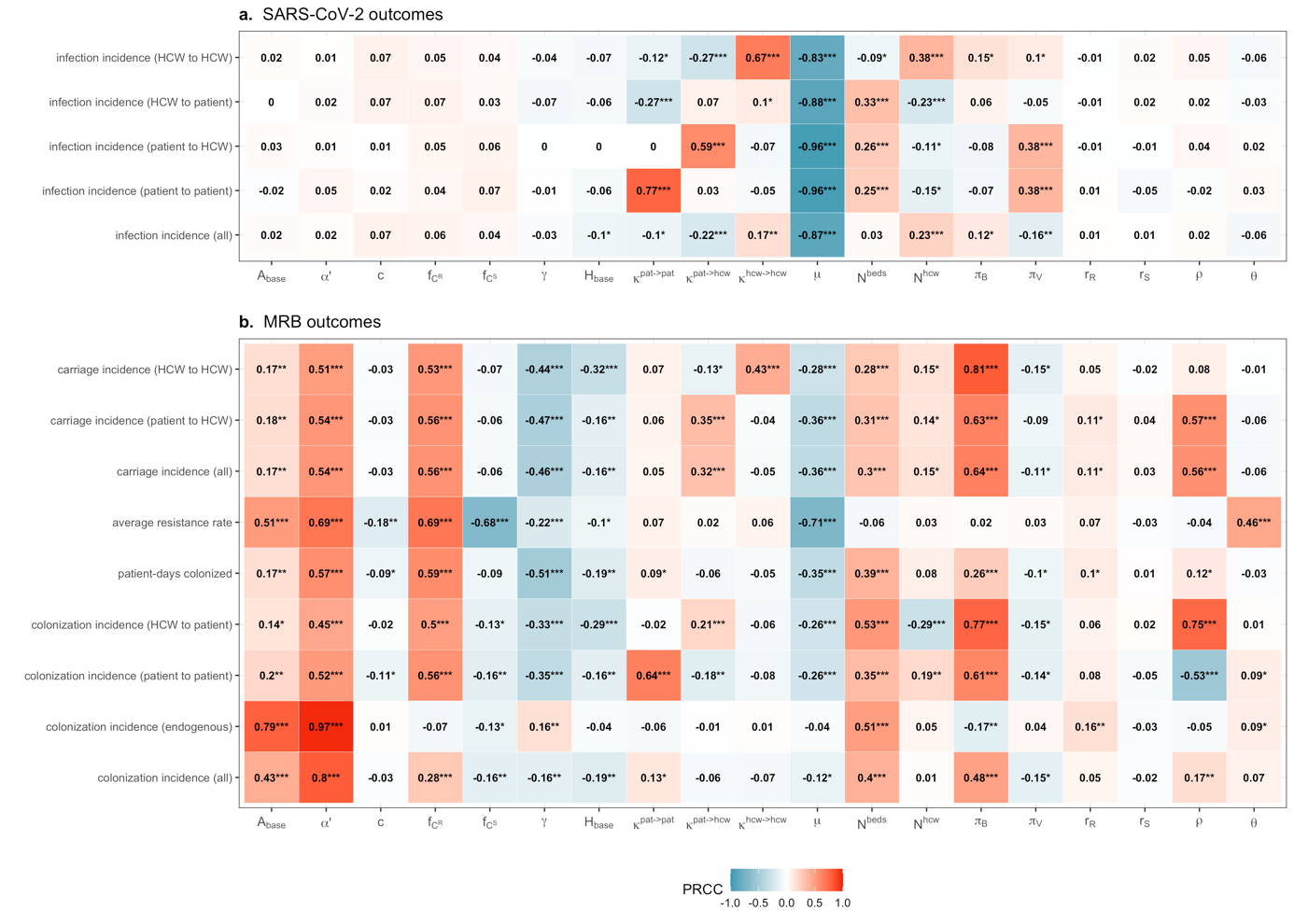

**Figure S7.** Partial rank correlation coefficients (PRCCs) between model parameters (x-axis) and epidemiological indicators (y-axis) from Monte Carlo simulations for generic MRB and generic hospitals (simulation set 1), here including all COVID-19 responses simultaneously ($\tau=0.5$). Asterisks denote p-value thresholds ($*=p<0.05; **=p<5\times{10}^{-4}; ***=p<5\times{10}^{-6}$).

***1.6 Supplementary results: Generic MRB in generic hospitals***

Generic hospitals were characterized by substantial heterogeneity, with a median 569 (95% UI: 124–972) patient beds, 1.65 (0.32–2.95) HCWs per patient, 94 (8–413) patients exposed daily to antibiotics, 22 (8–67) daily contacts per patient, 30 (10–96) daily contacts per HCW, 2.60 (0.09–41.41) daily admissions of patients already colonized with resistant bacteria, and 10 day (1–87) patient length of stay (**Figure S8**). In these hospitals, the epidemiological dynamics of generic MRB were also heterogeneous, with a median patient colonization prevalence of 11.0% (0.8%–61.0%), resistance rate of 47.1% (6.4%–95.9%), and similar average contributions of each colonization acquisition route, including a median 0.75 (0.03–8.12) daily patient colonization events resulting from transmission from other patients, 0.97 (0.02–20.54) from transmission from HCWs, and 0.75 (<0.01–17.75) from endogenous acquisition (**Figure S9, panels a-e**). In these hospitals in the absence of COVID-19 responses ($\tau=0$), SARS-CoV-2 introductions led to explosive outbreaks, with a median 19.2% (4.9%–24.9%) peak infection prevalence, 41-day (27–87) delay to peak infection incidence, and 1,970 (370–4,909) cumulative SARS-CoV-2 infections over 180 days, with HCW-to-patient transmission dominating relative to other routes (**Figure S9, panels f-h**).

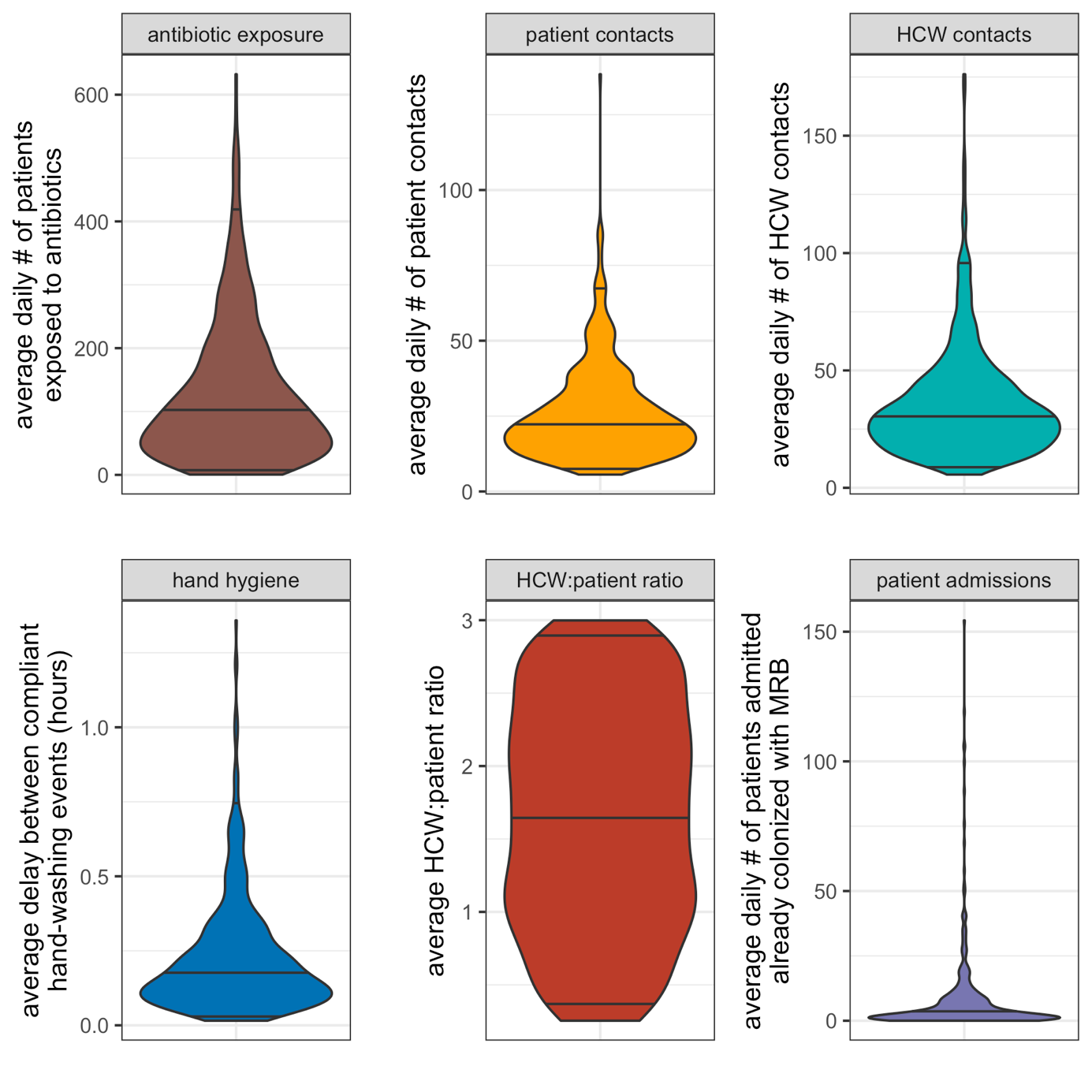

**Figure S8.** Baseline indicators for healthcare-associated behaviours and hospital demography at endemic equilibrium from simulation set 1 (i.e. in the absence of COVID-19 responses, all $\tau=0$). (**a**) The average daily number of patients exposed to antibiotics ($A_{base}\times N^{pat}$). (**b**) The average daily number of patient contacts with other individuals ($\kappa^{pat\to pat}+\kappa^{pat\to hcw}$). (**c**) The average daily number of HCW contacts with other individuals ($\kappa^{hcw\to hcw}+\kappa^{hcw\to pat}$). (**d**) The average delay between compliant handwashing events ($\omega/\mathrm{day}^{-1}\times24 hours/day$). (**e**) The average HCW:patient ratio (${N^{hcw}}/{N^{pat}}$). (**f**) The average daily number of patients admitted colonized with MRB ($\mu^{adm}\times N^{beds}\times f_{C^{R}}$).

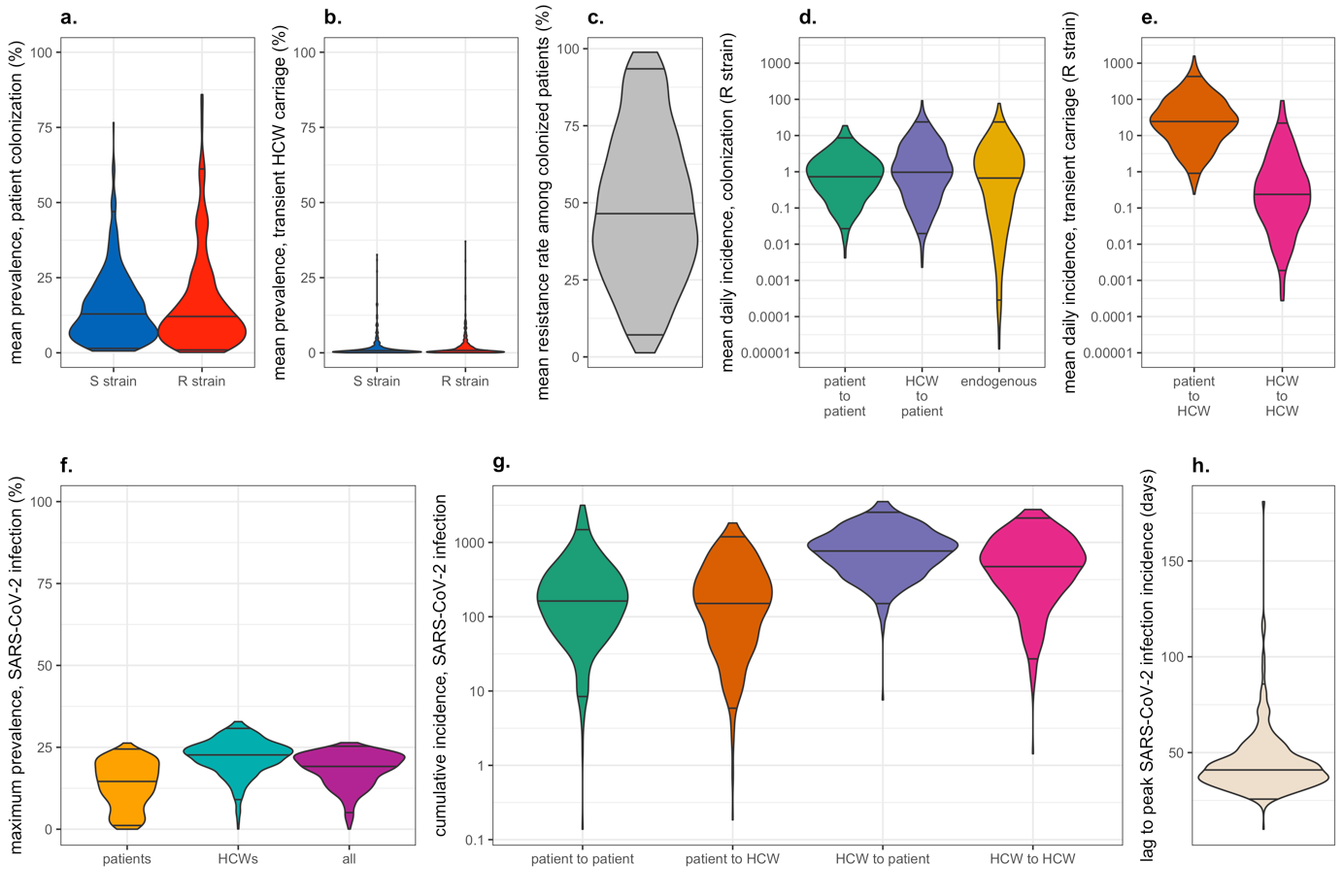

**Figure S9.** Baseline epidemiological indicators for generic MRB in generic hospitals in the absence of COVID-19 responses (all $\tau=0$), therefore assuming complete independence between bacterial colonization and SARS-CoV-2 infection. Bacterial indicators represent daily dynamics at endemic equilibrium, and SARS-CoV-2 indicators represent indicators calculated cumulatively from outbreaks simulated over 180 days. Here, in lieu of MRB, *R strain* is used to differentiate the focal antibiotic-resistant bacteria from the competing antibiotic-sensitive *S strain*. Parameter distributions underlying indicator uncertainty are defined in **Table S2**. (**a**) Equilibrium prevalence of patient colonization with antibiotic-sensitive bacteria (S strain) and antibiotic-resistant bacteria (R strain). (**b**) Equilibrium prevalence of HCW carriage of antibiotic-sensitive bacteria (S strain) and antibiotic-resistant bacteria (R strain). (**c**) The equilibrium bacterial resistance rate among colonized patients. (**d**) Equilibrium daily patient colonization incidence for the R strain, stratified by route of acquisition. (**e**) Equilibrium daily HCW carriage incidence for the R strain, stratified by route of acquisition. (**f**) Maximum prevalence of SARS-CoV-2 infection over 180 days, stratified by type of individual. (**g**) Cumulative incidence of nosocomial SARS-CoV-2 infection over 180 days, stratified by route of acquisition. (**i**) The time to peak SARS-CoV-2 infection incidence, relative to the introduction of the two index cases into the healthcare facility at $t=0$.

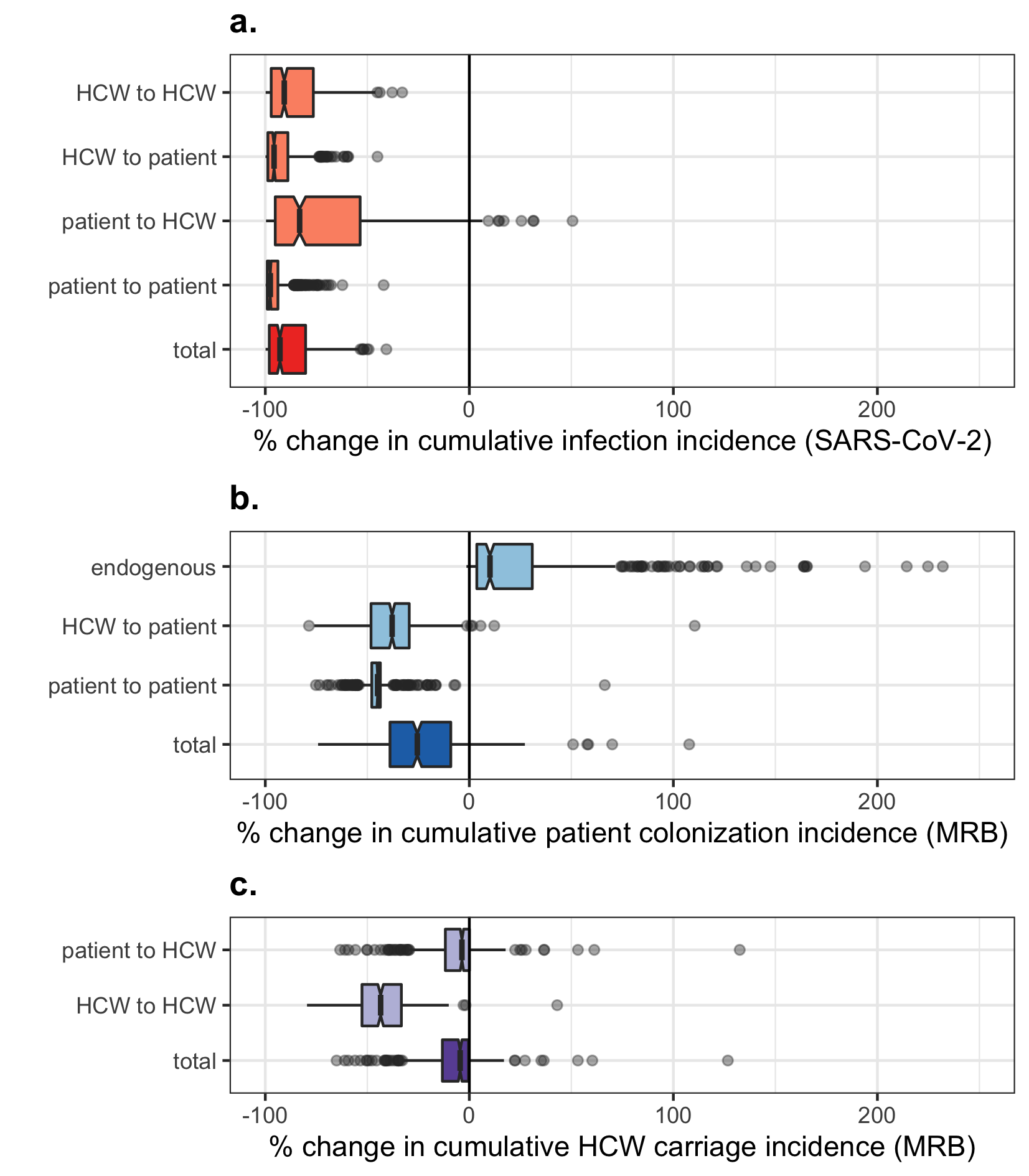

**Figure S10.** Impact of combined COVID-19 responses on cumulative nosocomial incidence of (**a**) patient and HCW SARS-CoV-2 infection, (**b**) patient MRB colonization incidence, and (**c**) HCW MRB transient carriage incidence. Each data point represents one of $n=500$ unique MRB-hospital pairs. For each indicator (row), change due to COVID-19 responses is calculated as the difference across matched simulations, i.e. those including all COVID-19 responses simultaneously ($\tau=0.5$) versus those including no COVID-19 responses ($\tau=0$). Indicators are calculated cumulatively over $t=180$ days of simulation, after introduction of two index cases of SARS-CoV-2 into the hospital at $t=0$. Boxplots represent the $IQR$; whiskers extend to the furthest value up to $\pm1.5\times IQR$; and notches extend $1.58\times IQR/\sqrt{n}$, giving an approximate 95% confidence interval for comparing medians. Scales are pseudo-log_10_-transformed using an inverse hyperbolic sine function (R package ggallin).

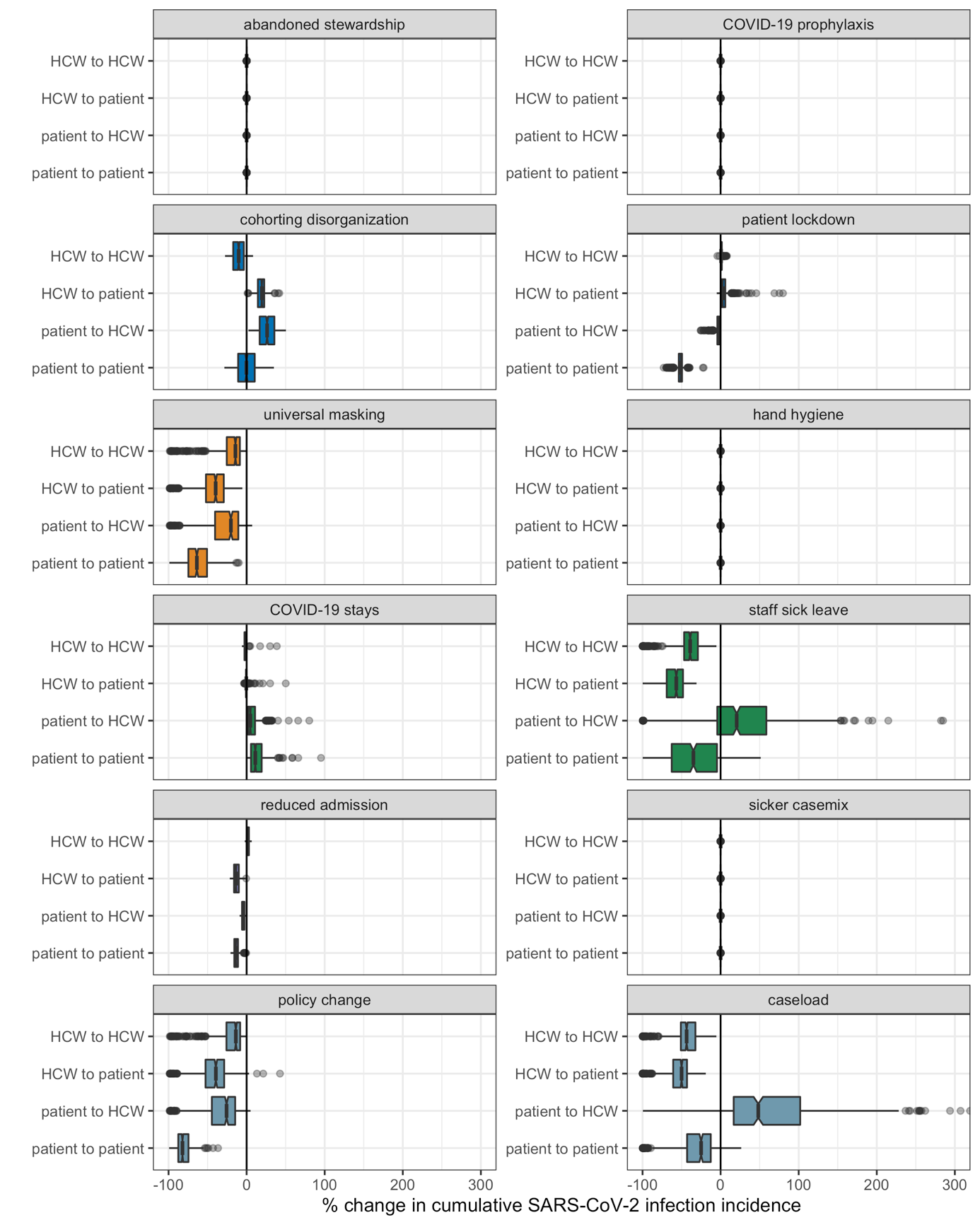

**Figure S11**. Impacts of COVID-19 responses on nosocomial acquisition of SARS-CoV-2, stratified by route of acquisition (y-axis) across different COVID-19 responses and combinations of COVID-19 responses (panels). Each data point represents one of $n=500$ unique MRB-hospital pairs. For each indicator (row), change due to COVID-19 responses is calculated as the difference across matched simulations, i.e. those including the given COVID-19 response(s) ($\tau=0.5$) versus those including no COVID-19 responses ($\tau=0$). Indicators are calculated cumulatively over $t=180$ days of simulation, after introduction of two index cases of SARS-CoV-2 into the hospital at $t=0$. Boxplots represent the $IQR$; whiskers extend to the furthest value up to $\pm1.5\times IQR$; and notches extend $1.58\times IQR/\sqrt{n}$, giving an approximate 95% confidence interval for comparing medians.

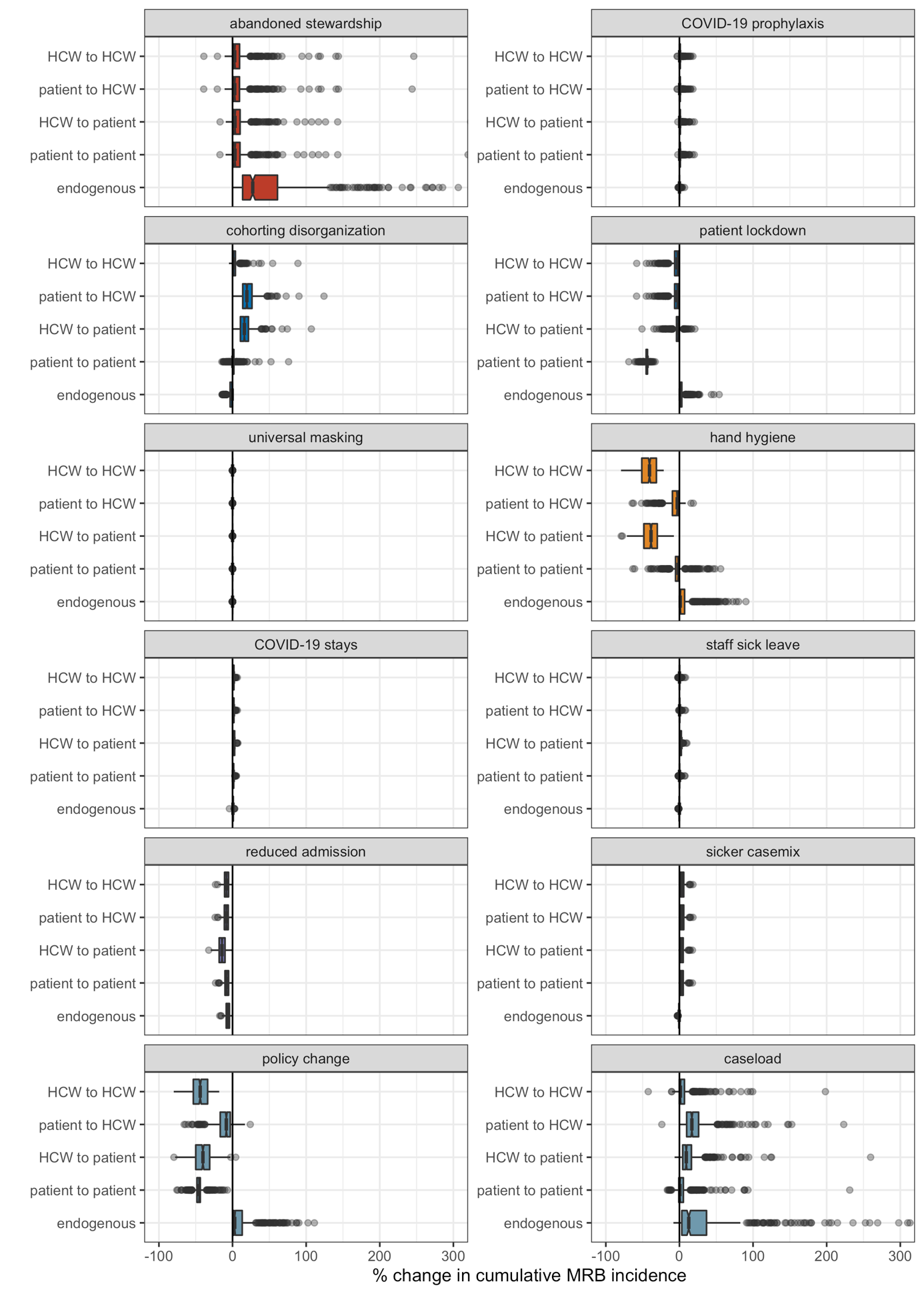

**Figure S12**. Impacts of COVID-19 responses on nosocomial acquisition of generic multidrug-resistant bacteria (MRB), stratified by route of acquisition (y-axis) across different COVID-19 responses and combinations of COVID-19 responses (panels). Each data point represents one of $n=500$ unique MRB-hospital pairs. For each indicator (row), change due to COVID-19 responses is calculated as the difference across matched simulations, i.e. those including the given COVID-19 response(s) ($\tau=0.5$) versus those including no COVID-19 responses ($\tau=0$). Indicators are calculated cumulatively over $t=180$ days of simulation, after introduction of two index cases of SARS-CoV-2 into the hospital at $t=0$. Boxplots represent the $IQR$; whiskers extend to the furthest value up to $\pm1.5\times IQR$; and notches extend $1.58\times IQR/\sqrt{n}$, giving an approximate 95% confidence interval for comparing medians.

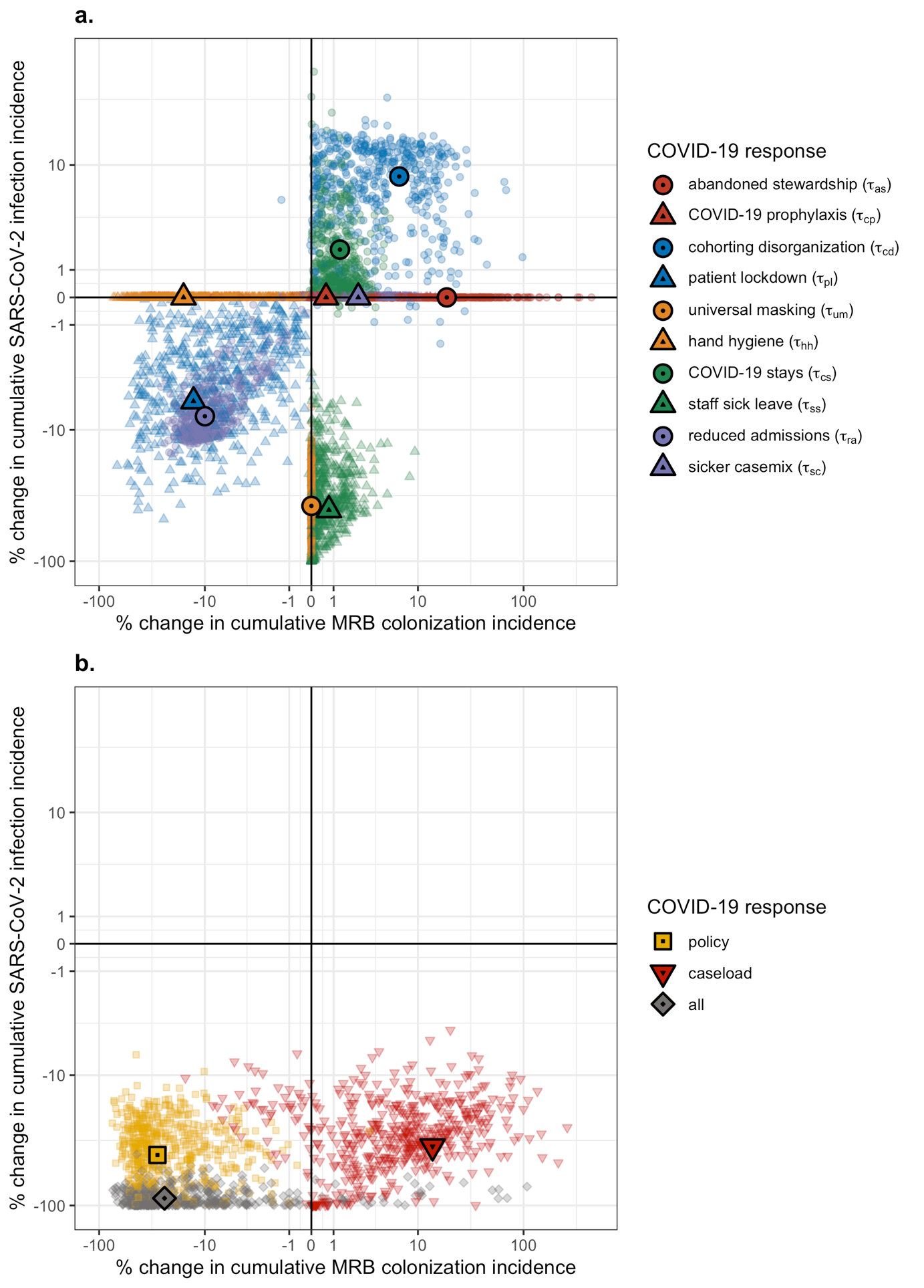

**Figure S13.** Change in cumulative MRB colonization incidence (x-axis) and cumulative SARS-CoV-2 infection incidence (y-axis) resulting from: (**a**) individual COVID-19 responses, each given by a unique colour-shape combination, and (**b**) combinations of COVID-19 responses. Small translucent points represents unique MRB-hospital pairs, and larger opaque points represent means across $n=500$ pairs. For each indicator, change due to COVID-19 responses is calculated as the difference between matched simulations, i.e. those including the respective COVID-19 response(s) ($\tau=0.5$) versus those including no COVID-19 responses ($\tau=0$). Indicators are calculated cumulatively over $t=180$ days of simulation, after introduction of two index cases of SARS-CoV-2 into the hospital at $t=0$. Scales are pseudo-log_10_-transformed using an inverse hyperbolic sine function (R package ggallin).

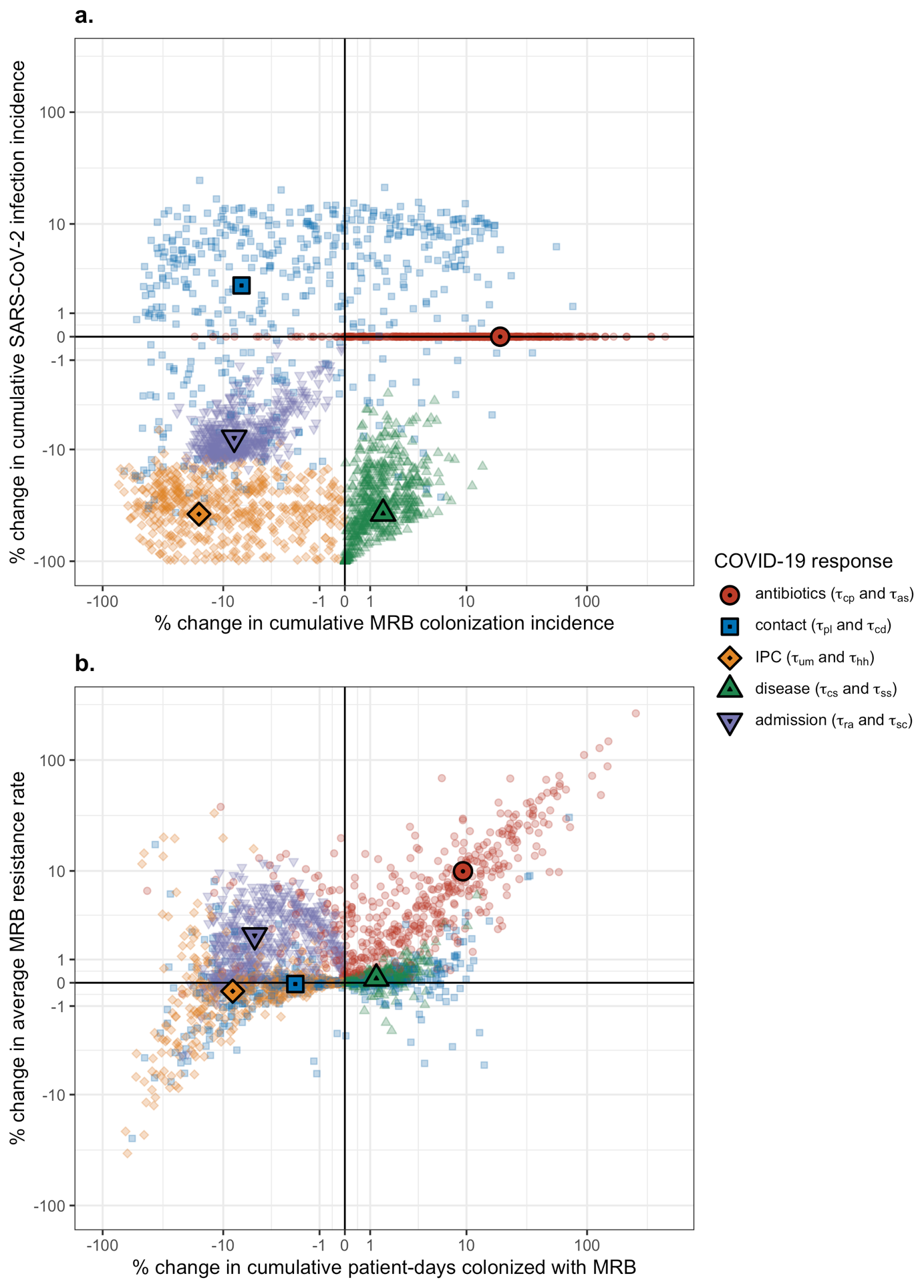

**Figure S14.** Change in epidemiological indicators resulting from pairwise combinations of categories of COVID-19 responses (antibiotics, contact, IPC, disease, admission), each given by a unique colour-shape combination. (**a**) Change in cumulative MRB colonization incidence (x-axis) and cumulative SARS-CoV-2 infection incidence (y-axis). (**b**) Change in the cumulative number of patient-days of MRB colonization (x-axis) and the average MRB resistance rate (y-axis). Small translucent points represents unique MRB-hospital pairs, and larger opaque points represent means across $n=500$ pairs. For each indicator, change due to COVID-19 responses is calculated as the difference between matched simulations, i.e. those including the respective COVID-19 responses ($\tau=0.5$) versus those including no COVID-19 responses ($\tau=0$). Indicators are calculated cumulatively over $t=180$ days of simulation, after introduction of two index cases of SARS-CoV-2 into the hospital at $t=0$. Scales are pseudo-log_10_-transformed using an inverse hyperbolic sine function (R package ggallin).

***1.7. Supplementary results: Case studies***

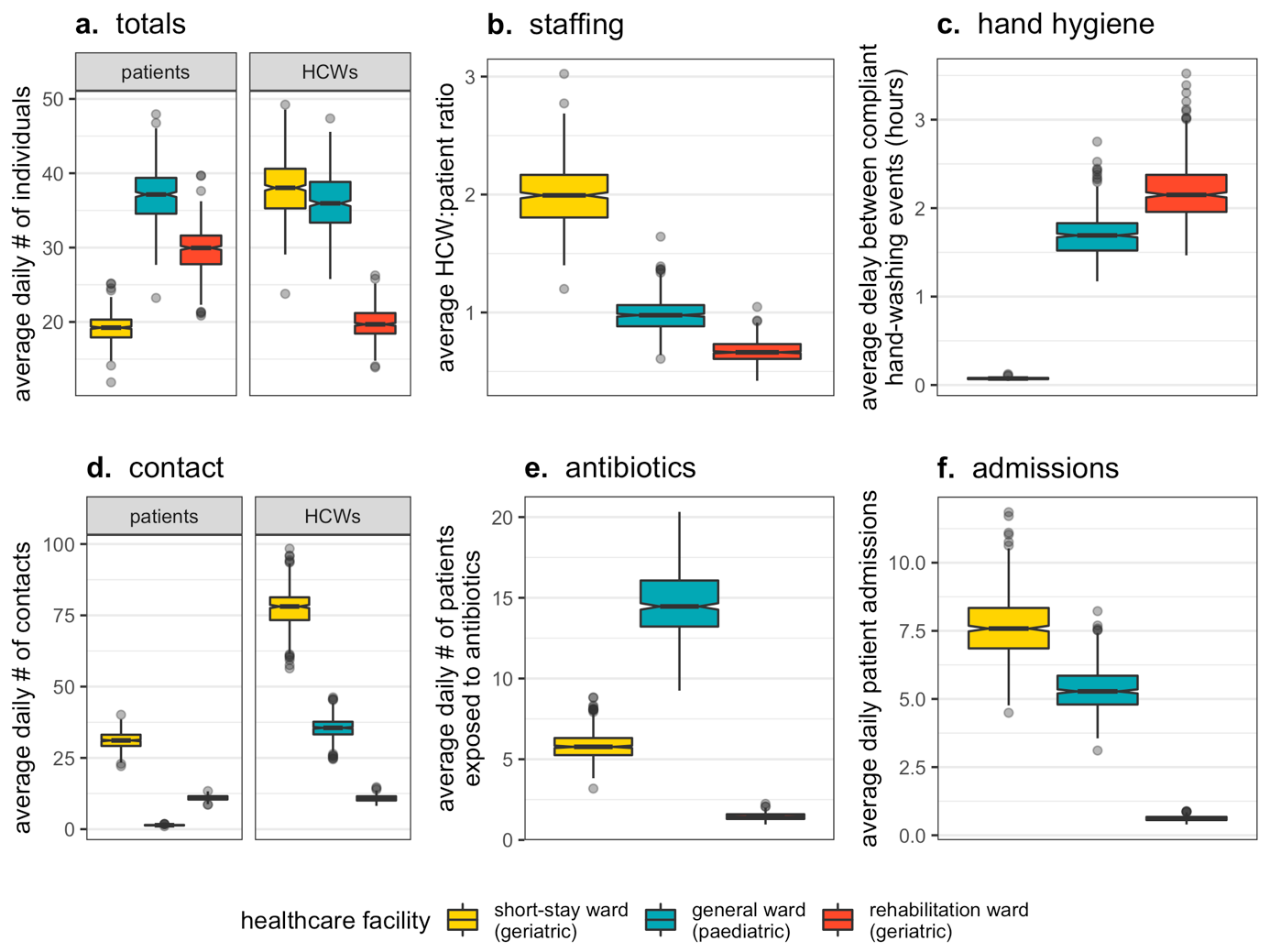

**Figure S15.** Baseline indicators for healthcare-associated behaviours and hospital demography at endemic equilibrium in each ward (colours). Indicators represent results from simulations excluding COVID-19 responses (all $\tau=0$). Each data point represents one of $n=500$ Monte Carlo simulations. Boxplots represent the $IQR$; whiskers extend to the furthest value up to $\pm1.5\times IQR$; and notches extend $1.58\times IQR/\sqrt{n}$, giving an approximate 95% confidence interval for comparing medians. (**a**) The average daily number of patients ($N^{pat}$) and HCWs ($N^{hcw}$) in the ward. (**b**) The average HCW:patient ratio (${N^{hcw}}/{N^{pat}}$). (**c**) The average delay between compliant handwashing events ($\omega/\mathrm{day}^{-1}\times24 hours/day$). (**d**) The average daily number of patient contacts with other individuals ($\kappa^{pat\to pat}+\kappa^{pat\to hcw}$) and HCW contacts with other individuals ($\kappa^{hcw\to hcw}+\kappa^{hcw\to pat}$). (**e**) The average daily number of patients exposed to antibiotics ($A_{base}\times N^{pat}$). (**f**) The average daily number of patients admitted to the ward ($\mu\times N^{pat}$).

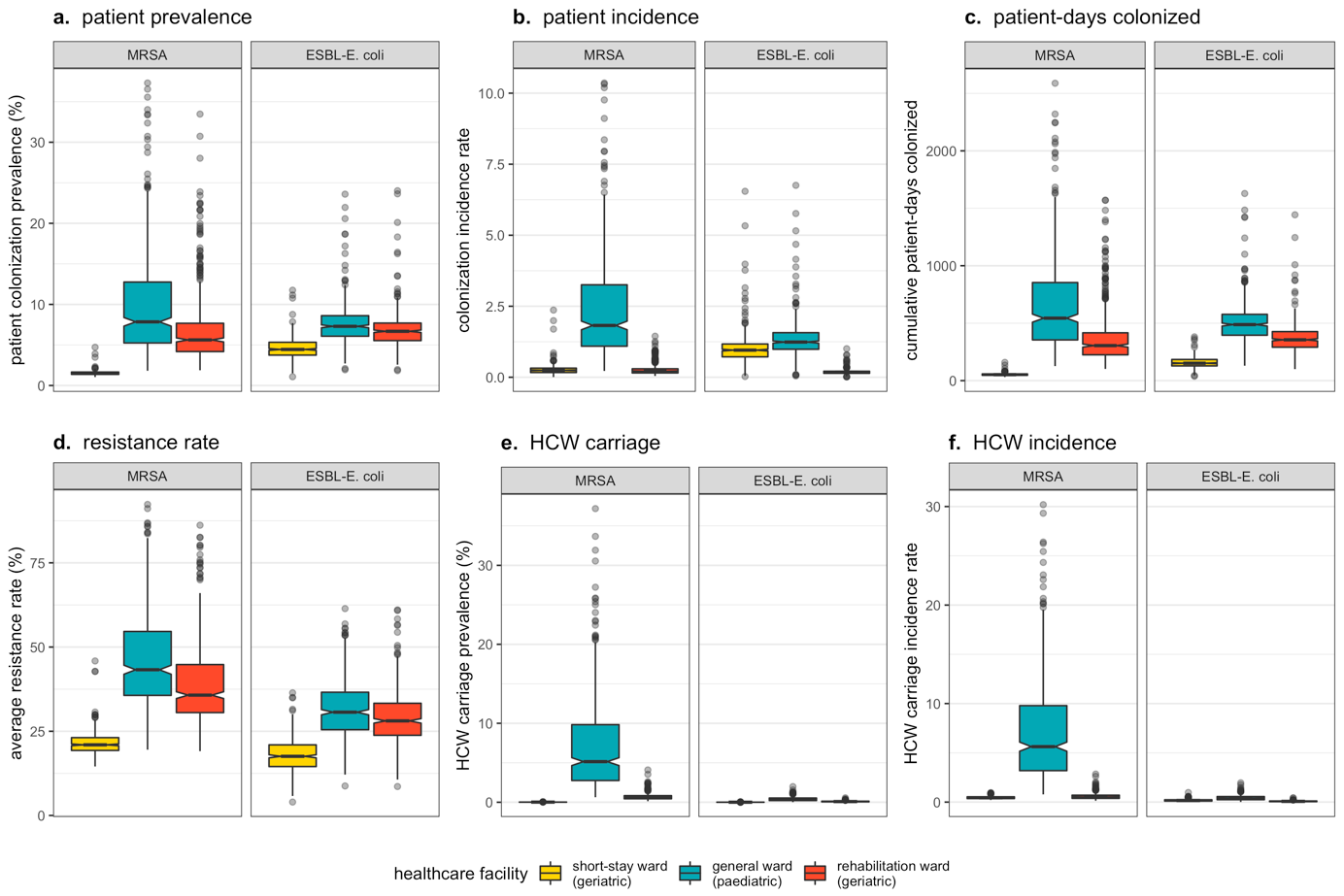

**Figure S16.** Baseline epidemiological dynamics of MRSA and ESBL-*E. coli* at endemic equilibrium in each ward (colours). Indicators represent results from simulations excluding COVID-19 responses (all $\tau=0$). Each data point represents one of $n=500$ Monte Carlo simulations. Boxplots represent the $IQR$; whiskers extend to the furthest value up to $\pm1.5\times IQR$; and notches extend $1.58\times IQR/\sqrt{n}$, giving an approximate 95% confidence interval for comparing medians. (**a**) The average daily percentage of patients colonized with each bacteria. (**b**) The rate of patient colonization acquisition per patient-day. (**c**) The cumulative number of patient-days colonized over a simulated 180-day period. (**d**) The average resistance rate. (**e**) The average daily percentage of HCWs carrying each bacteria. (**f**) The rate of HCW carriage acquisition per HCW-day.

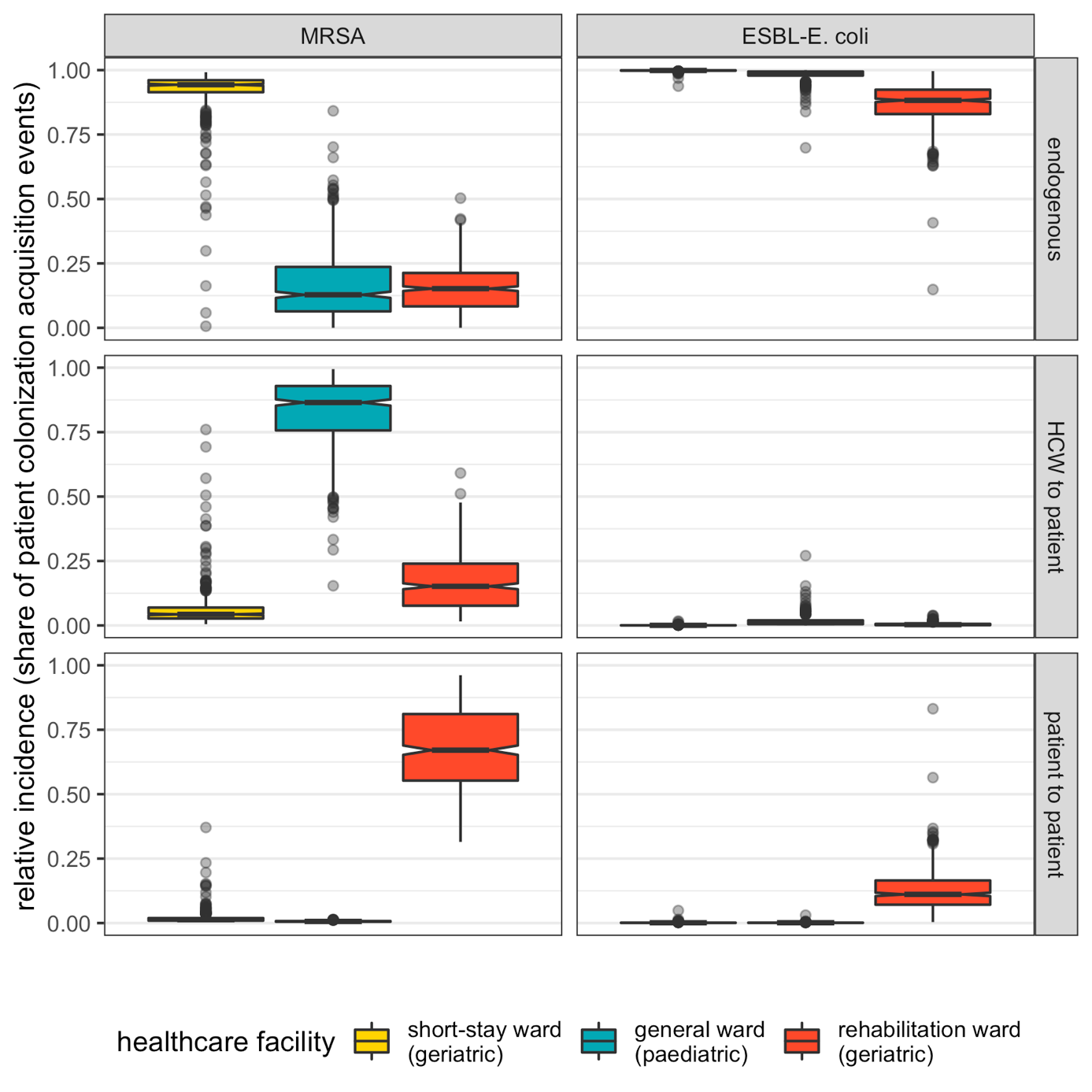

**Figure S17.** The relative share of patient colonization acquisition caused by endogenous acquisition (top row), by HCW-to-patient transmission (middle row) and by patient-to-patient transmission (bottom row) across bacteria (columns) and wards (colours) at endemic equilibrium. Each data point represents one of $n=500$ Monte Carlo simulations. Boxplots represent the $IQR$; whiskers extend to the furthest value up to $\pm1.5\times IQR$; and notches extend $1.58\times IQR/\sqrt{n}$, giving an approximate 95% confidence interval for comparing medians.

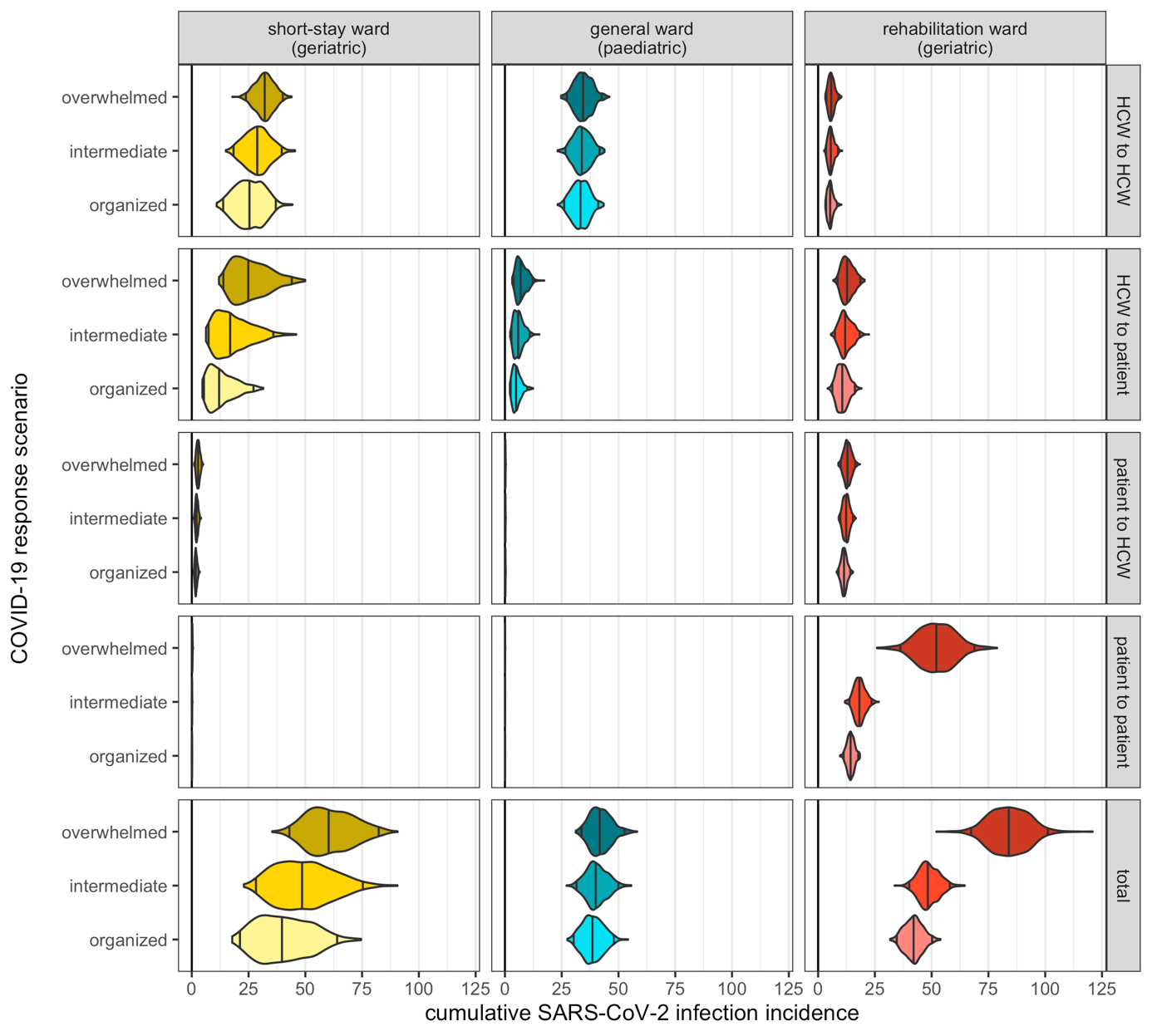

**Figure S18.** Cumulative nosocomial incidence of SARS-CoV-2 infection (x-axis) across COVID-19 response scenarios (y-axis) and hospital wards (columns). Incidence is stratified by route of infection, such that the bottom row is the sum total of the top four rows. Violin plots represent outcome distributions from $n=500$ Monte Carlo simulations. Indicators are calculated cumulatively over $t=180$ days of simulation, after introduction of two index cases of SARS-CoV-2 into the hospital at $t=0$. Here, baseline values of SARS-CoV-2 transmissibility ($\beta_{V}=1.28$) and policy implementation timing ($t_{policy}=21$) are assumed.

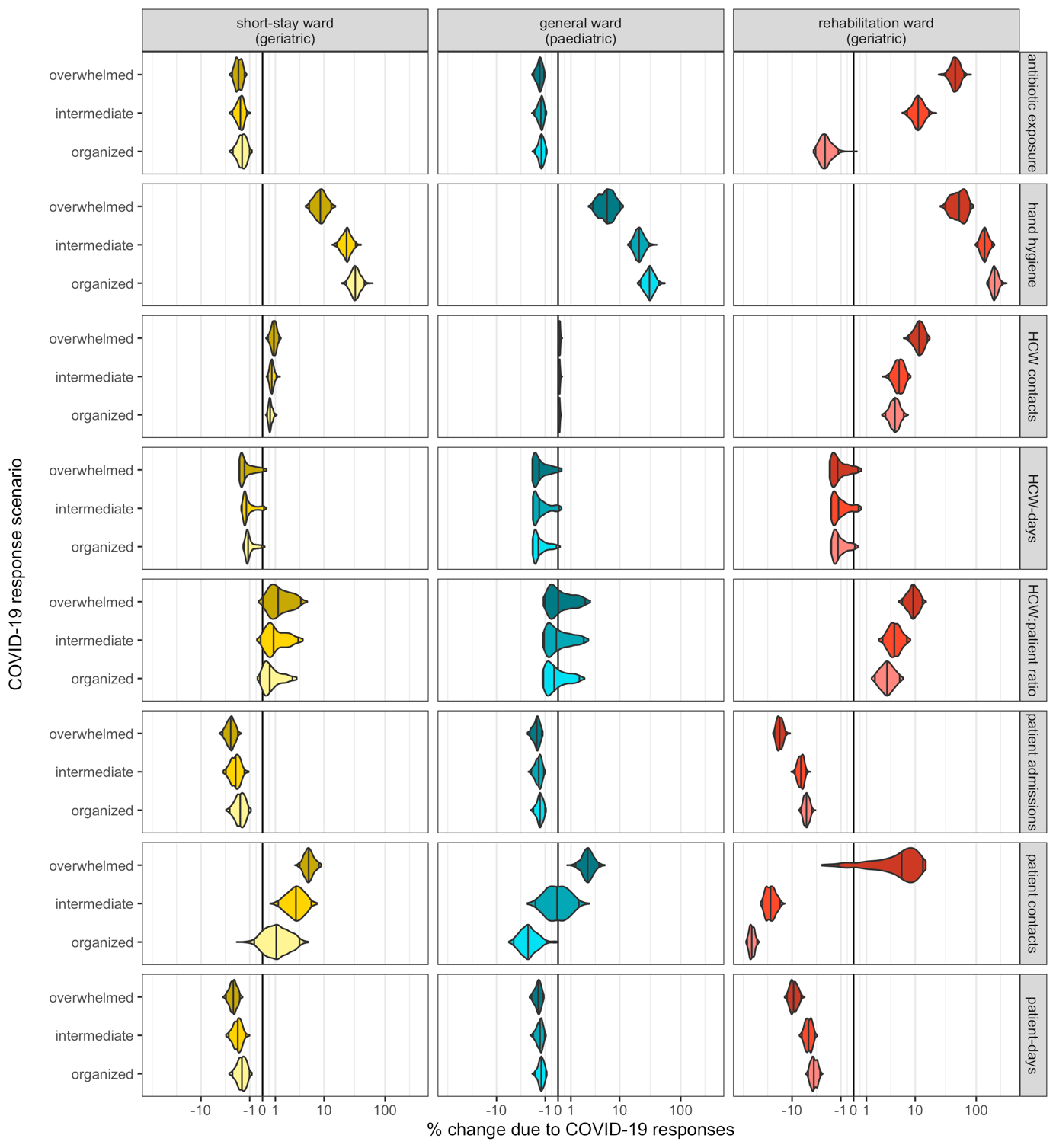

**Figure S19.** Violin plots representing outcome distributions from $n=500$ Monte Carlo simulations, and depicting change in behavioural and demographic indicators as a result of nosocomial SARS-CoV-2 outbreaks (x-axis) across different COVID-19 response scenarios (y-axis). For each ward, bacterial species and COVID-19 response scenario, change due to COVID-19 responses is calculated as the difference between matched simulations, i.e. those including the respective COVID-19 responses (organized, intermediate or overwhelmed) versus those including no COVID-19 responses ($\tau=0$). Indicators are calculated cumulatively over $t=180$ days of simulation, after introduction of two index cases of SARS-CoV-2 into the hospital at $t=0$. Here, baseline values of SARS-CoV-2 transmissibility ($\beta_{V}=1.28$) and policy implementation timing ($t_{policy}=21$) are assumed. Scales are pseudo-log_10_-transformed using an inverse hyperbolic sine function (R package ggallin).

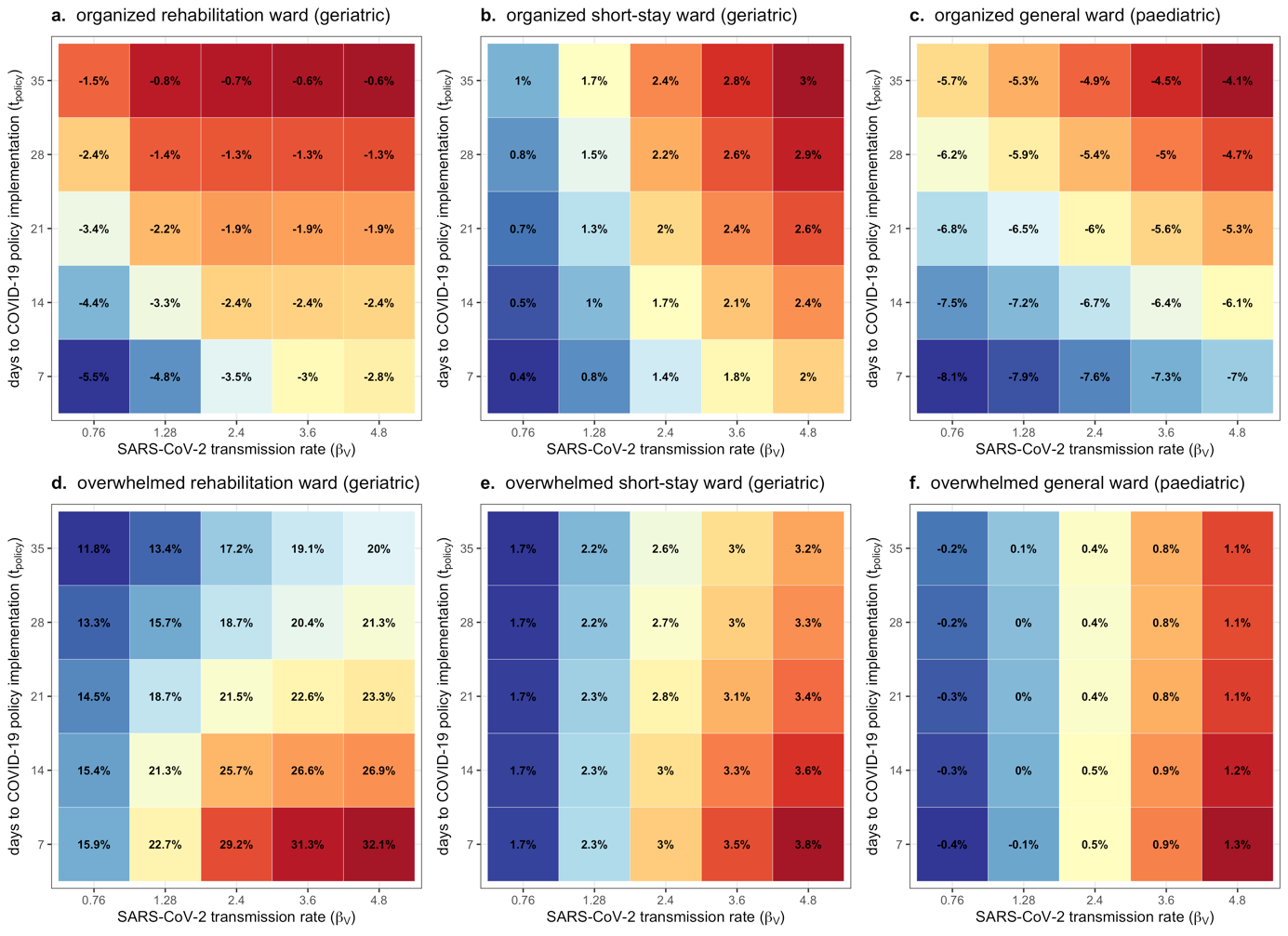

**Figure S20.** Heat map depicting change in the average resistance rate of MRSA, which varies with the SARS-CoV-2 transmission rate (x-axis) and the delay to COVID-19 policy implementation (y-axis). Each coloured tile represents the mean across $n=500$ Monte Carlo simulations. Each panel depicts results for a different hospital ward and COVID-19 response scenario (see plot subtitles). Change due to COVID-19 responses is calculated as the difference between matched simulations, i.e. those including respective COVID-19 responses (organized, overwhelmed; see **Table S5**) versus those including no COVID-19 responses ($\tau=0$). Average resistance rate is calculated cumulatively over $t=180$ days of simulation, after introduction of two index cases of SARS-CoV-2 into the hospital at $t=0$.

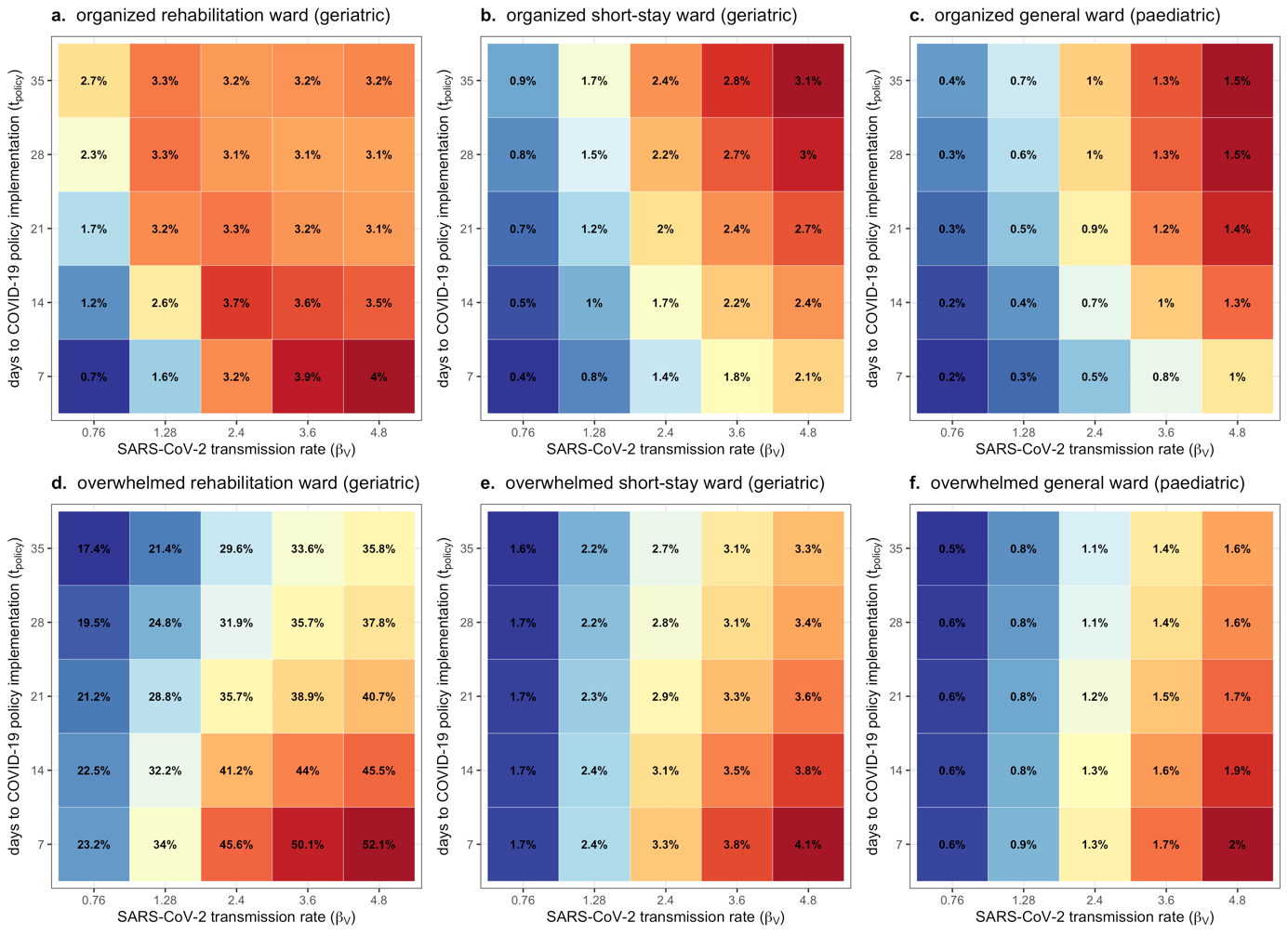

**Figure S21.** Heat map depicting change in the average resistance rate of ESBL-*E. coli*, which varies with the SARS-CoV-2 transmission rate (x-axis) and the delay to COVID-19 policy implementation (y-axis). Each coloured tile represents the mean across $n=500$ Monte Carlo simulations. Each panel depicts results for a different hospital ward and COVID-19 response scenario (see plot subtitles). Change due to COVID-19 responses is calculated as the difference between matched simulations, i.e. those including respective COVID-19 responses (organized, overwhelmed; see **Table S5**) versus those including no COVID-19 responses ($\tau=0$). Average resistance rate is calculated cumulatively over $t=180$ days of simulation, after introduction of two index cases of SARS-CoV-2 into the hospital at $t=0$.

[Bibliography](https://sciwheel.com/work/bibliography)

[1.    J. W. Tang, *et al.*, Dismantling myths on the airborne transmission of severe acute respiratory syndrome coronavirus-2 (SARS-CoV-2). *J. Hosp. Infect.* **110**, 89–96 (2021).](https://sciwheel.com/work/bibliography/10296329)

[2.    M. U. Mondelli, M. Colaneri, E. M. Seminari, F. Baldanti, R. Bruno, Low risk of SARS-CoV-2 transmission by fomites in real-life conditions. *Lancet Infect. Dis.* **21**, e112 (2021).](https://sciwheel.com/work/bibliography/9752188)

[3.    I. H. Spicknall, B. Foxman, C. F. Marrs, J. N. S. Eisenberg, A modeling framework for the evolution and spread of antibiotic resistance: literature review and model categorization. *Am. J. Epidemiol.* **178**, 508–520 (2013).](https://sciwheel.com/work/bibliography/1795601)

[4.    A. H. Melnyk, A. Wong, R. Kassen, The fitness costs of antibiotic resistance mutations. *Evol. Appl.* **8**, 273–283 (2015).](https://sciwheel.com/work/bibliography/1591706)

[5.    C. Tedijanto, S. W. Olesen, Y. H. Grad, M. Lipsitch, Estimating the proportion of bystander selection for antibiotic resistance among potentially pathogenic bacterial flora. *Proc Natl Acad Sci USA* **115**, E11988–E11995 (2018).](https://sciwheel.com/work/bibliography/6323434)

[6.    E. van Kleef, N. Luangasanatip, M. J. Bonten, B. S. Cooper, Why sensitive bacteria are resistant to hospital infection control. [version 2; peer review: 2 approved]. *Wellcome Open Res.* **2**, 16 (2017).](https://sciwheel.com/work/bibliography/9162958)

[7.    M. C. J. Bootsma, M. J. M. Bonten, S. Nijssen, A. C. Fluit, O. Diekmann, An algorithm to estimate the importance of bacterial acquisition routes in hospital settings. *Am. J. Epidemiol.* **166**, 841–851 (2007).](https://sciwheel.com/work/bibliography/10533358)

[8.    B. Stecher, L. Maier, W.-D. Hardt, “Blooming” in the gut: how dysbiosis might contribute to pathogen evolution. *Nat. Rev. Microbiol.* **11**, 277–284 (2013).](https://sciwheel.com/work/bibliography/926401)

[9.    A. Duval, *et al.*, Measuring dynamic social contacts in a rehabilitation hospital: effect of wards, patient and staff characteristics. *Sci. Rep.* **8**, 1686 (2018).](https://sciwheel.com/work/bibliography/8684589)

[10.   P. Vanhems, *et al.*, Estimating potential infection transmission routes in hospital wards using wearable proximity sensors. *PLoS ONE* **8**, e73970 (2013).](https://sciwheel.com/work/bibliography/732780)

[11.   S. M. Bartsch, *et al.*, Maintaining face mask use before and after achieving different COVID-19 vaccination coverage levels: a modelling study. *Lancet Public Health* **7**, e356–e365 (2022).](https://sciwheel.com/work/bibliography/12629215)

[12.   G. Shirreff, *et al.*, Measuring Basic Reproduction Number to Assess Effects of Nonpharmaceutical Interventions on Nosocomial SARS-CoV-2 Transmission. *Emerging Infect. Dis.* **28** (2022).](https://sciwheel.com/work/bibliography/13046772)

[13.   Q. Li, *et al.*, Early Transmission Dynamics in Wuhan, China, of Novel Coronavirus-Infected Pneumonia. *N. Engl. J. Med.* **382**, 1199–1207 (2020).](https://sciwheel.com/work/bibliography/8159772)

[14.   S. A. Lauer, *et al.*, The Incubation Period of Coronavirus Disease 2019 (COVID-19) From Publicly Reported Confirmed Cases: Estimation and Application. *Ann. Intern. Med.* **172**, 577–582 (2020).](https://sciwheel.com/work/bibliography/8378827)

[15.   X. He, *et al.*, Temporal dynamics in viral shedding and transmissibility of COVID-19. *Nat. Med.* **26**, 672–675 (2020).](https://sciwheel.com/work/bibliography/8690687)

[16.   Q. Ma, *et al.*, Global Percentage of Asymptomatic SARS-CoV-2 Infections Among the Tested Population and Individuals With Confirmed COVID-19 Diagnosis: A Systematic Review and Meta-analysis. *JAMA Netw. Open* **4**, e2137257 (2021).](https://sciwheel.com/work/bibliography/12855219)

[17.   G. M. Knight, *et al.*, Quantifying where human acquisition of antibiotic resistance occurs: a mathematical modelling study. *BMC Med.* **16**, 137 (2018).](https://sciwheel.com/work/bibliography/6089731)

[18.   D. R. Smith, L. Temime, L. Opatowski, Microbiome-pathogen interactions drive epidemiological dynamics of antibiotic resistance: A modeling study applied to nosocomial pathogen control. *eLife* **10** (2021).](https://sciwheel.com/work/bibliography/11770059)

[19.   R. D. Kouyos, P. Abel Zur Wiesch, S. Bonhoeffer, Informed switching strongly decreases the prevalence of antibiotic resistance in hospital wards. *PLoS Comput. Biol.* **7**, e1001094 (2011).](https://sciwheel.com/work/bibliography/5333453)

[20.   U. Obolski, L. Hadany, Implications of stress-induced genetic variation for minimizing multidrug resistance in bacteria. *BMC Med.* **10**, 89 (2012).](https://sciwheel.com/work/bibliography/1795566)

[21.   U. Obolski, G. Y. Stein, L. Hadany, Antibiotic Restriction Might Facilitate the Emergence of Multi-drug Resistance. *PLoS Comput. Biol.* **11**, e1004340 (2015).](https://sciwheel.com/work/bibliography/12405002)

[22.   P. Abel zur Wiesch, R. Kouyos, S. Abel, W. Viechtbauer, S. Bonhoeffer, Cycling empirical antibiotic therapy in hospitals: meta-analysis and models. *PLoS Pathog.* **10**, e1004225 (2014).](https://sciwheel.com/work/bibliography/2894755)

[23.   B. Tepekule, H. Uecker, I. Derungs, A. Frenoy, S. Bonhoeffer, Modeling antibiotic treatment in hospitals: A systematic approach shows benefits of combination therapy over cycling, mixing, and mono-drug therapies. *PLoS Comput. Biol.* **13**, e1005745 (2017).](https://sciwheel.com/work/bibliography/6258975)

[24.   B. Cooper, *et al.*, The burden and dynamics of hospital-acquired SARS-CoV-2 in England. *Res. Sq.* (2021) https:/doi.org/10.21203/rs.3.rs-1098214/v1.](https://sciwheel.com/work/bibliography/12143649)

[25.   D. R. MacFadden, D. N. Fisman, W. P. Hanage, M. Lipsitch, The Relative Impact of Community and Hospital Antibiotic Use on the Selection of Extended-spectrum Beta-lactamase-producing Escherichia coli. *Clin. Infect. Dis.* **69**, 182–188 (2019).](https://sciwheel.com/work/bibliography/6260479)

[26.   L. Isella, *et al.*, Close encounters in a pediatric ward: measuring face-to-face proximity and mixing patterns with wearable sensors. *PLoS ONE* **6**, e17144 (2011).](https://sciwheel.com/work/bibliography/732383)

[27.   R. Assab, L. Temime, The role of hand hygiene in controlling norovirus spread in nursing homes. *BMC Infect. Dis.* **16**, 395 (2016).](https://sciwheel.com/work/bibliography/9092216)

[28.   F. Di Ruscio, *et al.*, Quantifying the transmission dynamics of MRSA in the community and healthcare settings in a low-prevalence country. *Proc Natl Acad Sci USA* **116**, 14599–14605 (2019).](https://sciwheel.com/work/bibliography/7377956)

[29.   T. Gurieva, *et al.*, The Transmissibility of Antibiotic-Resistant Enterobacteriaceae in Intensive Care Units. *Clin. Infect. Dis.* **66**, 489–493 (2018).](https://sciwheel.com/work/bibliography/7174876)

[30.   R. Coello, J. R. Glynn, C. Gaspar, J. J. Picazo, J. Fereres, Risk factors for developing clinical infection with methicillin-resistant Staphylococcus aureus (MRSA) amongst hospital patients initially only colonized with MRSA. *J. Hosp. Infect.* **37**, 39–46 (1997).](https://sciwheel.com/work/bibliography/5496695)

[31.   E. S. Shenoy, M. L. Paras, F. Noubary, R. P. Walensky, D. C. Hooper, Natural history of colonization with methicillin-resistant Staphylococcus aureus (MRSA) and vancomycin-resistant Enterococcus (VRE): a systematic review. *BMC Infect. Dis.* **14**, 177 (2014).](https://sciwheel.com/work/bibliography/4942267)

[32.   H. Bar-Yoseph, K. Hussein, E. Braun, M. Paul, Natural history and decolonization strategies for ESBL/carbapenem-resistant Enterobacteriaceae carriage: systematic review and meta-analysis. *J. Antimicrob. Chemother.* **71**, 2729–2739 (2016).](https://sciwheel.com/work/bibliography/2914123)

[33.   R. Kouyos, E. Klein, B. Grenfell, Hospital-community interactions foster coexistence between methicillin-resistant strains of Staphylococcus aureus. *PLoS Pathog.* **9**, e1003134 (2013).](https://sciwheel.com/work/bibliography/2894934)

[34.   F. Laurent, *et al.*, Fitness and competitive growth advantage of new gentamicin-susceptible MRSA clones spreading in French hospitals. *J. Antimicrob. Chemother.* **47**, 277–283 (2001).](https://sciwheel.com/work/bibliography/2679877)

[35.   A. Ranjan, *et al.*, ESBL-plasmid carriage in E. coli enhances in vitro bacterial competition fitness and serum resistance in some strains of pandemic sequence types without overall fitness cost. *Gut Pathog.* **10**, 24 (2018).](https://sciwheel.com/work/bibliography/6683421)

[36.   T. Cravo Oliveira Hashiguchi, D. Ait Ouakrim, M. Padget, A. Cassini, M. Cecchini, Resistance proportions for eight priority antibiotic-bacterium combinations in OECD, EU/EEA and G20 countries 2000 to 2030: a modelling study. *Euro Surveill.* **24** (2019).](https://sciwheel.com/work/bibliography/7643014)

[37.   A. Scanvic, *et al.*, Duration of colonization by methicillin-resistant Staphylococcus aureus after hospital discharge and risk factors for prolonged carriage. *Clin. Infect. Dis.* **32**, 1393–1398 (2001).](https://sciwheel.com/work/bibliography/8810222)

[38.   F. Ebrahimi, *et al.*, Comparison of rates of fecal colonization with extended-spectrum beta-lactamase-producing enterobacteria among patients in different wards, outpatients and medical students. *Microbiol. Immunol.* **60**, 285–294 (2016).](https://sciwheel.com/work/bibliography/6250840)

[39.   G. R. Teesing, *et al.*, Increased hand hygiene compliance in nursing homes after a multimodal intervention: A cluster randomized controlled trial (HANDSOME). *Infect. Control Hosp. Epidemiol.* **41**, 1169–1177 (2020).](https://sciwheel.com/work/bibliography/9887763)

[40.   L. D. Moore, G. Robbins, J. Quinn, J. W. Arbogast, The impact of COVID-19 pandemic on hand hygiene performance in hospitals. *Am. J. Infect. Control* **49**, 30–33 (2021).](https://sciwheel.com/work/bibliography/9529501)

[41.   E. Ricchizzi, *et al.*, Antimicrobial use in European long-term care facilities: results from the third point prevalence survey of healthcare-associated infections and antimicrobial use, 2016 to 2017. *Euro Surveill.* **23** (2018).](https://sciwheel.com/work/bibliography/12683639)

[42.   D. Plachouras, *et al.*, Antimicrobial use in European acute care hospitals: results from the second point prevalence survey (PPS) of healthcare-associated infections and antimicrobial use, 2016 to 2017. *Euro Surveill.* **23** (2018).](https://sciwheel.com/work/bibliography/11600757)

[43.   A. Versporten, *et al.*, The Worldwide Antibiotic Resistance and Prescribing in European Children (ARPEC) point prevalence survey: developing hospital-quality indicators of antibiotic prescribing for children. *J. Antimicrob. Chemother.* **71**, 1106–1117 (2016).](https://sciwheel.com/work/bibliography/7134488)

[44.   B. J. Langford, *et al.*, Antibiotic prescribing in patients with COVID-19: rapid review and meta-analysis. *Clinical Microbiology and Infection* (2021).](https://sciwheel.com/work/bibliography/10265615)

[45.   W. G. Lindsley, F. M. Blachere, B. F. Law, D. H. Beezhold, J. D. Noti, Efficacy of face masks, neck gaiters and face shields for reducing the expulsion of simulated cough-generated aerosols. *Aerosol Science and Technology*, 1–12 (2020).](https://sciwheel.com/work/bibliography/11275956)

[46.   M. Stevenson, *et al.*, epiR: Tools for the analysis of epidemiological data (2018) (April 10, 2022).](https://sciwheel.com/work/bibliography/13307238)
